# Multiplexed, scalable analog recording of gene regulation dynamics over weeks using intracellular protein tapes

**DOI:** 10.1101/2025.05.10.653182

**Authors:** Lirong Zheng, Yixiao Yan, Bingxin Zhou, Jormay Lim, Dongqing Shi, Bobae An, BumJin Ko, Emily Klyder, Sethuramasundaram Pitchiaya, Denise J. Cai, Edward S. Boyden, Donglai Wei, Pietro Liò, Changyang Linghu

## Abstract

Gene expression is constantly regulated by gene regulatory networks that consist of multiple regulatory components to mediate cellular functions. An ideal tool for analyzing gene regulation processes would provide simultaneous measurements of the dynamics of many components in the gene regulatory network, but existing methodologies fall short of simultaneously tracking the dynamics of components over long periods of time. Here, we present CytoTape—a genetically encoded, modular, and scalable analog recorder for continuous, multiplexed *in situ* recording of gene regulation dynamics over multiple days and weeks at single-cell resolution. CytoTape consists of a flexible, thread-like, elongating intracellular protein self-assembly engineered via AI-guided rational design. Gene regulation dynamics, together with timestamps for reconstruction of the continuous time axis, are directly encoded via distinct molecular tags distributed along single CytoTape assemblies in live cells, to be readout at scale after fixation via standard immunofluorescence imaging. CytoTape recorders are modularly designed to record gene expression driven by a variety of activity-dependent promoters. We demonstrated the utility of CytoTape in mammalian embryonic kidney cells, cancer cells, glial cells, and neurons, achieving simultaneous recording of five cell plasticity-associated transcription factor activities and immediate early gene expression levels, namely CREB, c*-fos*, *Arc*, *Egr1*, and *Npas4* activities, within single cells in a spatiotemporally scalable manner. CytoTape revealed complex waveforms and nonlinear temporal couplings among these cellular activities, enabling investigations of how gene regulation histories and intrinsic signaling states shape transcriptional logics. We envision CytoTape to have broad applications in both basic and disease-related cell biology research.

## 1 Introduction

Gene regulatory networks orchestrate intricate biological processes within cells, determining cellular fates, coordinating environmental responses, and when disrupted, contributing to disease states^1–7^. These networks consist of a collection of dynamic components (*e.g.*, transcription factors) that interact with each other to achieve network dynamics that ultimately drive complex cellular computations and outcomes. However, such dynamic interactions can barely be adequately captured via static snapshots or time-resolved measurement of the activity of a single network component. Even if one could perform time-resolved measurement of each component separately in distinct cell populations, critical relationships among the components within single cells would be lost. For example, even a straightforward inverse relationship—A is high when B is low, and vice versa—cannot be established when A and B are measured in two separate cell populations. Recent studies suggest gene regulatory networks exhibit far more complex dynamical rules of action, including intricate activation-inhibition patterns and temporal sequences^8, 9^. Dissecting the fundamental principles of gene regulatory networks in action would greatly benefit from a technology that provides simultaneous, time-resolved recording of the temporal dynamics of multiple interacting network components within the same cell, capturing both their activation and inhibition patterns over time.

Despite growing interest in decoding the dynamics of gene regulatory networks, existing technologies fall short of meeting the combined demands of days- to weeks-long, single-cell multiplexed, and scalable recording of transcriptional activities in live cells. Genetically encoded fluorescent reporters allow real-time imaging of cellular activities in live cells^10–12^ and can be multiplexed via spectral multiplexing^13–15^, spatial multiplexing^16^, and temporal multiplexing^17^. However, long-term live imaging of these reporters typically requires continuous microscopy access, precise environmental control for long-term imaging, and repeated manual interventions, with restricted spatial and temporal scales due to limited field of view sizes of imaging systems and photobleaching effects of fluorescent probes over days and weeks. Nucleic acid-based molecular recorders have expanded our ability to capture and reconstruct intracellular events across large cell populations by encoding molecular information stably in nucleic acid sequences for downstream high-throughput sequencing readout^18–35^. Nevertheless, these methods do not yet support multiplexed recording of the complex temporal waveforms of cellular signals within single mammalian cells and typically require cell lysis and/or tissue dissociation that does not preserve the spatial and molecular information of intact cells and tissues that one often wants to examine *in situ*. Post-fixation readout methods support multiplexed spatial analysis of intact cells and tissues, but are generally limited to one or two time points^36, 37^. To study cellular processes beyond a few hours, there is an urgent need for a technology that provides long-term, multiplexed measurements of cellular activities. The recently proposed molecular recorder concept via intracellular linear protein assemblies^38, 39^ allows cellular measurements over days across large cell populations, because cellular activity is stored *in situ* for post-fixation scalable readout (by encoding temporal signals along the spatial dimension; analogous to how tree rings encode physiological histories of a growing tree), thus eliminating the need for live-cell imaging. However, current protein-based “ticker tape” recorders do not support simultaneous recording of multiple cellular signals, nor do they allow weeks-long recording due to the rigid structure of the protein assembly being restricted by the size of a cell.

In this work, we addressed this enduring technological gap through AI-guided protein engineering by designing CytoTape—a temporally precise and flexible protein tape system that achieves single-cell multiplexed, weeks-long analog recording in live cells in a highly modular manner (Fig. 1a; the protein design of the CytoTape monomer is shown in Fig. 1b). We discovered that multiplexed recording can be achieved on the same protein assembly by using distinct molecular tags where each encodes one kind of cellular signal. We also found that stabilizing the molecular dynamics of monomers of the assembly and increasing the electrostatic interaction level of the lateral binding surfaces of the monomer could encourage the assembly to be thin and flexible, allowing recording over weeks. Using CytoTape, we achieved simultaneous, spatiotemporally scalable recording of five cell plasticity-associated transcription factor activities and immediate early gene (IEG) transcriptional activities, namely cAMP response element-binding protein (CREB), c-*fos*, *Arc*, *Egr1*, and Npas4 activities, within the same protein assemblies in individual live cells (throughout this paper, transcription factor activities measured using transcription factor-responsive promoters are written in non-italic font, whereas gene transcriptional activities measured via their respective gene promoters are *italicized* to distinguish between the two). This capability surpasses existing state-of-the-art methodologies by providing in both temporal and multiplexed scalability and offering a powerful new platform for dissecting complex cellular dynamics. To demonstrate the utility of CytoTape, we applied it to analyze the long-term temporal coupling between CREB activity and downstream IEG expression at single-cell resolution—an experiment otherwise infeasible with existing tools. Recent studies suggest that dynamic transcriptional signatures play important roles in regulating CREB–Fos regulation, but the underlying temporal logic remains unclear^40–42^. Using CytoTape, we observed that CREB activation and Fos expression can become decoupled in cells exhibiting active Fos transcriptional activity prior to stimulation, suggesting a potential role for transcriptional history in shaping downstream responses^43–45^. In primary cultured neurons, we also observed that *Arc*- and *Egr1*-promoter-driven gene expression display complex, multi-peak activation dynamics following a single upstream stimulation, revealing distinct temporal couplings dependent on stimulus type. Together, these results highlight how temporal features of gene activity and signaling history contribute to transcriptional outcomes, offering a framework to dissect gene regulation dynamics in diverse cellular contexts.

**Fig. 1.**
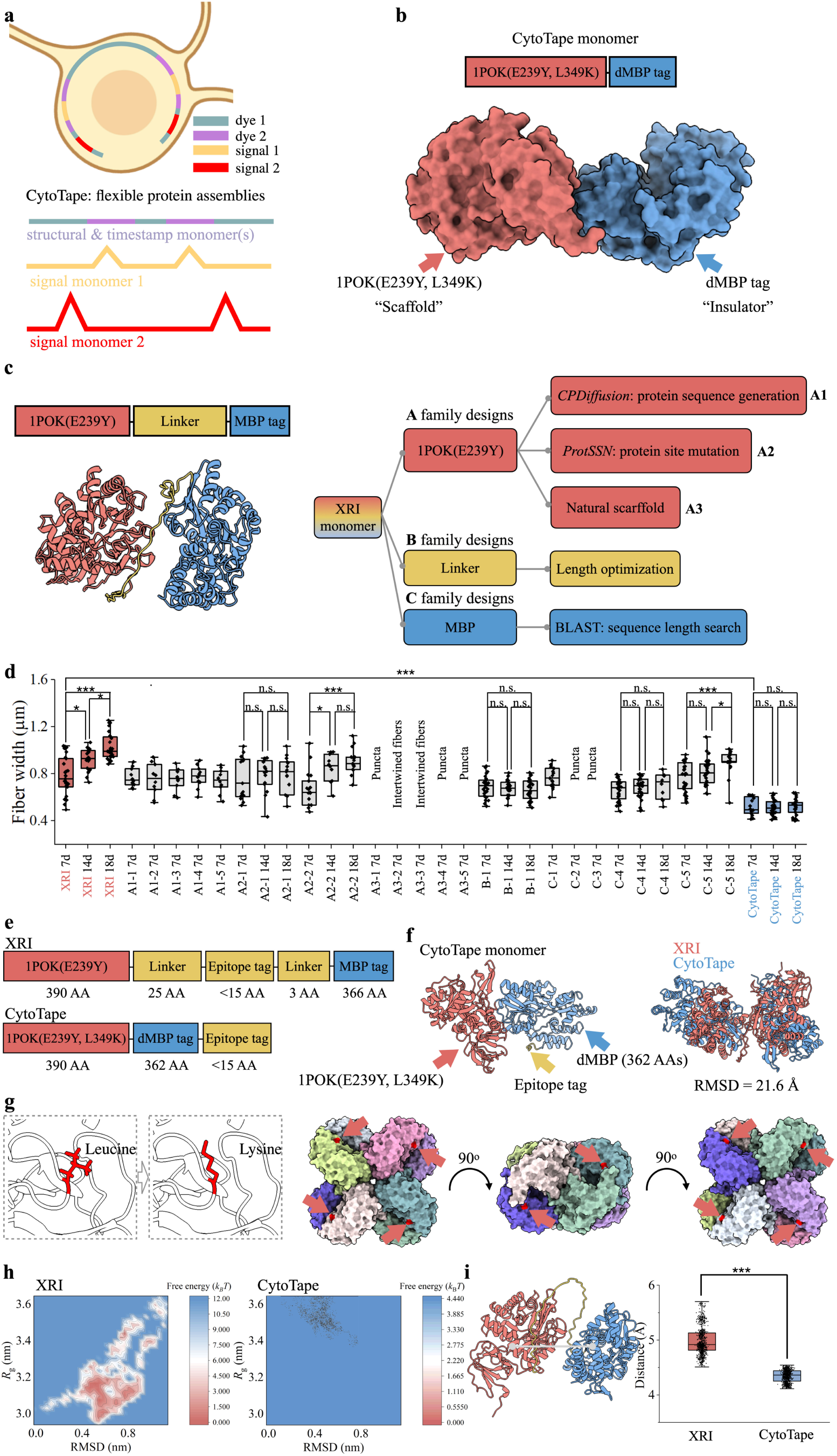
Design of CytoTape. (**a**) Schematic of the CytoTape system for long-term, multiplexed recording of cellular physiological histories via elongating protein self-assembly. CytoTape is formed by structural monomers that create the intracellular protein assembly. Timestamp monomers, which encode temporal information, are sparsely incorporated along this assembly to embed the time axis along the spatial dimension. Signal monomers encode cellular physiological activities and are also incorporated along the same assembly to achieve recording. (**b**) Schematic (top row) and AlphaFold3-predicted structure (bottom row) of the CytoTape monomer engineered via the design strategy described in **c**. (**c**) Design strategy of CytoTape using XRI as a template. Schematic (left panel, top row) and AlphaFold3-predicted structure (left panel, bottom row) of the XRI monomer. Right panel, AI-assisted protein design (**A** family designs; *CPDiffusion* and *ProtSSN* are used for sequence generation and site mutation, respectively, using 1POK(E239Y) as the template) and rational design (**B** and **C** family designs; 1POK(E239Y) was replaced with other assembly proteins, and the linker and MBP were optimized by tuning their sequence lengths, adjusting the number of their tandem repeats, and replacing them with homologous sequences) were combined to develop the CytoTape monomer. After *in silico* design, design candidates were screened in cultured neurons. The *UbC* promoter was used to drive the steady expression of candidate protein monomers. (**d**) Statistical analysis of protein assembly width for different protein designs that showed fiber-like structures in cultured neurons at 7 days, 14 days, and 18 days after transfection. The sample size (n) is listed below as (X, Y, Z), which dentotes n = X protein assemblies from Y neurons from Z cultures. XRI: 7d (31, 25, 5), 14d (26, 20, 5), 18d (32, 30, 5). A1-1: 7d (10, 5, 1). A1-2: 7d (10, 6, 1). A1-3: 7d (10, 7, 1). A1-4: 7d (10, 6, 1). A1-5: 7d (10, 7, 1). A2-1: 7d (15, 10, 2), 14d (15, 15, 2), 18d (15, 12, 3). A2-2: 7d (17, 10, 2), 14d (10, 10, 2), 18d (15, 15, 2). B-1: 7d (31, 28, 4), 14d (25, 20, 6), 18d (25, 25, 3). C-1: 7d (15, 12, 3). C-4: 7d (28, 20, 3), 14d (30, 30, 4), 18d (10, 10, 2). C-5: 7d (31, 30, 2), 14d (24, 20, 2), 18d (18, 15, 3). CytoTape: (16, 16, 3), 14d (28, 20, 5), 18d (25, 20, 5). The plasmid constructs of all protein assembly designs are listed in **Supplementary information**. The details of protein assembly width quantification are described in **Methods**. For each design group (7, 14, and 18 days), Kruskal–Wallis analysis of variance followed by Dunn’s post hoc tests was used; *, *P <* 0.05; **, *P <* 0.01; ***, *P <* 0.001; n.s., not significant. To compare the width of XRI and CytoTape, Mann–Whitney U test was used; ***, *P <* 0.001. Middle line in box plot, median; box boundary, interquartile range; whiskers, minimum and maximum; black dots, individual data points. (**e**) Schematic constructs of XRI (top row) and CytoTape (bottom row). AA, amino acid. (**f**) AlphaFold3-predicted structure of the CytoTape monomer with an epitope tag fused at the C-terminus (left panel) and the superposition of XRI monomer structure (red) and CytoTape monomer structure (blue) (right panel). The alignment on *C_α_* is conducted by the PyMOL software. (**g**) Position of the L349K mutation (highlighted in red; also indicated by red arrows) in the octamer structure (obtained from Protein Data Bank, ID: 1POK) of the original *E. coli* IadA protein. Each colored domain in the octamer structure represents an individual, identical monomer. The E239Y mutation enables the octamers to further self-assemble longitudinally into a fiber. (**h**) Free energy landscape of a single XRI or CytoTape protein monomer calculated from MD simulations. The radius of gyration (*R*_g_) is the root mean square distance of the monomer from the center of mass of the structure. RMSD is the root mean squared deviation of the structure in the first MD snapshot after equilibration. (**i**) Distance between the centroid of the scaffold (1POK(E239Y) for XRI, 1POK(E239Y, L349K) for CytoTape) and the insulator (MBP for XRI, dMBP for CytoTape) obtained from MD simulations. The MD simulations were replicated three times. The details of MD simulation are shown in **Methods**. ***, *P <* 0.001; Mann–Whitney U test. Middle line in box plot, median; box boundary, interquartile range; whiskers, minimum and maximum; black dots, individual data points. See **Supplementary Table 2** for details of statistical analysis.

## 2 Results

### 2.1 Design, screening, and *in silico* characterization for CytoTape

We first set out to design an intracellular linear protein assembly that enables multiplexed, weeks-long recording. We reasoned that recording more kinds of information for longer durations would demand an increased information storage capacity of the linear protein assembly. Because information is encoded and stored into molecular tags along the protein assembly in protein tape recording systems^38, 39^, the storage capacity is in principle proportional to the product between the number of the tags along the elongation (longitudinal) axis of the assembly (but not along the lateral axis orthogonal to the elongation axis) and the number of possible tag variants at each tag location (*i.e.*, the bit depth of the tag). Increasing the maximum length of the assembly permitted in live cells would enhance the former (for a given line density of tags defined by the line density of tag-bearing monomers of the protein assembly), while increasing the number of tag variations would enhance the latter, and we proceeded with these two directions.

We first worked towards increasing the maximum permitted length of the protein assembly in live cells. We reasoned that if the assembly is rigid, once its length reaches the size of the cell, further elongation of the assembly will either be disrupted by the intracellular spatial constraints or distort the cell membrane that has a risk of causing unwanted physiological changes of the cell being recorded^46^. If the assembly is flexible and could curl up in a thread-like manner in cells, it may reach lengths much longer than the cell size. In addition, we concluded that minimal lateral growth of the assembly orthogonal to the elongation axis is critical to support time-resolved recording, since the XRI system showed significant lateral growth of the assembly does not preserve the temporal order of the tags along the assembly^38^. We performed protein engineering based on the XRI design because, compared to the iPAK4 design^39, 47–49^, XRI is not rigid, bioothogonal to the mammalian genome, and has a symmetrical bidirectional growth that can be conveniently used for cross-validation of recorded information from live cells. In addition, the XRI design utilizes the 1POK(E239Y) protein subunit engineered from the *E. coli* isoaspartyl dipeptidase IadA to achieve supramolecular self-assembly^50^. This assembly has been experimentally identified as a unique “agglomerate-type” arising from specific interactions among well-folded subunits, in contrast to the “aggregate-type” resulting from non-specific interactions among misfolded subunits, and has been reported to be benign in live cells without altering cellular physiology via electrophysiology, transcriptomics, proteomics, and immunohistochemistry analysis *in vitro* and *in vivo*^38, 51^. We hypothesized that if we could reduce the thickness of the XRI assembly, we would soften its mechanical rigidity and increase its flexibility. We also speculated that the thickness of the protein fiber is influenced by electrostatic interactions among amino acids (AAs) on the lateral binding surfaces and the size of protein monomer, and thus minimizing lateral binding kinetics of protein monomer may suppress lateral growth of protein assembly. This, in turn, would result in thinner assemblies that are more flexible and better suited for long-term growth and recording in live cells.

The molecular architecture of the XRI protein monomer^38, 50^ consists of three distinct domains: the self-assembling scaffold domain 1POK(E239Y)^51^, the insulator domain derived from *E. coli* maltose-binding protein (MBP), and an unstructured glycine-rich linker with an epitope tag between the two domains (Fig. 1c, left panel; the atomic structure is predicted by AlphaFold3^52^). The scaffold domain facilitates protein self-assembly, the linker connects the scaffold to the insulator, and the insulator domain reduces lateral growth of the protein assembly^38^. We first modified each of the domains individually to gain insights into how changes in each domain alter the assembly morphology in live mammalian cells, expecting that later we could combine these changes and insights to build a new protein monomer as a whole.

The overall design and screening workflow proceeded as follows (Fig. 1c and d). To begin, multiple computational and rational strategies were employed to generate and optimize protein monomers, covering the design space widely. These candidates were fused to the hemagglutinin (HA) epitope tag and then expressed in primary cultures of mouse hippocampal neurons via calcium phosphate transfection and evaluated across multiple timescales (7, 14, and 18 days after transfection) (see Table S1 for amino acid sequences of the motifs and Table S2 for all tested constructs in **Supplementary information**). Following immunofluorescence staining of the HA tag, confocal microscopy was used for imaging, and statistical analyses were performed on the measured widths of resulting protein assemblies. To benchmark our new designs, we measured the protein assembly width of the XRI design at 7, 14, and 18 days. The XRI assembly showed a fiber-like morphology that progressively increased in thickness over 18 days (Fig. 1d, highlighted in red), suggesting continuous lateral incorporation of protein monomers.

We independently optimized the scaffold (1POK(E293Y)), the linker, and the insulator (MBP) domains by using multiple design methods described below (Fig. 1c). For scaffold domain, we employed three distinct design strategies in parallel: (1) applying a sequence generation model, *CPDiffusion*^53^, to design novel scaffold AA sequences (we set the target sequence identity between generated sequences and the original scaffold sequence to be below 80% to favor novel designs and to broadly sample the sequences space) (referred to as the A1 family designs), (2) using a self-supervised protein sequence and structure representation model, *ProtSSN*^54^, to predict beneficial single-site mutations (referred to as the A2 family designs), and (3) exploring alternative naturally existing self-assembling proteins^55–59^ as potential replacements for the original scaffold domain (referred to as the A3 family designs). For the linker domain, we first relocated the HA epitope tag to the C-terminus (whereas in the XRI design, the HA epitope tag is positioned next to the linker domain between 1POK(E239Y) and MBP) and then optimized the linker by varying its length (referred to as the B family designs). For the insulator domain, we conducted homology-based sequence searches (BLAST)^60^ to identify MBP homologs with different radius of gyration (*R*_g_) and also performed truncation and duplication of the original MBP (referred to as the C family designs).

#### *CPDiffusion* for protein sequence generation

*CPDiffusion*^53^ generates novel protein sequences using a discrete diffusion probabilistic model (Extended Data Fig. 1a), capturing the joint distribution of residues across the entire sequence for a given protein backbone while integrating additional constraints, such as AA substitution frequencies and fixed conserved positions (*e.g.*, catalytic sites and binding sites). It has been deployed to facilitate the design of novel enzymes with multi-domain molecular structures through *in silico* generation and screening. Surprisingly, we found that the novel protein scaffold designs generated by *CPDiffusion* (designs A1-1 to A1-5) successfully formed fibers in cultured neurons after 7 days, with thickness comparable to that of XRI (Mann-Whitney U (MWU) tests with Bonferroni correction, *P* = 0.855, 0.585, 1.000, 0.952, 0.585 with the significance level adjusted to 0.01, for A1-1, A1-2, A1-3, A1-4, A1-5, respectively), despite having sequence identities between 70% and 80% (mutating 80 to 120 AAs simultaneously in each design) relative to the original 1POK(E293Y) scaffold (Fig. 1d, Extended Data Fig. 2). These results demonstrate the potential of diffusion models to generate novel, beyond-micron-scale protein assemblies and architectures in live mammalian cells. This capability arises from the generative model’s ability to learn sequence–structure relationships from natural proteins and to optimize protein sequences by introducing multiple AA substitutions simultaneously, while preserving protein function. However, there were many short fibers with aspect ratios below 3 presents in a large portion of the cells and we reasoned that they are suboptimal for our protein tape recording purposes due to their limited spatial continuity. Although we decided not to further pursue these five generated designs in this study, we envision that future efforts using generative AI algorithms may facilitate rapid improvement of customized intracellular protein assemblies for various applications.

#### *ProtSSN* for single site mutation

*ProtSSN*^54^ is a pre-trained deep learning model for zero-shot mutation fitness prediction (Extended Data Fig. 1b). It captures hidden local and global residue patterns using a protein language model and equivariant graph neural networks. Due to limited experimental data on design parameters of the protein sequence and structure that influence the morphology and growth kinetics of protein assemblies in live cells, *ProtSSN* provides a promising framework for exploring these design parameters *in silico*. We deployed this model to score and rank single-site saturation mutations of the 1POK(E293Y) scaffold and selected the top 2 mutations, L349K and E149I, for further experimental evaluation. We found that the L349K mutant (design A2-2) resulted in thinner protein fibers compared to the XRI fibers at 7 days of expression (Fig. 1d) (MWU test, *P* = 0.020). Lateral growth for this design was still observed between day 7 and day 14, but not between day 14 and day 18, indicating an improved suppression of long-term lateral growth compared to the XRI design. In contrast, the E149I mutant (design A2-1) did not lead to noticeable changes in fiber thickness compared to the XRI design (Fig. 1d) (MWU test, *P* = 0.751). Based on these observations, we retained the L349K mutation for subsequent design iterations. **Natural scaffold**: In nature, many proteins can form thin fibers, such as the *B. subtilis* cytoskeletal protein MreB_BS_^59^, *E. coli* cytoskeletal protein MreB_EC_^57, 58^, and cytochrome-based nanowires from prokaryote^55, 56^. To explore whether these proteins have the potential to form thin and well-structured fibers in live mammalian cells, we replaced 1POK(E239Y) with these proteins in the XRI design (designs A3-1 to A3-5) and then tested them in cultured neurons. We found these designs failed to form well-structured fibers and instead produced irregular, intertwined assemblies or puncta (Fig. 1d, Extended Data Fig. 2). We therefore decided not to further pursue these designs since their assembly morphologies are unlikely to support spatially resolved signal encoding and subsequent readout. **Linker optimization**: We next optimized the linker domain (Fig. 1d, Extended Data Fig. 2). We hypothesized that the 36-AA long linker domain between 1POK(E239Y) and MBP may allow large conformational fluctuations in the protein monomer (this is supported by the molecular dynamics (MD) simulations described below) and some of the permitted monomer conformations could potentially provide unwanted lateral binding surfaces along the fiber, resulting in lateral growth over time. Thus, we shortened the linker to 2 AAs while also moving the HA tag next to the linker to the C-terminus of the entire protein monomer (design B-1) and found that it indeed results in thinner protein fibers (MWU test, *P* = 0.0150) with reduced lateral growth over 18 days, indicating that linker shortening is a working design element. **MBP size tuning and optimization**: We also optimized the MBP domain (designs C1 to C5) (Fig. 1d, Extended Data Fig. 2). When the original 366-AA MBP domain from *E. coli* in the XRI design was replaced with a smaller, homologous 246-AA MBP from *K. pneumoniae* (*Kp*MBP) identified by BLAST (design C-1), fibers with asymmetrical thickness (one end thicker than the other end) were observed after 7 days of expression in cultured neurons (MWU test, *P* = 0.879). When replacing the original MBP with a larger, homologous 453-AA MBP from *P. stutzeri* (*Ps*MBP) identified by BLAST (design C-2) or with tandem repeats of the original 366-AA *E. coli* MBP (design C-3), only puncta but not fibers were formed. We also fused mEGFP to the XRI monomer using a self-cleavage P2A linker (design C-5), expecting the monomeric mEGFP can sparsely and stochastically coating along the fiber to block lateral monomer binding and reduce lateral growth^38^, but found this design has a fiber thickness comparable to the XRI fiber (MWU test, *P* = 0.699) that is also increasing over weeks. Knowing that shortening the unstructured linker domain decreases fiber thickness and suppresses lateral growth, we took the design B-1 above with reduced linker length and further removed the first four residues from the N-terminus of MBP (termed dMBP, 362 AAs), resulting in design C-4. Those four neighboring AAs to the linker domain were less structured than the rest of the MBP domain based on its crystal structure^61^. We hypothesized that removing these residues would further suppress lateral growth, as their flexible nature may contribute to linker-like behaviors. As expected, the resulting fibers were notably thinner than those formed by XRI (MWU test, *P* = 0.003) and exhibited no significant lateral growth over 7, 14, and 18 days. **Design Combination**: We combined the working design elements above and found that the incorporation of the L349K mutation, complete removal of the linker, moving the epitope tag to the C-terminus of the entire protein monomer, and the use of dMBP together resulted in a design that produced fibers thinner than those of XRI and all other designs while maintaining the same thickness over weeks (MWU test, *P* < 0.0001) (Fig. 1d, highlighted in blue). This optimized design is referred to as CytoTape throughout the rest of the paper (Fig. 1e).

To investigate the structural basis for the enhanced performance of CytoTape, we computationally analyzed the CytoTape design. AlphaFold3-predicted structural alignment between the XRI monomer and the CytoTape monomer revealed substantial conformational divergence (RMSD = 21.6 Å) (Fig. 1f). Analysis of the 1POK octamer structure showed that the L349K mutation is located on the lateral interface of the scaffold (Fig. 1g), replacing a hydrophobic AA (leucine) with a hydrophilic AA (lysine). This AA mutation may have reduced lateral fiber growth by decreasing hydrophobic surface interactions among protein monomers at the lateral surfaces^62^ and enhancing hydrophilic interactions between these lateral surfaces and the surrounding aqueous microenvironment (as most of the mammalian cytoplasm is water^63^). We further used MD simulations to characterize the conformational dynamics of the CytoTape monomer. The CytoTape monomer exhibited a narrower free energy landscape than the XRI monomer (Fig. 1h). Additionally, we calculated the distance between the centroids of the 1POK(E239Y, L349K) and dMBP, and found reduced conformational fluctuation in CytoTape monomer compared to XRI monomer (Fig. 1i). Both results indicate that the increased conformational stability of the CytoTape monomer may have contributed to the formation of thinner protein fibers, as limiting the protein conformational fluctuations and transitions could in principle minimize unintended binding surfaces and favor only the longitudinal binding surfaces by design, thereby suppressing lateral growth.

### 2.2 Characterization of the CytoTape assembly in cultured neurons, HEK, and HeLa cells

We characterized the intracellular structure and kinetics of CytoTape in primary cultures of mouse hippocampal neurons, HEK, and HeLa cells, while comparing them to those of XRI and iPAK4. Cultured hippocampal neurons serve as a popular *in vitro* system for studying activity-dependent gene regulation dynamics and associated signaling activities and neuroplasticity outcomes^64–67^. HEK293T cells (referred to as “HEK” throughout the paper), derived from the human embryonic kidney, are commonly used to model transcriptional regulation in proliferative epithelial-like cells, among other usages^68, 69^. HeLa cells, originating from human cervical cancer, provide a model of dysregulated gene expression and are frequently used to study cell cycle control, stress responses, and oncogenic pathways^70, 71^. Here, we used a constitutive, ubiquitous promoter, the human ubiquitin C (*UbC*) promoter, to drive the continuous expression of the CytoTape monomer (fused to the HA tag for visualization via immunofluorescence imaging; the CytoTape monomer, with or without the HA tag, driven by the *UbC* promoter is referred to as the “structural monomer” throughout the paper) and produce growing protein assembly structures in mammalian cells. In cultured mouse hippocampal neurons, we observed differences in morphology among XRI, iPAK4, and CytoTape fibers after days- and weeks-long expressions. iPAK4 fibers induced noticeable distortions in cell membrane morphology as early as 3 days after calcium phosphate transfection (Extended Data Fig. 3a), suggesting its limited capability to support multi-week intracellular recording. This may be attributed to previous observations that iPAK4 forms rigid crystalline fibers^39, 48, 49^. XRI fibers were capable of bending when being longer than the size of the soma and did not induce noticeable distortions in cell morphology, but exhibited progressive thickening after 7 days, reflecting sustained lateral growth (Fig. 2a, top row). In contrast, CytoTape fibers remain thin even after 18 days and exhibit substantial, thread-like bending without altering cell morphology, demonstrating its large potential for long-term recording (Fig. 2a, bottom row). To characterize protein assembly kinetics in cultured neurons, we quantified the widths and lengths of XRI and CytoTape fibers at multiple time points up to 18 days (Fig. 2b). Both XRI and CytoTape underwent an initial phase of lateral thickening within the first 2 days, and XRI continued to grow laterally beyond this period. In contrast, CytoTape entered a plateau phase, maintaining a stable width thereafter (Fig. 2b, left panel), indicating a self-limiting lateral growth process. Additionally, CytoTape elongated more rapidly than XRI under the same plasmid dosage in transfection, resulting in longer assemblies between day 4 to day 18 (Fig. 2b, right panel). This observation suggests that limiting lateral growth in CytoTape facilitates more efficient incorporation of protein monomers at fiber termini. We also characterized the flexibility of protein fibers by their maximum curvatures (Fig. 2c), indicating the thinner fibers of CytoTape are more flexible than the thicker ones of XRI, consistent with our initial design rationale. Additionally, we found over 80% of the CytoTape assemblies were localized in the neuronal soma, while the remaining ones were either partially extended to or fully located in the neurites (Extended Data Fig. 3f). In the soma region, more than 95% of neurons (Extended Data Fig. 3g) contained fibers rather than puncta.

**Fig. 2.**
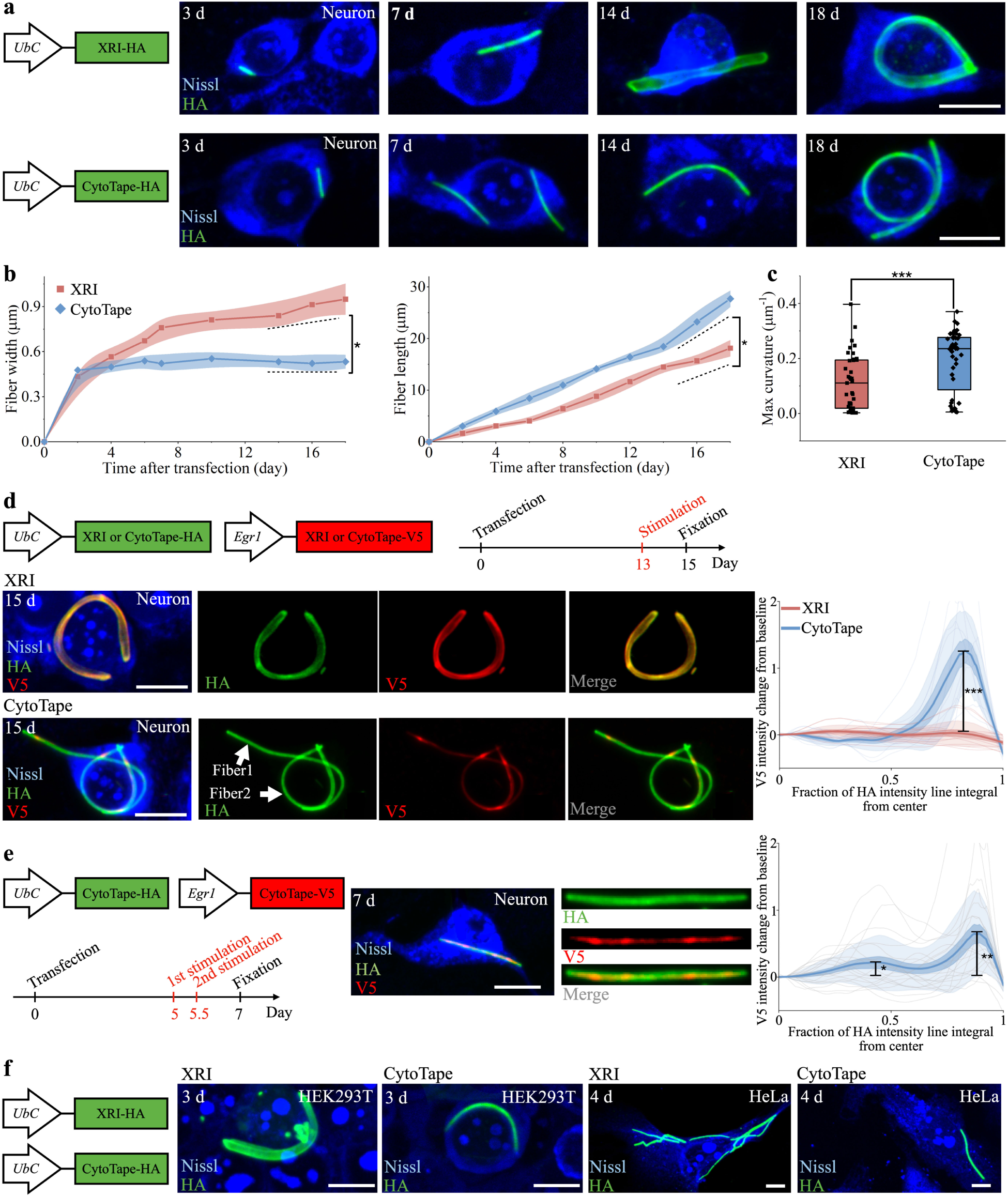
Characterization of the CytoTape assembly in live mammalian cells. (**a**) Confocal images of cultured primary mouse hippocampal neurons expressing XRI (top row) and CytoTape (bottom row) fused to the HA tag, taken after fixation at different time points (7, 14, and 18 days after calcium phosphate transfection) and then Nissl staining and immunostaining against the HA tag. Scale bars, 10 µm. (**b**) Kinetics of protein assembly width (left panel) and length (right panel) for XRI and CytoTape in cultured neurons over 18 days. Data are from n = 21 CytoTapes from 20 neurons from four cultures. The dashed lines indicate the slope of fiber width after reaching the plateau (left panel) and the slope of fiber length (right panel) between XRI and CytoTape. *, *P <* 0.05; Mann–Whitney U test. (**c**) Maximum curvature along each XRI or CytoTape assembly in cultured neurons on day 18. Data are from n = 21 CytoTapes from 20 neurons from four cultures. ***, *P <* 0.001; Mann–Whitney U test. Middle line in box plot, median; box boundary, interquartile range; whiskers, minimum and maximum; black dots, individual data points. (**d**) Left panel, first row, schematic of the constructs transfected into cultured neurons and the experimental timeline. XRI-V5 or CytoTape-V5 refers to XRI or CytoTape fused to the V5 tag as the signal monomer, which is driven by the *Egr1* (also known as *zif/268*) promoter. Confocal images of cultured mouse hippocampal neurons expressing XRI (middle row) and CytoTape (bottom row) fused to the HA tag and the V5 tag, taken after fixation on day 15, followed by Nissl staining and immunostaining against the HA and V5 tags. Neurons were stimulated with 55 mM KCl for 1 h. Right panel, V5 signal relative to baseline (calculated as the ratio of V5 signal to the baseline V5 signal at the center of the protein assembly) plotted as a function of the fraction of the HA intensity line integral. For XRI, n = 13 protein assemblies from 13 neurons from two cultures. For CytoTape, n = 11 protein assemblies from 11 neurons from two cultures. ***, *P <* 0.001; Mann–Whitney U test for the peak amplitude of the V5 signal between XRI and CytoTape. Scale bars, 10 µm. (**e**) Left panel, schematic of the constructs transfected into neuron cultures and the experimental timeline. Middle panel, confocal images of cultured mouse hippocampal neurons expressing CytoTape with the HA and V5 tags, taken after fixation on day 7, followed by Nissl staining and immunostaining against the HA and V5 tags. Neurons were stimulated twice with 55 mM KCl for 1 h. The two stimulations are 12 h apart. Right panel, V5 signal relative to baseline plotted as a function of the fraction of the HA intensity line integral. n = 20 CytoTapes from 20 neurons from two cultures. *, *P <* 0.05; **, *P <* 0.01; Mann–Whitney U test between the amplitude of each of the two sequential peaks and the baseline of the V5 signal. Scale bars, 10 µm. (**f**) Confocal images of XRI and CytoTape in HEK and HeLa cells. The constructs transfected into cells are shown at the left row. Confocal images were taken after fixation, Nissl staining, and immunostaining against the HA tag. Scale bars, 10 µm. See **Supplementary Table 2** for details of statistical analysis.

To test whether CytoTape maintains its ability to capture and store physiological processes over multi-week timescales, we co-expressed a CytoTape monomer with a V5 epitope tag under the *Egr1* promoter^72^ together with the CytoTape structural monomer in cultured neurons and 13 days after transfection and stimulated the neurons via KCl-induced depolarization (55 mM KCl, 1 h), a condition known to activate *Egr1* expression^73^ (the CytoTape monomer driven by an activity-dependent promoter is referred to as the “signal monomer” throughout the paper). For comparison, we performed the same experiment using XRI under identical conditions (Fig. 2d, top row). We fixed the neurons 15 days after transfection and performed immunofluorescence imaging, and found that under this condition XRI did not report the expected V5 signal increase along the fiber. This outcome may result from significant lateral growth of fibers over time, which could permit V5-tagged signal monomers to be incorporated along the lateral surfaces of fibers and thus the temporal order of monomers along the longitudinal axis is no longer preserved (Fig. 2d, second row). In contrast, CytoTape maintained a thin, flexible architecture and reported symmetrical V5 signal peaks along the fiber. Quantitative analysis confirmed that CytoTape was capable of recording physiological signals two weeks post-transfection in cultured neurons (Fig. 2d, right panel). We also tested whether CytoTape could resolve hours-scale temporal features of cellular activities and capture multiple cellular events over time, a desired capability for studying many cell biology processes. We performed two discrete KCl stimulations (55 mM KCl, 1 h each) separated by 12 h to the same neurons starting on day 5. We found CytoTape captured both events as distinct peaks along the fiber (Fig. 2e, left panel), indicating its ability to resolve hours-scale sequential transcriptional events over time. This result was further supported by statistical analysis (Fig. 2e, right panel).

Next, we evaluated the CytoTape performance in HEK and HeLa cells, comparing it to XRI and iPAK4. In HEK cells, XRI formed thick fibers and puncta (Fig. 2f) while iPAK4 fibers distorted cell morphology (Extended Data Fig. 3b, left panel), in agreement with previous reports^38, 39^. In contrast, CytoTape produced thin, flexible, and well-structured fibers without altering cellular morphology (Fig. 2f). Similarly, in HeLa cells, iPAK4 slightly changed cell morphology (Extended Data Fig. 3b, right panel) and XRI assemblies appeared intertwined, whereas CytoTape consistently formed uniform linear structures (Fig. 2f). Additionally, over 90% of transfected cells contained CytoTape fibers, rather than puncta, in HEK and HeLa cells (Extended Data Fig. 3h and i). CytoTape’s ability to maintain structural integrity and compatibility across cell types could be attributed to the low free-energy landscape of its monomer (Fig. 1h and i), making it less affected by perturbations from the surrounding microenvironment. We also observed that the growth rate of CytoTape fiber varies across different cell types, with the monomer expression driven by the same *UbC* promoter and transfected at comparable plasmid dosages within the same order of magnitude (Fig. 2b, Extended Data Fig. 3d and e); specifically, the growth rate was highest in HEK cells, followed by HeLa cells, and lowest in cultured neurons. We further tested whether CytoTape can capture physiological processes in HeLa cells. We expressed a V5-epitope–tagged reporter under the heat shock-responsive human *HSPA1A* promoter^74^ and applied heat shock stimulation (42 °C, 1 h) in HeLa cells (Extended data Fig. 3c). We observed a clear increase in V5 signal towards the CytoTape termini, indicating that CytoTape can encode physiological signals in HeLa cells. Collectively, these findings suggest CytoTape is a flexible and versatile intracellular recording platform across cell types, promising to achieve enhanced temporal scalability of recording durations.

**Fig. 3.**
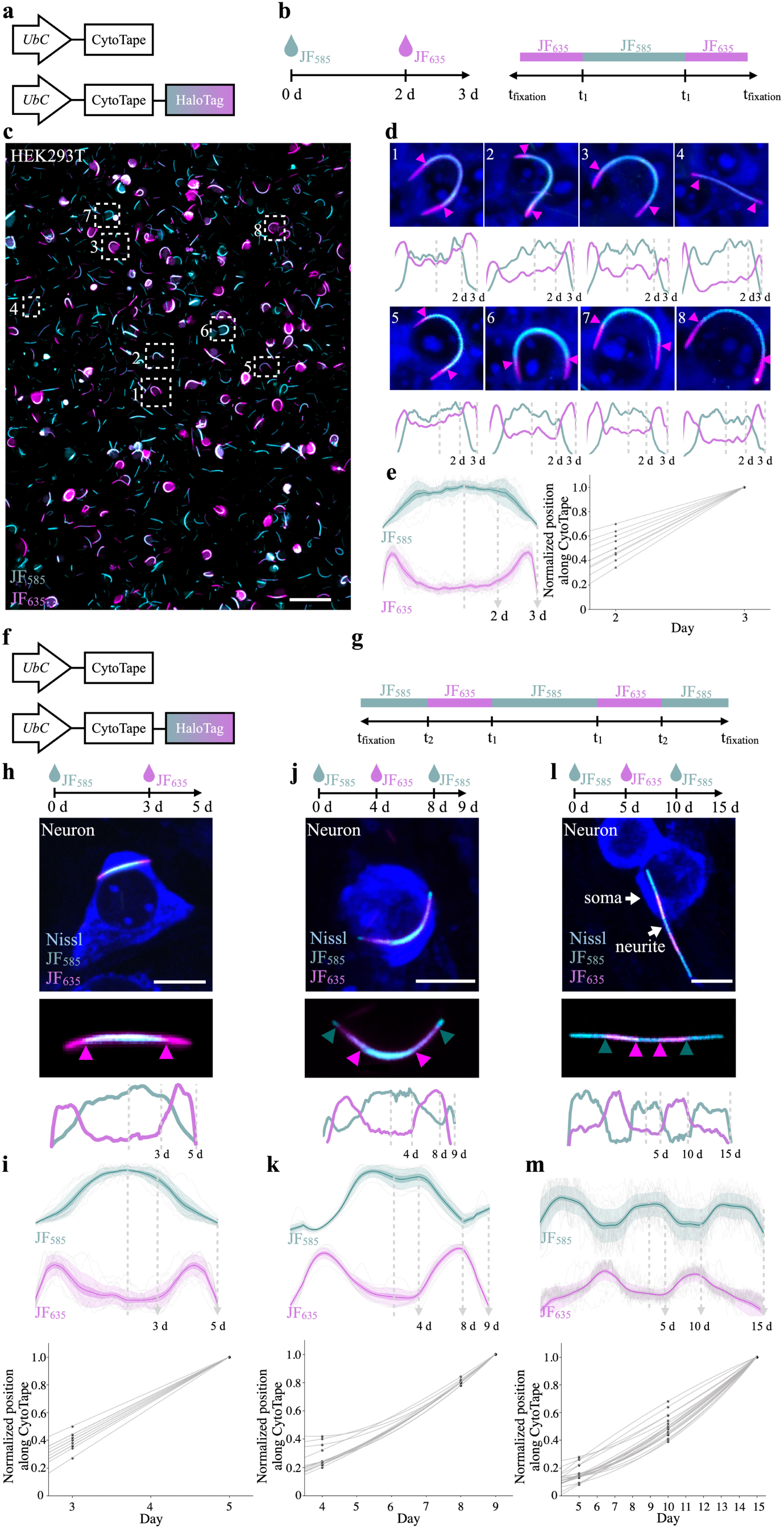
Development of timestamps via sequentially labeled dyes along CytoTape. (**a**) Schematic of the constructs transfected into HEK cells. CytoTape-HaloTag incorporates JF dyes, permitting labeling of the protein assembly with fiducial timestamps. (**b**) Time points of JF_585_ and JF_635_ addition (left panel) and expected dye distribution along the protein fiber (right panel). (**c**) Low-magnification images of CytoTape labeled with JF_585_ and JF_635_ in HEK cells, which is taken after fixation on day 3, and Nissl staining and immunostaining against the HA tag. Scale bar, 100 µm. Some fibers do not display clear timestamp transitions in this low magnification image, because the optimal image contrasts for each color channel to display the timestamp transitions are different across individual cells. See (**d**) for the images of individual cells with the optimal image contrast to display the timestamp transitions. The success rate of CytoTape showing clear timestamps is shown in Extended Data Fig. 4i. (**d**) Enlarged confocal images of HEK cells indicated in (**c**). Arrows in images indicate the positions of dye switches within the protein fiber. Fluorescence line profiles of JF_585_ and JF_635_ are shown below each confocal image. (**e**) Statistical analysis (left panel) of fluorescence line profiles from the experiments described in (**b**) n = 10 CytoTapes from 10 HEK cells from two cultures, and interpolation between the spatial axis along fiber and the time axis (right panel), using the timestamps from left panel. Time axis recovery via linear curve fitting of timestamps and fixation point. Each raw trace on the left was normalized to its peak to show relative changes before averaging. Thick centerline, mean; darker boundary in the close vicinity of the thick centerline, s.e.m.; lighter boundary, s.d.; light gray thin lines, data from individual CytoTapes. (**f**) Schematic of the constructs transfected into cultured neurons and (**g**) expected dye distribution along the CytoTape. (**h**), (**j**), and (**l**) Timestamps for different temporal scales with varying temporal resolutions. First row, time points of JF_585_ and JF_635_ addition; second row, images of CytoTape at timescales of 5 days, 9 days, and 15 days; third row, enlarged view of CytoTape; fourth row, fluorescence line profiles. Scale bars, 10 µm. CytoTape shown in (**l**) extended into the initial segment of a neurite from the soma. (**i**), (**k**), and (**m**) Top panel, statistical analysis of fluorescence line profiles from the experiments described in (**h**) n = 8 CytoTapes from 8 neurons from two cultures, (**k**) n = 10 CytoTapes from 10 neurons from two cultures, and (**m**) n = 18 CytoTapes from 18 neurons from two cultures. Bottom panel, interpolation between the spatial axis along fiber and the time axis using the timestamps from top panel. Time axis recovery via B-spline curve fitting of timestamps and fixation point. Each raw trace was normalized to its peak to show relative changes before averaging. Thick centerline, mean; darker boundary in the close vicinity of the thick centerline, s.e.m.; lighter boundary, s.d.; light gray thin lines, data from individual CytoTapes. Normalized position along CytoTape is calculated as dividing the distance along the half fiber starting from the optimal split point by the total length of this half fiber. The details of the interpolation between the spatial axis along fiber and the time axis using timestamps are described in **Methods**.

### 2.3 Recovery of continuous time axis from timestamps along CytoTape

Protein-assembly-based recording systems convert the time dimension into the spatial dimension and it is critical to establish the conversion relationship between the two dimensions. In XRI, a chemically inducible Cre system was used to initiate time encoding, enabling a global time calibration across cells. However, a single global time axis does not account for the variabilities in the nucleation time (*i.e.*, the time when the fiber begins to form) and the elongation rate of protein assembling across cells and cell types (Fig. 2b, Extended data Fig. 3d and e), and thus the precision of the space-to-time conversion is limited by these variabilities. Establishing individualized time courses at single-cell resolution instead would allow for more precise temporal interpretation of protein-assembly-based recordings, with each fiber recording and maintaining its own internal time axis. The iPAK4-based system demonstrated that the labeling of HaloTag molecules along linear protein assemblies via temporal switching of multiple HaloTag ligand dyes has effective time constants below 1 hour^39^, providing sufficient temporal precision for our purpose to revolve and record long-term (multi-day and multi-week) cellular physiological activities. Therefore, we employed the HaloTag/dyes system as timestamps along the CytoTape. To implement this, we fused HaloTag to the C-terminus of the CytoTape monomer, which was expressed under the *UbC* promoter (referred to as the “timestamp monomer” throughout the paper) (Fig. 3a). By sequentially switching dyes of distinct colors, and using the cell fixation event as the final timestamp (corresponding to the termini of the fiber), we introduced discrete timestamps along the fiber to calibrate its own time axis. To recover the continuous time axis along the CytoTape, we calculated the relationship between the spatial axis and the time axis by spatiotemporal interpolation of the timestamps. Although this interpolation introduces time uncertainties in fiber segments in between the timestamps, the resulting time accuracy (< 1 day) from day-scale dye-switches is sufficient for our purpose to resolve days- and weeks-long cellular events. Nevertheless, we speculate that increasing the density of timestamps along the fiber improves the precision of the reconstructed time axis, enabling more accurate temporal profiling at single-cell resolution.

We first tested HaloTag-based time encoding in HEK and HeLa cells by co-expressing the CytoTape structural and timestamp monomers. On the day of plasmid transfection into HEK cells, JF_585_ (Janelia Fluor) dye was added to the culture medium. Two days later, the medium was washed five times and then replaced with JF_635_-containing medium (Fig. 3b). We observed that the dye-switching events are successfully recorded as timestamps along individual fibers across the cell population (Fig. 3c), resulting in a decline in JF_585_ intensity accompanied by a simultaneous rise in JF_635_ intensity upon dye switch (Fig. 3d). Additionally, we observed that the positions of timestamps along fibers varied among cells, confirming that single-cell–level timestamps can improve the precision of the reconstructed time axis (Fig. 3e, left panel). Since this case included only one dye-switching event and the fixation event, we applied a linear interpolation to approximate the elongation kinetics, resulting in a low-precision estimate of the continuous time axis (Fig. 3e, right panel). We also tested this timestamp approach in HeLa cells (Extended Data Fig. 4a). JF_635_ dye was added on day 0, switched to JF_585_ on day 2, then switched back to JF_635_ on day 3, and cells were fixed on day 4 (Extended Data Fig. 4b). As expected, two distinct dye-switching events were encoded along the CytoTape fibers (Extended Data Fig. 4c), allowing us to generate a more precise time axis via interpolation (Extended Data Fig. 4d). We further tested a two-dye switching scheme (JF_585_ and JF_635_) and a three-dye switching scheme (JF_585_, JF_635_, and JF_503_) for timestamps in HEK cells, which also performed well (Extended Data Fig. 4e-h).

Next, we tested whether CytoTape recording is temporally scalable, achieving user-defined control of temporal resolution and recording duration by adjusting both the timing of dye switches and the duration of fiber elongation. To demonstrate this unique capability (Fig. 3f and g), we tested three distinct temporal configurations in cultured neurons: (1) a single dye switch at day 3 followed by fixation at day 5 (Fig. 3h, top row); (2) two dye switches at days 4 and 8 followed by fixation at day 9 (Fig. 3j, top row); and (3) two dye switches at days 5 and 10 followed by fixation at day 15 (Fig. 3l, top row). Single-fiber results (Fig. 3h, j, and l, bottom row) and statistical analysis (Fig. 3i, k, and m, top panel) showed one signal transition in the single-dye-switch condition and two transitions in the double-dye-switch condition. For each case, we performed interpolation between the spatial axis along fiber and the time axis using the dye switching and cell fixation time points, to recover the continuous time axis along the fiber (Fig. 3i, k, and m, bottom panel). These results validate that both temporal resolution and timescale can be flexibly defined by the user based on specific experimental needs. Finally, we quantified the percentage of CytoTape that successfully incorporated HaloTag dye timestamps. In both cultured neurons and HEK cells, over 90% of fibers showed timestamp incorporation, while in HeLa cells, the success rate exceeded 80% (Extended data Fig. 4i).

**Fig. 4.**
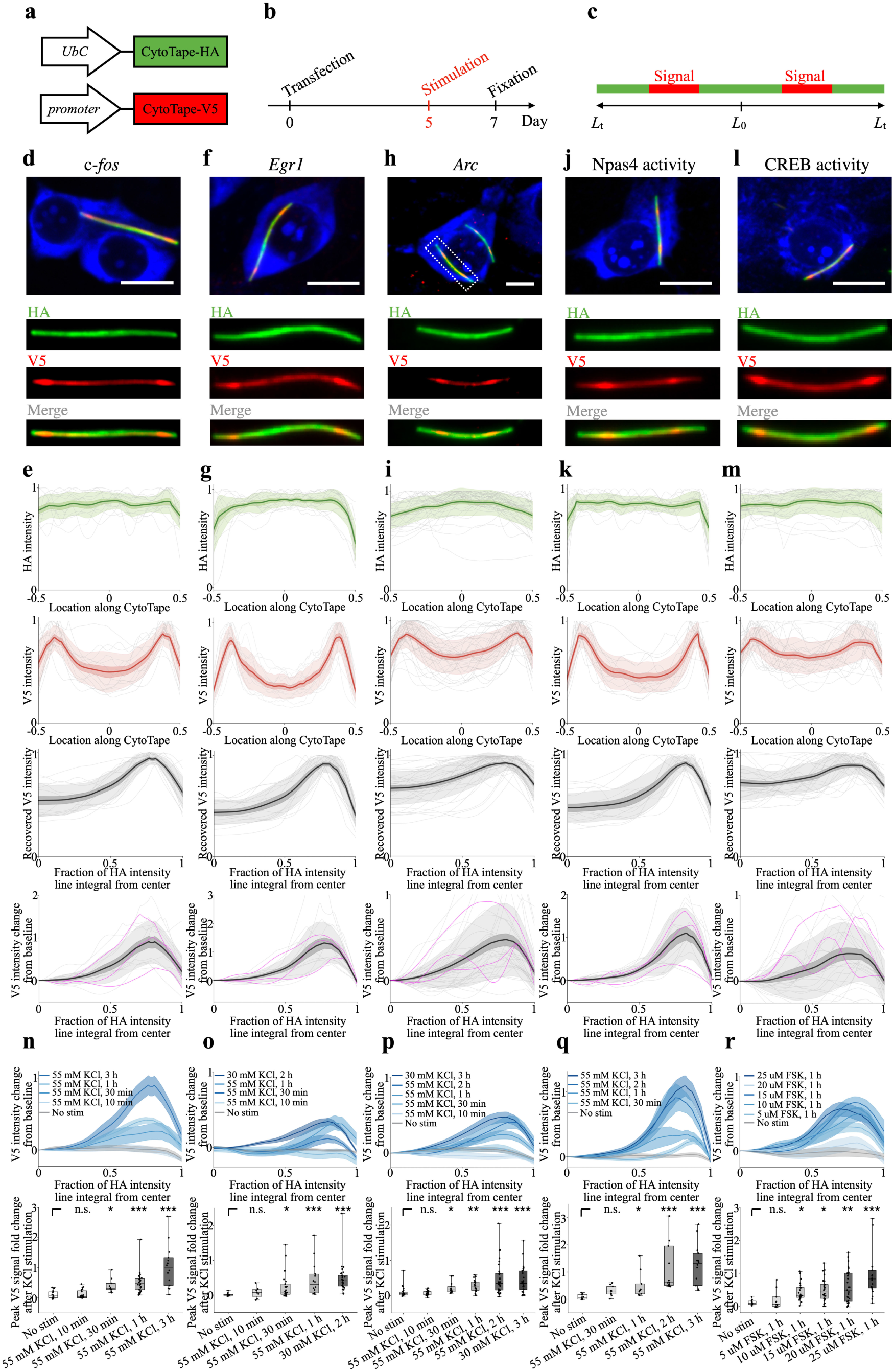
Development of transcriptional recorders for tracking c-*fos*, *Arc*, *Egr1*, Npas4 activity, and CREB activity-dependent-promoter-driven expression histories with CytoTape. (**a**) Schematic of the constructs transfected into cultured neurons and (**b**) the experimental timeline. CytoTape-V5, the signal monomer, is driven under an activity-dependent promoter of interest. (**c**) Expected HA tag and V5 tag distribution along the CytoTape. (**d**), (**f**), (**h**), (**j**), and (**l**) Representative images of cultured neurons expressing the constructs shown in (**a**). c-*fos*, SARE-ArcMin, *Egr1* (also known as zif/268), *N*-RAM, and 6x*CRE*-CMVMin promoters that are responsive to transcriptional activities of c-*fos*, *Arc*, and *Egr1* genes and transcription factor activities of Npas4 and CREB proteins, respectively, were used to drive the signal monomer expression. Images were captured after fixation, Nissl staining, and immunostaining for HA and V5 tags. Neurons were stimulated with 55 mM KCl for 3 h for c-*fos*, 30 mM KCl for 2 h for *Egr1*, 55 mM KCl for 3 h for *Arc*, 55 mM KCl for 3 h for Npas4, and 10 µM forskolin (FSK) for 1 h for CREB. Enlarged views of the CytoTape in the top-row panels are shown in the three rows of rectangular panels below. Scale bars, 10 µm. (**e**), (**g**), (**i**), (**k**), and (**m**) Profiles of HA and V5 signal intensities along CytoTape, based on the experiment in (**b**). First row, HA intensity profile; second row, V5 intensity profile; third row, recovered V5 signal, calculated from the intensity profiles, plotted as a function of the fraction of the HA intensity line integral; fourth row, V5 signal relative to baseline (calculated as the ratio of V5 signal to the center V5 signal) plotted as a function of the fraction of the HA intensity line integral. In the first three rows, raw traces were normalized to their peaks to highlight relative changes before averaging. Data were from c-*fos*, n = 13 CytoTapes (13 neurons, two cultures); *Egr1*, n = 29 CytoTapes (29 neurons, two cultures); *Arc*, n = 26 CytoTapes (26 neurons, three cultures); Npas4, n = 14 CytoTapes (14 neurons, two culture); CREB, n = 11 CytoTapes (11 neurons, two cultures). Thick centerline, mean; darker boundary near the centerline, s.e.m.; lighter boundary, s.d.; thin light lines, intensity profiles from all individual CytoTapes; thin magenta lines, three representative intensity profiles for each group. (**n**), (**o**), (**p**), (**q**) and (**r**) Top panel, V5 signal relative to baseline with different KCl or FSK stimulations. Centerline, mean; shaded boundary, s.e.m. The sample size (n) is listed below as (X, Y, Z), which dentotes n = X protein assemblies from Y neurons from Z cultures. (**n**) c-*fos*: “55 mM KCl, 3 h”, (13, 13, 2); “55 mM KCl, 1 h”, (8, 8, 2), “55 mM KCl, 30 min”, (26, 26, 4), “55 mM KCl, 10 min”, (20, 20, 3), “No Stim”: (13, 13, 2). (**o**) *Egr1*: “55 mM KCl, 3 h”, (29, 29, 2); “55 mM KCl, 1 h”, (14, 14, 3); “55 mM KCl, 30 min”, (23, 23, 2); “55 mM KCl, 10 min”, (8, 8, 2); “No Stim”, (12, 12, 2). (**p**) *Arc*: “30 mM KCl, 3 h”, (26, 26, 3); “55 mM KCl, 2 h”, (36, 36, 2); “55 mM KCl, 1 h”, (20, 20, 2); “55 mM KCl, 30 min”, (15, 15, 2); “55 mM KCl, 10 min”, (27, 27, 3); “No Stim”, (24, 24, 2). (**q**) Npas4 activity: “55 mM KCl, 3 h”, (14, 14, 2); “55 mM KCl, 2 h”, (11, 11, 2); “55 mM KCl, 1 h”, (10, 10, 2); “55 mM KCl, 30 min”, (10, 10, 2); “No Stim”, (11, 11, 2). (**r**) CREB activity: “25 µM FSK, 1 h”, (21, 21, 3); “20 µM FSK, 1 h”, (27, 27, 2); “15 µM FSK, 1 h”, (26, 26, 3); “10 µM FSK, 1 h”, (26, 26, 3); “5 µM FSK, 1 h”, (11, 11, 2); “No Stim”: (9, 9, 2). Bottom panel, box plots comparing (top panel) peak relative change in V5 signal. *, *P <* 0.05; **, *P <* 0.01; ***, *P <* 0.001; n.s., not significant; Kruskal–Wallis analysis of variance followed by Dunn’s post hoc tests. Middle line in box plot, median; box boundary, interquartile range; whiskers, minimum and maximum; black dots, individual data points. See **Supplementary Table 2** for details of statistical analysis.

### 2.4 Development of CytoTape-based transcriptional recorders to measure gene regulation dynamics

Immediate early genes (IEGs) play a critical role in cell biology, acting as rapid responders to diverse stimuli and serve as gateways in the regulation of many cellular processes, such as those associated with gene expression, cell cycles and states, and cell plasticity in both health and disease^75–79^. We tested CytoTape with four well-characterized IEG promoters of c-*fos*^80^, *Egr1*^72^, *Arc*^81^, and Npas4^82, 83^, which have been used extensively to link reporter expression to these IEG activities. We also tested the cAMP response element-binding protein (CREB)-responsive promoter with CytoTape^84^, since the transcription factor activity of CREB has been reported to play a key role in regulating IEG expression and is involved in a broad range of cellular processes, including cell proliferation, differentiation, survival, immune responses, calcium signaling, metabolism, and stress adaptation^85–88^. We previously validated the fidelity of the XRI recording system in cultured neurons, demonstrating the protein ticker tape-based readout of c-*fos*-promoter activity not only recapitulated expected transcriptional kinetics reported in literature but also closely mirrored the expression patterns captured by conventional c-*fos*-GFP live imaging under identical stimulation conditions. This direct comparison indicates that temporal signals from activity-dependent promoters can be reliably embedded and decoded from self-assembling protein structures. We used the IEG- or CREB-responsive promoter to drive expression of the CytoTape monomer fused to the V5 epitope tag, *i.e.*, the signal monomer, and co-expressed it with the structural monomer under the constitutive *UbC* promoter in cultured neurons, to enable post-fixation readout of the recorded promoter activities via the V5 signal intensity profile along the fiber (Fig. 4a). The signal monomer plasmid was diluted to 25% of the amount of the structural monomer plasmid in transfection to ensure that the structural monomers dominate the fiber formation and elongation, thereby providing a consistent substrate for the incorporation of signal monomers over time. Neurons were stimulated on day 5 after transfection with either KCl, to induce depolarization and activate IEG activities^73^, or forskolin (FSK), to raise intracellular cAMP levels and activate the CREB activity^16^, and then fixed and immunostained against HA and V5 tags on day 7 (Fig. 4b and c). As expected, in KCl or FSK stimulated neurons, we observed low pre-stimulation baseline of V5 immunofluorescence intensity at the center of the CytoTape fiber and peak(s)-like V5 intensity profiles on each of the two sides of the fiber that eventually declined towards the fiber termini (Fig. 4d–m). This characteristic pattern was absent in unstimulated neurons, where V5 signals remained at a uniformly low baseline level along each fiber (Extended Data Fig. 5). Although when averaged across cells, the mean signal appeared to be a single peak waveform, we found that individual neurons could exhibit complex waveform patterns. For example, *Arc* and CREB signals in some neurons display two peaks following a single KCl stimulation and FSK stimulation, respectively. This result indicates that the temporal dynamics of these recorded transcriptional activities over days are highly heterogeneous across cells even under identical stimulation conditions (Fig. 4d-m, fourth row, three representative traces are highlighted in magenta). We also compared the recorded signals across multiple CytoTape assemblies within the same neuron, when there are more than one assembly per neuron (Extended Data Fig. 5n). We found that distinct assemblies within the same cell reported highly similar signal waveforms, suggesting that CytoTape robustly records transcriptional activity in cells and that the cell-to-cell variability observed in our previous experiments reflects biological differences in cellular transcriptional responses rather than artifacts introduced by the CytoTape assembly. To assess the sensitivity of the CytoTape recorders, we titrated a series of doses and durations of KCl and FSK to neurons expressing the structural monomer and the signal monomer under IEG- and CREB responsive-promoters. We found stronger and longer stimulation protocols yielded higher and steeper peak(s) in the V5 signal waveforms than weaker or shorter stimulations (Fig. 4n–r), demonstrating that CytoTape is an analog recorder that captures not only the transcriptional events but also the amplitudes of these events in response to variable physiological inputs. Importantly, in the absence of stimulation, V5 signals remained flat at the baseline level over time, confirming that the observed responses were specifically induced by the applied stimuli rather than spontaneous neuronal activities. Interestingly, CytoTape also captured IEG-promoter-driven gene expression in mouse hippocampal glial cells that were co-cultured with neurons, showing its potential for studying gene regulation dynamics in glia and the coordination of gene regulation between neuronal and glial cell populations (Extended Data Fig. 6).

### 2.5 Multiplexed, multi-week, multi-event recording of activity-dependent promoter-driven expression histories via CytoTape

We next evaluated whether CytoTape could simultaneously record gene regulation dynamics and timestamps by integrating activity-dependent promoters with the previously described timestamp strategy via HaloTag and dye switches. We co-expressed the structural monomer, the timestamp monomer, and the c-*fos* signal monomer in HEK cells^89, 90^ (Extended Data Fig. 7a). JF_635_ dye was added on day 0 and followed by a switch to JF_585_ on day 2 concurrent with FSK stimulation (10 µM, 1 h) (Extended Data Fig. 7b). Cells were fixed on day 3. Following immunostaining, CytoTape fibers exhibited clear dye switches and c-*fos*-promoter-driven CytoTape monomer expression across the cell population (Extended Data Fig. 7c, left panel). Spatially localized V5 signal bands were observed along the fibers, indicating capturing of stimulus-induced transcriptional activity (Extended Data Fig. 7c, right panel). To temporally resolve these signals, we reconstructed a continuous time axis by interpolating the timestamps and fixation time point, and then aligned the resulting timeline with the V5 intensity profile. Plotting the V5 signal relative to baseline as a function of reconstructed time revealed a distinct peak after day 2, consistent with the actual timing of FSK stimulation (Extended Data Fig. 7d). Furthermore, we observed variability in the shape and amplitude of V5 signal waveforms across individual cells, suggesting heterogeneous c-*fos*-promoter-driven transcriptional dynamics within the cell population. To test whether CytoTape could resolve sequential transcriptional events, we performed two FSK stimulations separated by 1 day in the same HEK cell culture (Extended Data Fig. 7e). We recovered two discrete peaks in the V5 signal profile, corresponding to the timing of each stimulation (Extended Data Fig. 7g, left panel). We further tested the heat shock-responsive *HSPA1A* promoter to evaluate whether CytoTape could record cellular stress responses in HEK cells, a process that gained popularity in cell stress and survival research^74, 91, 92^ (Extended Data Fig. 7h). After heat shock treatment, we observed clear V5 signal induction along fibers (Extended Data Fig. 7j, left panel), while the untreated control group showed no significant changes in the V5 signal (Extended Data Fig. 7g and j, right panel). In addition, we demonstrated CytoTape recording of the time courses of IEG-promoter-driven activities (c-*fos*, *Egr1*, *Arc*, and Npas4) in response to KCl stimulation in cultured neurons with timestamps (Extended Data Fig. 8).

To evaluate CytoTape’s potential for long-term recording, we extended the recording period to 21 days. Since we already tested CytoTape’s ability to record in the initial week in experiments with shorter recording durations (Extended Data Fig. 7 and 8), here we focused on testing whether the timestamps and cellular signals in the final days of the 21-day period could be reliably recorded and recovered. We again co-expressed the structural monomer, the timestamp monomer, and the c-*fos* signal monomer in cultured neurons (Fig. 5a). JF_585_ dye was added on day 0, followed by JF_635_ on day 18, and JF_503_ on day 20. Cells were fixed on day 21 (Fig. 5b). Neurons were stimulated with 55 mM KCl for 1 h on both day 18 and day 20. After 21 days of CytoTape growth, CytoTape formed long, flexible fibers (Fig. 5c). We then recovered the time axis from timestamps and analyzed the V5 signal intensity along the fiber and observed two distinct peaks corresponding to the two sequential stimulation events, with the V5 signal onset occurring after each stimulation. Additionally, we observed variability in the shape and amplitude of V5 signal waveforms across individual cells under identical KCl stimulation conditions, suggesting heterogeneity in transcriptional dynamics in neurons (Fig. 5d). We further analyzed the amplitudes of the two peaks following the two identical KCl stimulations across neurons and found no significant difference between the two peaks (Fig. 5e, left panel). We also calculated the full width at half maximum (FWHM) of the V5 signal peaks (Fig. 5e, right panel) and found that the FWHMs were on the order of several hours and were comparable between the two sequential V5 signal peaks, indicating that CytoTape can resolve hours-scale transcriptional activity.

**Fig. 5.**
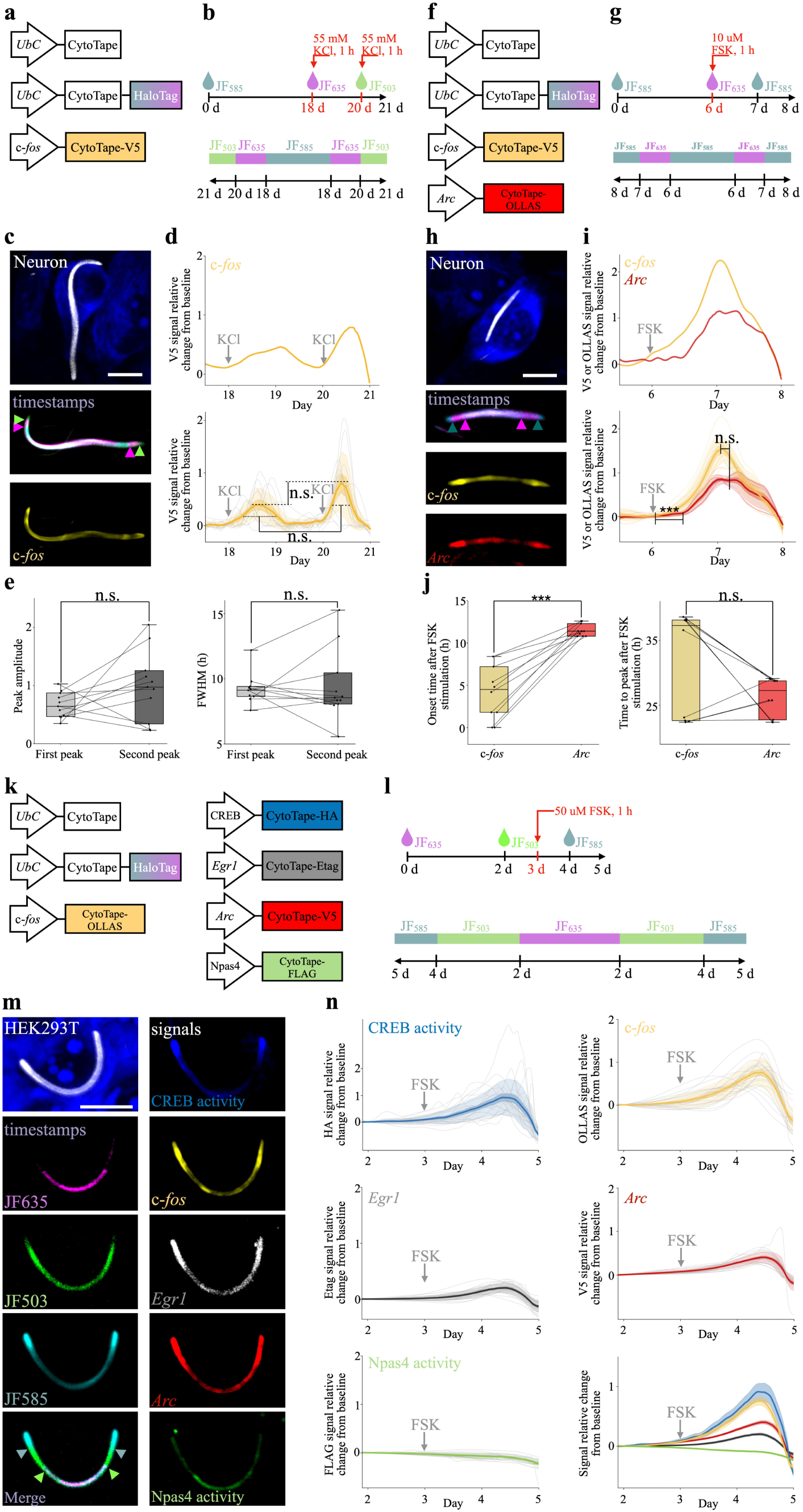
Recording activity-dependent-promoter-driven expression histories with CytoTape. (**a**) Schematic of the constructs transfected into cultured neurons, (**b**) the experimental timeline (top panel), and the expected dye distribution along the CytoTape (bottom panel). CytoTape-V5 refers to CytoTape fused to the V5 tag. (**c**) Images of neurons expressing the constructs shown in (**a**). Images were captured after fixation, Nissl staining, and immunostaining for V5 tag. Neurons were stimulated with 55 mM KCl for 1 h on day 18 and day 20. First row, a composite image of all imaged channels showing the CytoTape in the cell; second row, the image in the timestamp channels; third row, the image in the c-*fos* signal channel. Scale bar, 10 µm. CytoTape shown in (**c**) extended into the initial segment of a neurite from the soma. (**d**) Top panel, c-*fos* signal (from (**c**)) relative change from baseline plotted against recovered time (interpolated with timestamps and fixation time) after calcium phosphate transfection. Bottom panel, statistical analysis of c-*fos* signal relative change from baseline plotted against recovered time after calcium phosphate transfection. Thick centerline, mean; darker boundary near the centerline, s.e.m.; lighter boundary, s.d.; thin gray lines, individual CytoTape data. Data are from n = 11 CytoTapes from 11 neurons from two cultures. (**e**) Statistical analysis of the peak amplitudes (left panel) and widths (full width at half maximum, FWHM) (right panel) between the sequential V5 signals. n.s., not significant; Wilcoxon signed-rank test. Middle line in box plot, median; box boundary, interquartile range; whiskers, minimum and maximum, not indicated in the box plot; black dots, individual data points. (**f**) Schematic of the constructs transfected into cultured neurons, (**g**) the experimental timeline (top panel), and the expected dye distribution along the CytoTape (bottom panel). CytoTape-V5 and CytoTape-OLLAS refers to CytoTape fused to the V5 tag and the OLLAS tag, respectively, as signal monomers. (**h**) Images of neurons expressing the constructs shown in (**f**). Images were captured after fixation, Nissl staining, and immunostaining for V5 and OLLAS tags. Neurons were stimulated with 10 µM FSK for 1 h on day 6. First row, a composite image of all imaged channels showing the CytoTape in the cell; second row, the image in the timestamp channels; third row, the image in the c-*fos* signal channel; fourth row, the image in the *Arc* signal channel. Scale bar, 10 µm. (**i**) Top panel, c-*fos* and *Arc* signal (from (**h**)) relative change from baseline plotted against recovered time (interpolated with timestamps and fixation time) after calcium phosphate transfection. Bottom panel, statistical analysis of c-*fos* and *Arc* signals relative change from baseline plotted against recovered time after calcium phosphate transfection. Thick centerline, mean; darker boundary near the centerline, s.e.m.; lighter boundary, s.d.; thin light lines, individual CytoTape data. Data are from n = 11 CytoTapes from 11 neurons from two cultures. (**j**) Statistical analysis of the onset time (left panel) and the time to peak (right panel) between the c-*fos* and *Arc* signals. ***, *P <* 0.001; n.s., not significant; Wilcoxon signed-rank test. Middle line in box plot, median; box boundary, interquartile range; whiskers, minimum and maximum; black dots, individual data points. (**k**) Schematic of the constructs transfected into HEK cells, (**l**) the experimental timeline (top panel), and the expected dye distribution along the CytoTape (bottom panel). CytoTape-HA refers to CytoTape fused to the HA tag, CytoTape-Etag refers to CytoTape fused to the Etag tag, CytoTape-OLLAS refers to CytoTape fused to the OLLAS tag, CytoTape-V5 refers to CytoTape fused to the V5 tag, and CytoTape-FLAG refers to CytoTape fused to the FLAG tag. (**m**) Images of a representative HEK cell expressing the constructs shown in (**k**). Images were captured after three rounds of processing: first, imaging of JF dyes and the HA tag followed by fixation and immunostaining for HA tag; second, photobleaching of JF dye signals followed by immunostaining for HA, Etag, OLLAS, V5, and FLAG tags (with the HA tag used for alignment of CytoTape between the two rounds of immunostaining); third, Nissl staining. HEK cells were stimulated with 50 µM FSK for 1 h on day 3. Left, first row, the CytoTape within the cell; left, second row, JF_635_ dye; left, third row, JF_503_ dye; left, fourth row, JF_585_ dye; left, fifth row, merge of JF_635_, JF_503_, and JF_585_ dyes. Right, first row, CREB signal; right, second row, c-*fos* signal; right, third row, *Egr1* signal; right, fourth row, *Arc* signal; right, fifth row, Npas4 signal. Scale bar, 10 µm. (**n**) HA (left, first row), OLLAS (right, first row), Etag (left, second row), V5 (right, second row), and FLAG (left, third row) signal relative change from baseline plotted against recovered time (interpolated with timestamps and fixation time) after calcium phosphate transfection. Right, third row, comparison of c-*fos*, *Egr1*, *Arc*, CREB activity, and Npas4 activity. Data are from n = 20 CytoTapes from 20 HEK cells from two cultures. Thick centerline, mean; darker boundary near the centerline, s.e.m.; lighter boundary, s.d.; thin gray lines, individual CytoTape data. See **Supplementary Table 2** for details of statistical analysis.

To evaluate whether CytoTape can achieve simultaneous recording of multiple IEG-promoter-driven signals—a desired capability for studying complex processes such as development, disease progression, and neural computation^7, 93, 94^—we co-expressed the structural monomer, the timestamp monomer, the c-*fos* signal monomer fused to the V5 tag, and the *Arc* signal monomer fused to the OLLAS tag in cultured neurons (Fig. 5f). Dye switches were performed on day 6 and day 7 as timestamps, and cells were fixed on day 8 (Fig. 5g), and neurons were stimulated with 10 µM FSK for 1 h on day 6. We observed that both c-*fos*-promoter-driven (V5) and *Arc*-promoter-driven (OLLAS) signals increased after FSK stimulation, validating that CytoTape can simultaneously record multiple distinct gene regulation dynamics along the same fiber and within the same cell (Fig. 5h and i). Further analysis of the signal waveforms revealed that, while the c-*fos*-promoter-driven signal initiated earlier than the *Arc*-promoter-driven signal, the two signals exhibited synchronized peak timing (Fig. 5j). To assess the multiplexing capacity of CytoTape, we co-expressed the structural monomer, the timestamp monomer, the CREB activity signal monomer fused to the HA tag, the *Egr1* signal monomer fused to the E tag, the c-*fos* signal monomer fused to the OLLAS tag, the *Arc* signal monomer fused to the V5 tag, and the Npas4 activity signal monomer fused to the FLAG tag in HEK cells (Fig. 5k). Timestamps were introduced by dyes switches on days 2 and 3, a 1-h stimulation with 50 µM FSK was applied on day 3, and cells were fixed on day 5 (Fig. 5l). We found that CytoTape can simultaneously record both the timestamps and the responses in the five distinct gene regulation dynamics in response to FSK stimulation, all within a single protein fiber (Fig. 5m). Unlike other signals, we did not observe a significant change in the Npas4 signal (Fig. 5n), in agreement with the literature that the Npas4 promoter cannot be activated by FSK^95^.

### 2.6 CytoTape provides new insights into temporal principles of gene regulation dynamics

We next applied CytoTape to investigate the temporal relationship between the CREB activity and the Fos activity—two extensively studied transcription factor activities involved in cell proliferation, signal integration, and transcriptional regulators in human cell lines such as the HEK cells^96^. While CREB is canonically recognized as an upstream activator of Fos, we explored whether they could exhibit complex dynamical features and non-linear temporal couplings beyond simple, linear correlations^97, 98^. Such regulatory complexity could support history-dependent responses, nonlinear signal integration, and memory-like behaviors of gene regulatory networks^99^. To independently track the dynamics of each component, we co-expressed the structural monomer, the timestamp monomer, and two signal monomers driven by the CREB-responsive promoter (regulated by phosphorylated CREB protein) and the *F*-RAM promoter (regulated by Fos protein activity)^82^ (Fig. 6a), respectively, in HEK cells for 4 days (Fig. 6b), with FSK (50 μM, 1 h) applied on day 2.25 (6 h after day 2) to activate the cAMP– CREB pathway.

**Fig. 6.**
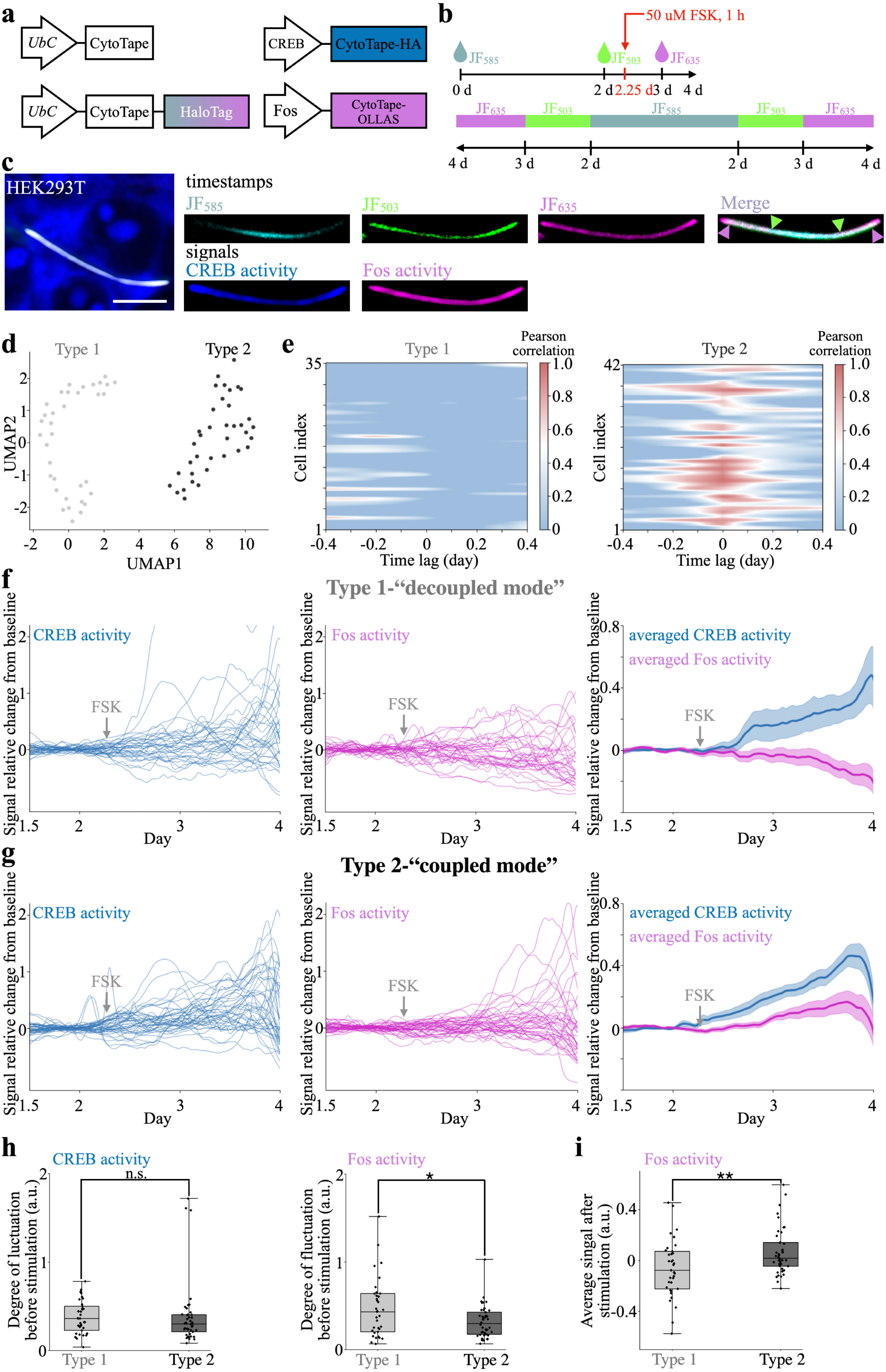
Multiplexed recording CREB and Fos activities with CytoTape. (**a**) Schematic of the constructs transfected into HEK cells, (**b**) the experimental timeline (top panel), and the expected dye distribution along the CytoTape (bottom panel). The CREB activity and Fos activity is detected by 6x*CRE*-CMVMin promoter and *F*-RAM promoter, respectively. (**c**) Images of HEK cells expressing the constructs shown in (**a**). Images were captured after fixation, Nissl staining, and immunostaining for HA and OLLAS tags. HEK cells were stimulated with 50 μM FSK for 1 h at day 2.25 (6 h after day 2). Left panel, the CytoTape in the HEK cells. First row, timestamps; second row, CREB activity and Fos activity. Scale bar, 10 μm. (**d**) UMAP plot based on time-lagged correlations between CREB activity and Fos activity across single cells. Data are from n = 77 CytoTapes from 77 HEK cells from five cultures. (**e**) Heatmaps illustrate time lagged correlations between CREB and Fos signals for Type 1 (left panel, n = 35 CytoTapes from 35 HEK cells) and Type 2 (right panel, n = 42 CytoTapes from 42 HEK cells) HEK cells. x-axis represents a time-lagged shift window of ±0.4 days (1/10 of the entire recording duration) (both left and right panels). y-axis represents HEK cells. (**f**) Type 1 and (**g**) Type 2, single traces of CREB (left panel) and Fos (middle panel) signals relative change from baseline, plotted against recovered time (interpolated with timestamps and fixation time) after calcium phosphate transfection in two responsive types. Right panel, averaged curve of CREB and Fos signals relative change from baseline plotted against recovered time after calcium phosphate transfection. Thick centerline, mean; darker boundary near the centerline, s.e.m. (**h**) Quantification of the degree of baseline fluctuations in CREB and Fos signals prior to stimulation for Type 1 and Type 2 cells. The degree of baseline fluctuation is quantified by line length, which is calculated by summing the absolute differences between consecutive points, between day 1.5 and day 2.25. (**i**) Average post-stimulation signal of Fos in Type 1 and Type 2 cells. *, *P* < 0.05; **, *P* < 0.01; n.s., not significant; Mann–Whitney U test. Middle line in box plot, median; box boundary, interquartile range; whiskers, minimum and maximum; black dots, individual data points. See **Supplementary Table 2** for details of statistical analysis.

To investigate the temporal relationship of the CREB–Fos transcription factor activities, we calculated their time-lagged correlations within individual cells, to quantify how changes in one signal are correlated with the changes in another. Dimension reduction of the time-lagged correlation analysis revealed two distinct clusters of signal pairs in single cells (Fig. 6d), defined as Type 1 and Type 2. Cells in the Type 1 cluster (“decoupled mode”) displayed low or no correlation between CREB and Fos activities at any time lag, indicating a decoupled regulatory mode (Fig. 6e, left panel). By contrast, cells in the Type 2 cluster (“coupled mode”) showed strong positive correlations around zero-time lag, indicative of tightly coupled CREB and Fos dynamics (Fig. 6e, right panel). This bifurcation was further evident in the recorded waveforms: while Type 2 cells showed a canonical CREB–Fos coupling pattern with post-stimulation Fos induction (Fig. 6g), Fos responses in Type 1 cells appeared to be more irregular and chaotic despite a comparable CREB activation pattern (Fig. 6f). In addition to the differences in CREB and Fos dynamics following FSK stimulation, we hypothesized that their pre-stimulation baseline activity patterns may reflect aspects of cellular states and thus contribute to the divergent post-stimulation responses observed in these two types. Interestingly, although the level of fluctuation (quantified by line length, a popular metric in waveform analysis of neuronal electrophysiology^100^) of CREB activity in the pre-stimulation period was similar between the two types (Fig. 6h), the pre-stimulation fluctuation level of Fos activity was modestly higher in the decoupled group. In addition, following stimulation, decoupled cells exhibited significantly lower Fos activity levels than their coupled counterparts (Fig. 6i). This suggests that a history of a more active Fos activity—potentially reflecting a refractory or saturated regulatory state—may attenuate the cell’s transcriptional response to CREB activation^101^. As mechanistic and causal investigations of this process are beyond the scope of this study, future research may elucidate whether a form of molecular homeostasis is involved, where prior activities or occupancy of associated molecular components dampens the responsiveness of Fos to the subsequent CREB activation—a potential regulatory mechanism that is difficult to capture without simultaneous multiplexed recording at the single-cell level.

Previous studies have reported both the coordinated and differential roles of distinct IEGs in mammalian neurons^82, 102–105^. To investigate how IEGs are coordinately and differentially regulated across time, we applied CytoTape to simultaneously record the transcriptional dynamics driven by the *Arc* promoter and the *Egr1* promoter in primary cultured mouse hippocampal neurons (Fig. 7). *Arc* and *Egr1* were selected for their central and also potentially distinct roles in activity-dependent neuroplasticity: *Arc* has been shown to modulate synaptic strength via AMPA receptor trafficking and cytoskeletal remodeling, while *Egr1* has been reported to drive longer-term gene expression essential for memory consolidation and neuronal adaptation^106^. Although previous studies reported that both can be induced by neuronal depolarization (*e.g.*, via KCl stimulation) or activation of the cAMP signaling pathway (*e.g.*, via FSK stimulation), the long-term temporal correlations of their behaviors within single neurons remains unclear.

**Fig. 7.**
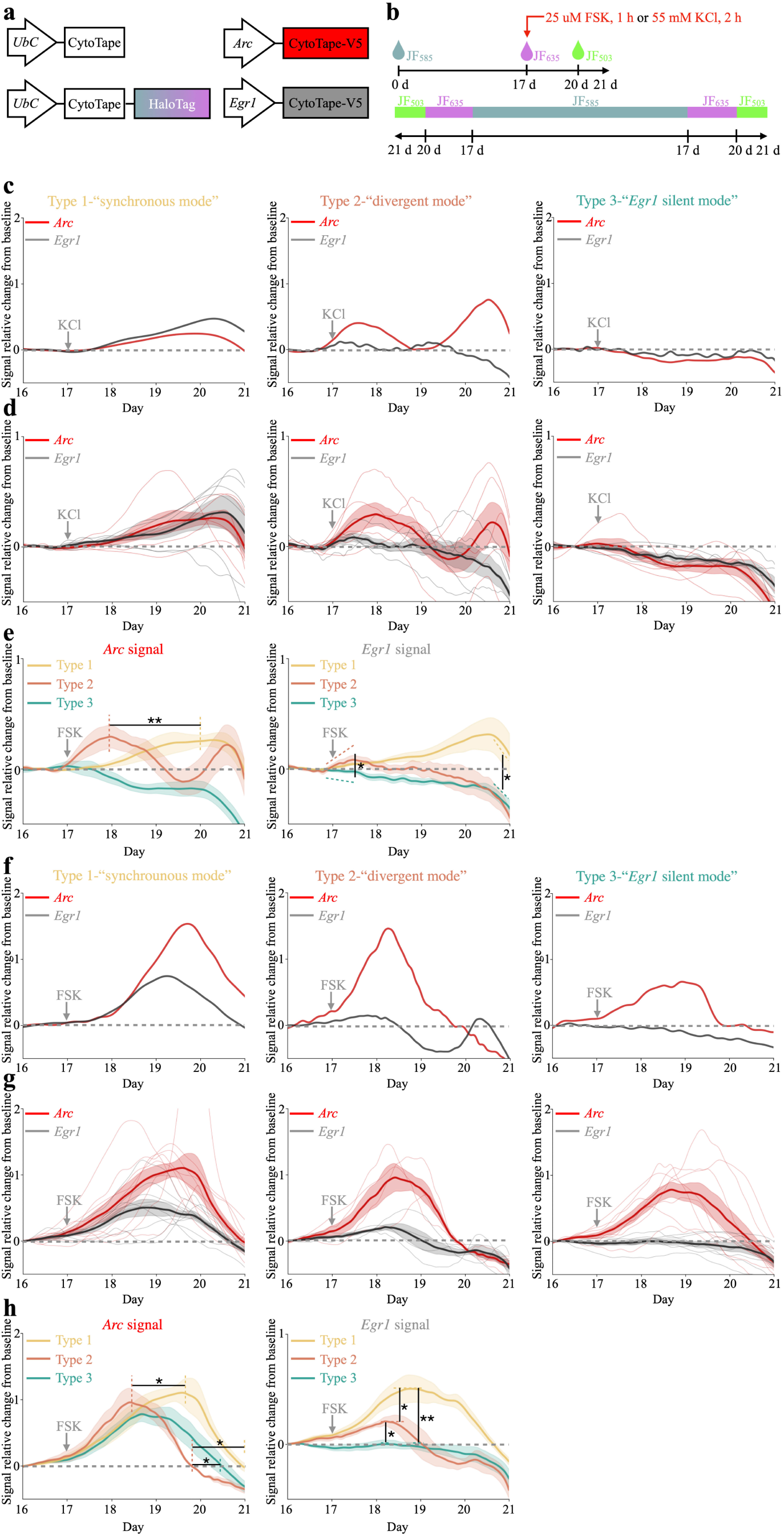
Multiplexed recording *Arc*- and *Egr1*-promoter-driven expression histories with CytoTape. (**a**) Schematic of the constructs transfected into cultured neurons, (**b**) the experimental timeline (top panel), and the expected dye distribution along the CytoTape (bottom panel). (**c**) Representative single traces of *Arc* and *Egr1* signals relative change from baseline, plotted against recovered time (interpolated with timestamps and fixation time) after calcium phosphate transfection in three responsive types under KCl stimulation. (**d**) Averaged *Arc* and *Egr1* signals relative change from baseline plotted against recovered time after calcium phosphate transfection. Data are from n = 16 CytoTapes from 16 neurons from three cultures. Thick centerline, mean; darker boundary near the centerline, s.e.m.; lighter boundary, s.d.; thin light lines, individual CytoTape data. (**e**) Comparison of *Arc* (left panel) and *Egr1* (right panel) signals across three responsive cell types. (**f**) Representative single traces of *Arc* and *Egr1* signals relative change from baseline, plotted against recovered time (interpolated with timestamps and fixation time) after calcium phosphate transfection in three responsive types under FSK stimulation. (**g**) Averaged *Arc* and *Egr1* signals relative change from baseline plotted against recovered time after calcium phosphate transfection. Data are from n = 21 CytoTapes from 21 neurons from three cultures. Thick centerline, mean; darker boundary near the centerline, s.e.m.; lighter boundary, s.d.; thin light lines, individual CytoTape data. (**h**) Comparison of *Arc* (left panel) and *Egr1* (right panel) signals across three responsive cell types. Thick centerline, mean; darker boundary near the centerline, s.e.m.; lighter boundary, s.d. *, *P <* 0.05; **, *P <* 0.01; n.s., not significant; Kruskal–Wallis analysis of variance followed by Dunn’s post hoc tests. See **Supplementary Table 2** for details of statistical analysis.

To probe the joint transcription dynamics of *Arc* and *Egr1*, we stimulated cultured neurons with either 55 mM KCl for 2 hours or 25 µM forskolin for 1 hour on day 17 after transfection of CytoTape constructs (Fig. 7a), and incorporated sequential HaloTag dye switches to reconstruct continuous temporal trajectories (Fig. 7b). Following KCl stimulation, we observed three distinct transcriptional response patterns (Fig. 7c and d). Type 1 neurons (“synchronous mode”) showed concurrent strong induction of both *Arc* and *Egr1* (Fig. 7c and d, left panel). In Type 2 neurons (“divergent mode”), *Arc* exhibited a biphasic response, with two distinct peaks following a single KCl stimulation, while *Egr1* showed only a weak induction (Fig. 7c and d, middle panel). In contrast, Type 3 neurons (“*Egr1* silent mode”) displayed minimal *Arc* and *Egr1* activation (Fig. 7c and d, right panel). These distinctions were confirmed by side-by-side statistical comparisons of single traces among the three groups (Fig. 7e, Extended Data Fig. 9b). The initiation kinetics of the *Egr1* signal was faster in Type 2 neurons than in Type 3, while no significant difference was observed between Type 1 and Type 2. Type 1 neurons displayed a rapid decline in *Egr1* signal after 3 days of recording, in contrast to the sustained weak response observed in Type 3 neurons. The *Arc* signal in Type 2 neurons peaked twice after stimulation, with the onset and decay of the first peak occurring well before the time when the Arc signal in Type 1 neurons reached their single peaks. Type 3 neurons, on the other hand, showed a more attenuated *Arc* response, distinct from the dynamics observed in Types 1 and 2.

When stimulated with forskolin, we also identified three distinct transcriptional response types, though the signal behaviors differed markedly from those observed under KCl stimulation (Fig. 7f–h). Type 1 neurons (“synchronous mode”) displayed tightly coordinated induction of both *Arc* and *Egr1* signals (Fig. 7f and g, left panel). In type 2 neurons (“divergent mode”), *Arc* exhibited one strong activation, and *Egr1* expression showed oscillatory waveforms (Fig. 7f and g, middle panel). In contrast, Type 3 neurons (“*Egr1* silent mode”) showed robust *Arc* activations with minimal *Egr1* activation (Fig. 7f and g, right panel). Comparative analysis (Fig. 7h, Extended Data Fig. 9c) showed that *Egr1* signal intensity was highest in Type 1 neurons, where a single strong peak of *Egr1* activity was observed. In contrast, Type 2 neurons showed oscillatory *Egr1* waveforms at significantly lower signal amplitudes compared to those in Type 1 neurons. For *Arc*, the overall signal strength was similar across all three neuron types. However, in Type 2 neurons, *Arc* signal reached the peak and then returned to baseline faster than that in Type 1 neurons. These correlation analyses provide a temporal perspective to the previously reported regulatory link between *Arc* and *Egr1* in neurons, suggesting a coupling between the timing of *Arc* transcription and the amplitude and the number of *Egr1* transcriptional event(s)^102, 103^.

The transcriptional modes and waveforms from multiplexed recordings are in agreement with single-activity recordings of *Arc*- and *Egr1*-promoter-driven gene expression, respectively, under identical stimulation conditions (Extended Data Fig. 9a), reporting comparable dynamics of each activity. This result indicates that multiplexing on CytoTape does not alter its ability to record each individual signal, at least at the time scale we validated in this study. In conclusion, these multiplexing results demonstrate the power of CytoTape to dissect within-cell correlations among multiple cellular activities, which are otherwise inaccessible from single-activity recordings alone, facilitating a new kind of study on the complex dynamics and interactions in gene regulation networks as well as their couplings to signaling pathways.

## 3 Discussion

In this study, we developed CytoTape, a genetically encoded, modular protein assembly-based system for multiplexed, analog recording of cellular activities at single-cell resolution and multi-week timescales. Through a combination of AI-guided design and rational design, CytoTape produces thin, flexible, thread-like protein assemblies that achieve long-term, multiplexed recording in both dividing and post-mitotic cell types, complementing prior systems such as XRI and iPAK4 (Extended Data Table 1). CytoTape is an analog recorder that embeds temporal signals along the spatial dimension, recording continuously over the time axis with the recorded signal intensities across time points being directly comparable. This unique feature enables measurement of complex temporal waveforms, a desired capability for analyzing complex cellular dynamics. To achieve spatial encoding of the time axis, we fused CytoTape to HaloTag and introduced sequential switches of HaloTag-ligand dyes that served as discrete timestamps on top of the continuously recorded analog signals along CytoTape assembly. Interpolation of these timestamps and their corresponding real-world time points enabled the reconstruction of individualized, continuous time axes for each cell. We demonstrated that CytoTape can encode transcriptional responses driven by a range of activity-dependent promoters, including immediate early gene activity (c-*fos*, *Egr1*, and *Arc*), transcription factor activity (CREB, Npas4, and Fos), and cellular stress-associated activity (*HSPA1A*), in response to a variety of cellular stimuli such as neuronal depolarization, chemical activation of signaling pathways, and heat shock, across cell types. Furthermore, CytoTape supports simultaneous recording of up to five transcriptional signals within a single fiber and revealed heterogeneous expression dynamics across cell populations. These capabilities establish CytoTape as a powerful platform for spatiotemporally resolved, multiplexed, weeks-long analog recording in live cells.

Compared to nucleic acid-based recording systems which typically require cell lysis, tissue dissociation, or specialized sequencing platforms—CytoTape offers a genetically encoded, protein-based alternative that preserves the integrity of cellular content and the spatial context for applications where this information is important and provide new capabilities to resolve analog signal waveforms in a multiplexable manner within single cells. Readout is performed using standard immunostaining and fluorescence microscopy that only takes a few hours to complete after cell fixation, eliminating the need for high-throughput sequencing or custom instrumentation. Compared to real-time fluorescent reporters such as GCaMP^107^, which captures fast and transient physiological signals, CytoTape provides a complementary capability for scalable analog recording of slower cellular activities over days and weeks and stores it in a stable, chronologically ordered format that can be decoded retrospectively. By enabling multiplexed recording of cellular activities over multi-week timescales, CytoTape could open new directions to decode long-term cellular processes—such as tracking the spatiotemporal signatures and differential roles of IEG activities in neurons^108, 109^, revealing how cellular stress and immune pathways interact within single cancer cells in healthy and diseased states and under drug exposure^110, 111^, and mapping spatially resolved gene expression histories during cell development or regeneration^112, 113^, to name a few.

We focused this initial study on cultured human cell lines (HEK and HeLa cells) and mouse primary cells (cultured neurons and glial cells). These systems are popular in both basic cell biology and disease-relevant research to model proliferative, stress-responsive, and activity-dependent gene regulation programs, respectively. As the scope of this work centers on the development and validation of a novel technology for cell biology research, we prioritized our efforts on mammalian cell culture system that allow highly controlled manipulation and benchmarking of transcriptional signals. Although we anticipate that the general approach described in this paper may ultimately be applicable to *in vivo* applications, akin to how DNA ticker tape systems were initially tested in cell culture^18, 20–26, 28–35^ before expansion to cell lineage reconstruction *in vivo*^19, 27^, we envision the current CytoTape platform to facilitate novel cell biology experiments and discoveries on the temporal principles of gene network dynamics, a critically understudied area, even before future iterations of optimization and validation for tissue-level and *in vivo* deployment.

Several limitations of CytoTape are noted. First, the temporal resolution is limited not only by the rate of CytoTape fiber elongation—which can be tuned by modifying the expression level of structural monomers, such as through user selection of a promoter to desired strength to drive structural monomers—but also by the intrinsic time uncertainty of monomer incorporation onto the assembly in live cells. Even if the fiber elongates rapidly, individual monomers may require minutes to hours to diffuse and integrate into the assembly. This kinetic delay constrains the temporal precision of event encoding, particularly in applications when resolving rapid or closely spaced transcriptional events in time are critical. Second, although CytoTape has been demonstrated to simultaneously record up to five dynamic nodes in gene regulatory networks, the complexity of many intracellular processes may require multiplexing beyond five signals to fully dissect the temporal principles in much larger networks.

Third, the current system is limited to transcriptional readouts via activity-dependent promoters, whereas many physiologically relevant signals—such as transients of fast signaling activities^107^ or translational dynamics^105^—are not directly accessible with this platform. To address these limitations, we envision several future directions below. The application and development of deep learning methods^114–120^ hold great potential for advancing protein sequence design and property prediction for self-assembling protein complexes. Achieving this goal involves two key aspects: protein design and subsequent screening. In the design stage, promising strategies include enhancing model awareness of environmental context in live cells and the distinct features of protein assemblies compared to monomers. This could be achieved by incorporating training datasets annotated with assembly-specific labels, developing coarse-grained generative models tailored for oligomeric complexes, and integrating modules that encode surface features relevant to assembly behavior into the protein representation. Building on these capabilities, we envision that future protein tape systems could achieve dynamic control of fiber growth rates (beyond simply changing the promoter strength of the structural monomer construct) and geometries (perhaps building a single-molecule-thick filament). This would allow application-specific adjustments to the tradeoff between temporal resolution and recording duration, which is intrinsic to a given intracellular protein tape recorder system limited by the maximum allowed information storage along the protein assembly in cells, and also deployment of new assemblies with enhanced information storage capacities to breakthrough this tradeoff. For example, faster elongation with denser timestamp incorporation could be applied to high-resolution, short-duration recordings, whereas slower elongation could be used to monitor long-term transcriptional activity over weeks or months, albeit with reduced temporal resolution. Additionally, future work could combine CytoTape with other cell activity reporter systems, such as the Fucci system^121^ to correlate transcriptional activity with cell cycle progression, and multi-round immunostaining techniques to visualize many more kinds of molecular tags, thus dramatically increasing the number of simultaneously recordable cellular signals, *i.e.*, the multiplexing capacity, along a single assembly. Expanding the recorder repertoire to include reporters for calcium^122, 123^ and kinase activities^124^ would further broaden CytoTape’s utility, while additional engineering efforts may be required to push the hours-scale temporal precision down to minutes- or even-seconds scales for these faster cellular activities. Because CytoTape supports multi-week recordings across broad spatial and temporal scales while maintaining single-cell resolution, integrating it with spatial omics platforms and AI-powered analysis pipelines for high-throughput readout and data analysis could enable comprehensive four-dimensional maps of cellular activities and states in cell populations, capturing both spatial and temporal context simultaneously. Beyond recording, intracellular architectures like CytoTape may have potential to serve as molecular scaffolds for a broad range of future applications, including synthetic cellular barcoding^125^, regulation of cellular behavior or enzymatic activity^126^, implementation of synthetic gene circuits^127^, construction of engineered molecular logic systems^128^, and development of human-designed interfaces between cells and external systems^16^.

## 4 Methods

### *ProtSSN*-based protein mutation prediction

*ProtSSN*^54^ formulates a zero-shot mutation effect prediction task to suggest favorable modifications on a template protein. It takes the sequence and structure of a template protein (1POK(E239Y) in our case) as input and uses a deep learning model to extract residue-wise representations. Based on these representations, it produces a predicted matrix 𝑌^#^ ∈ ℝ^𝑙×33^ for the template protein with each of 𝑙 residues being one of 33 tokens, including 20 regular residue types and other special tokens defined for training purposes. This matrix can be used to compute mutation fitness scores by comparing the differences between mutants and the template protein at the residue level, thus identifies top-ranked candidate mutants. In the following sections, we describe the model architecture and scoring procedure in detail.

The core of *ProtSSN* consists of a pre-trained BERT-style masked protein language model (PLM) that captures long-range residue dependencies along the sequence, followed by a learnable equivariant graph neural network (EGNN)^129^ that enhances local spatial interactions of residues (**Extended Data** Fig. 1a). For an arbitrary input protein, we first define it as an undirected graph 𝒢 = (𝒱, ℰ, 𝑊_𝑉_, 𝑊_𝐸_, 𝑋_𝑉_) using a *k*-nearest neighbor (kNN) algorithm. Each node 𝑣 ∈ 𝑉 corresponds to a residue. For each central node, undirected edges 𝑒 ∈ ℰ are added to up to *k* nearest neighboring residues within a 30 Å radius in Euclidean space. Node features 𝑊_𝑉_ are 1024-dimensional hidden embeddings extracted from the frozen ESM2-t33 model^130^ (checkpoint available at https://github.com/facebookresearch/esm). Following graph denoising neural networks^131^, edge features 𝑊_𝐸_ ∈ ℝ^|𝐸|×93^ capture pairwise relationships between connected residues based on inter-atomic distances, local backbone directions (N–C vectors), and relative sequence positions. Additionally, to maintain roto-translation equivariance and permutation invariance for node attributes during geometric message passing, 𝑋_𝑉_ stores the 3D coordinates of residues in Euclidean space.

The parameters of the EGNN layers are learned through a self-supervised denoising task. For each residue in the input sequence, a random perturbation is applied with probability 𝑝 using a replacement matrix Θ(⋅), defined by empirical observations. This perturbation updates each residue 𝑣 to *v̂* The frozen protein language model then encodes the perturbed residue sequence and generates the node features 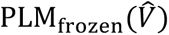, where 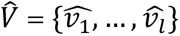 is the perturbed sequence. The EGNN layers further process the representation, enhancing local environmental information, and output 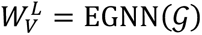. This is subsequently passed to the readout layers *ϕ*(·) to provide the recovered sequence 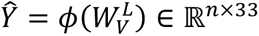. The training objective is to minimize Loss(𝑌, *Ŷ*), *i.e.*, the difference between the denoised and the ground-truth residue tokens.

Once the model is properly trained, it can be used to infer mutation effect scores for mutants by comparing with the template residue types 𝑉. In this case, the template sequence and structure are fed into the frozen PLM and trained EGNN layers (without additional perturbation, which is different from the training steps) to output *Ŷ.* Suppose the mutant of interest has mutated sites τ (|𝜏| ≥ 1). Its fitness score is computed as 𝐹_𝑥_ = ∑_{𝑡∈𝜏}_ log 𝑝(*Ŷ*_𝑡_) − log 𝑝(𝑣_𝑡_), where 𝑦_𝑡_ and 𝑣_𝑡_ represent the mutated and wild-type residues at the 𝑡th position, respectively. While the absolute value of these fitness scores does not have a direct physical meaning, when multiple mutants are under consideration, their corresponding fitness scores are effective references for ranking the preference of these mutations. In the case of recommending single-site mutations for 1POK(E239Y) in this study, we employed *ProtSSN* to score single-site saturation mutations and selected the top 2 mutations for further investigation and validation through wet lab experiments.

#### *CPDiffusion*-based protein sequence generation

*CPDiffusion*^53^ is a discrete diffusion probabilistic model designed for the protein inverse folding task, *i.e.*, recovering the residue sequences 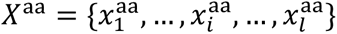 conditioned on a given protein backbone 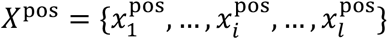. To meet specific application needs and improve generation quality, the model allows the incorporation of additional conditions during the generation process, such as fixing conserved residue positions, providing reference secondary structure sequences, or including homologous sequences in the training set. In addition, the generated sequences are further refined through a series of structure-based filtering strategies to enhance the success rate of novel protein designs.

To achieve this generative objective, *CPDiffusion* establishes a progressive diffusion process over residue types and trains a denoising network to gradually recover the input sequence distribution (𝑋^aa^|𝑋^pos^). The forward diffusion process adds noise step-by-step to one-hot encoded residue types based on a transition probability matrix (*e.g.*, the BLOSUM62 matrix) until time 𝑇, when 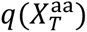 asymptotically converges to a uniform distribution and becomes independent of the initial state 𝑋^aa^. In the denoising phase, a multi-layer EGNN is trained to predict 𝑋^aa^ from the noisy sequence 𝑋_𝑡_ at time 𝑡, using the input protein backbone (represented as a *k*NN undirected graph) and additional conditions (defined in the node attributes, such as secondary structure information). During training, the network parameters 𝜃 in EGNN(⋅) are learned from a joint training set consisting of protein crystal structures in CATH 4.2^132^ and homologous sequences of 1POK (*i.e.*, *E. coli* IadA) along with their structures predicted by AlphaFold2 ^133^, with the objective of minimizing the cross-entropy loss between the predicted sequences 𝑝^(𝑋^aa^) = EGNN_𝜃_(𝑋_𝑡_, 𝐸, 𝑡) and the ground-truth sequences 𝑋 for each input protein. Optionally, a mask matrix ℳ can be defined to fix conservative sites. In this case, the predicted sequences by EGNN requires an additional step of adjustment 𝑝^(𝑋^aa^|𝑋_𝑡_) = ℳ ⊙ 𝑝(𝑋^aa^) + (1 − ℳ) ⊙ 𝑝^(𝑋^aa^|𝑋_𝑡_). We fixed tyrosine (Y) at the 293 position of the amino acid sequence during the generation process to keep 293Y in all the generated novel sequences, because E293Y has been reported to be critical for the linear assembly of 1POK(E293Y)^50^. During inference, for a given target protein backbone (*e.g.*, the crystal structure of 1POK, *i.e.*, *E. coli* IadA), AA sequences 𝑋_𝑡_ are sampled iteratively starting from a uniform sampling 𝑋_𝑇_ at time 𝑇 until time 𝑡 = 1 when reaching 𝑝(𝑋^aa^|𝑋_1_). Multiple sequences can then be generated by sampling from this distribution.

Following the above procedure, we used *CPDiffusion* to generate 100 sequences and predicted their structures using AlphaFold2. We then evaluated the predicted structures by comparing them to the original 1POK/IadA crystal structure in terms of RMSE, TM-score, and the residue-wise pLDDT trends of the generated sequences versus the wild-type template sequence predicted by AlphaFold2. Based on these criteria, we selected the optimal-performed 5 sequences for further experimental validation.

#### Molecular dynamics simulations

The structure of the protein monomer for simulations was predicted by AlphaFold3^52^ (**Fig. 1h and i**). The simulations were replicated three times. Protein and a large number of water molecules were filled in a virtual cubic box in the simulation. Chlorine counter ions were added to keep the system neutral in charge. The CHARMM36m force field was used for the complex and the CHARMM-modified TIP3P model was chosen for water. The simulations were carried out at 310 K. After the 4000-step energy-minimization procedure, the systems were heated and equilibrated for 100 ps in the NVT ensemble and 500 ps in the NPT ensemble. The 500-ns production simulations were carried out at 1 atm with the proper periodic boundary condition, and the integration step was set to 2 fs. The covalent bonds with hydrogen atoms were constrained by the LINCS algorithm. Lennard-Jones interactions were truncated at 12 Å with a force-switching function from 10 to 12 Å. The electrostatic interactions were calculated using the particle mesh Ewald method with a cutoff of 12 Å on an approximately 1 Å grid with a fourth-order spline. The temperature and pressure of the system are controlled by the velocity rescaling thermostat and the Parrinello-Rahman algorithm, respectively. All molecular dynamics simulations were performed using GROMACS 2020.4 packages.

#### Molecular cloning

The DNAs encoding the protein motifs used in this work were mammalian-codon optimized and synthesized by Epoch Life Science, and then cloned into mammalian expression backbones, pAAV-*UbC* (for constitutive expression), pAAV-c*-fos* (for expression driven by the c-*fos* promoter), pAAV-*Arc* (for expression driven by the SARE-ArcMin promoter), pAAV-*Egr1* (for expression driven by the *Egr1* promoter), pAAV-Npas4 (for expression driven by the *N*-RAM promoter), pAAV-Fos (for expression driven by the *F*-RAM promoter), pAAV-CREB (for 6x*CRE*-CMVMin-dependent expression) or pAAV-*HSPA1A* (for expression driven by the *HSPA1A* promoter) for DNA transfection in cultured neurons, glial cells, HEK293T clone 17 (referred to as HEK throughout this paper; ATCC CRL-11268) cells, and HeLa cells (ATCC CCL-2). See **Table S1** for sequences of the motifs, **Table S2** for all constructs of protein monomer designs, and **Tables S3-5** in **Supplementary information** for all designed signal constructs.

#### Animals and preparation of primary mouse hippocampal neuron-glia co-cultures

All procedures involving animals at the University of Michigan were conducted in accordance with the United States National Institutes of Health Guide for the Care and Use of Laboratory Animals, and were reviewed and approved by the University of Michigan Institutional Animal Care & Use Committee. Cultured hippocampal neurons with glial cells were prepared from neonatal (postnatal day 0 or 1) Swiss Webster mice (Taconic; both male and female mice were used) as previously described^38^. Briefly, the brains of ice-anesthetized neonatal mice were dissected out, and the hippocampal tissue was further dissected from the brains in an ice-cold dissection buffer (1 mM Kynurenic acid, 10 mM MgCl_2_, 35 mM D-glucose in HBSS). The hippocampal tissue was subsequently treated with papain for 10 min at 37 °C, followed by a 5-minute ovomucoid inhibition. The dissociated tissues were triturated, and a single-cell suspension was prepared using neuronal culture media (Minimum Essential Medium (MEM; Gibco) supplemented with 10 mM HEPES, 10% heat inactivated fetal bovine serum (HI FBS; Corning), 2% B27 supplement (Gibco), 2 mM L-Glutamine, 1 µM transferrin, 25 mM D-glucose; pH adjusted to 7.4). Cells were plated at a density of 40,000 cells per 100 µL on Matrigel (Corning)-coated coverslips (Carolina) and then incubated in a humidified cell culture incubator at 37 °C with 5% CO_2_. After the cells had settled and attached to the coverslips, 1 mL of neuronal culture media was added. When glial cells reached approximately 80% confluency (usually two days after plating), 1 mL neuronal culture media supplemented with 4 µM AraC was added to the existing media in each well.

#### CytoTape expression in cultured neurons and glial cells

Cultured neurons and glial cells were transfected at 7 days *in vitro* (DIV 7) for protein assembly design screening (**Fig. 1d**, **Extended Data Fig. 2b**), CytoTape with timestamps (**Fig. 3**), and CytoTape-based transcriptional recorders (**Fig. 4**, **Extended Data Fig. 5**). For recording the time course of c-*fos*, *Arc*, *Egr1*, Npas4, and CREB-promoter-driven expression (**Fig. 5** and **7**, **Extended Data Fig. 8**), cells were transfected at DIV 4. A commercial calcium phosphate transfection kit (Invitrogen) was used as previously described^16, 38^, with the following transfection parameters. For protein assembly design screening (**Fig. 1**, **Extended Data Fig. 2**), each well of a 24-well glass-bottom plate received 50 ng of the structural monomer plasmid and 1450 ng of the pUC19 plasmid (a “dummy” plasmid without a mammalian open reading frame to maintain the optimal mass ratio between DNA and calcium phosphate for the formation of co-precipitates for transfection). In the timestamped CytoTape expression experiment (**Fig. 3**), each well received 50 ng of the structural monomer plasmid (*UbC*-CytoTape), 5 ng of the timestamp monomer plasmid (*UbC*-CytoTape-HaloTag), and 1445 ng of pUC19 plasmid. For the CytoTape-based transcriptional recorder experiment (**Fig. 4**, **Extended Data Fig. 5**), each well was transfected with 50 ng of the structural monomer plasmid (*UbC*-CytoTape), 12.5 ng of CytoTape-HaloTag plasmid (*UbC*-CytoTape-HaloTag), and 1437.5 ng of pUC19 plasmid. In the single signal recording experiment (**Fig. 5**, **Extended Data Fig. 8**), each well received 300 ng of the structural monomer plasmid (*UbC*-CytoTape), 30 ng of the timestamp monomer plasmid (*UbC*-CytoTape-HaloTag), 75 ng of the signal monomer plasmid (c-*fos*, *Egr1*, *Arc*, or Npas4-CytoTape-V5), and 1095 ng of the pUC19 plasmid. For the two-signal recording experiment (**Fig. 5** and **7**), each well was transfected with 300 ng of the structural monomer plasmid (*UbC*-CytoTape), 30 ng of the timestamp monomer plasmid (*UbC*-CytoTape-HaloTag), 2×75 ng of the signal monomer plasmids (c-*fos*-CytoTape-V5 and *Arc*-CytoTape-OLLAS), and 1020 ng of pUC19 plasmid. After a 45-min incubation with DNA-calcium phosphate precipitates, cells were washed with a preformulated acidic buffer, HMEM (that is the MEM buffer supplemented with 15 mM HEPES and then adjusted to a final pH of 6.70−6.74 with acetic acid (Millipore Sigma) over the time course of 6 h) to remove residual precipitates. Cells were then placed back to the cell culture incubator.

#### CytoTape expression in HEK and HeLa cells

HEK and HeLa cells at a low passage number (*<* 10 passages) were plated at 40% confluence onto a 200-mm tissue culture dish. Dulbecco’s Modified Eagle’s Medium (DMEM; Gibco; containing high glucose, GlutaMAX supplement, and pyruvate) supplemented with 10% HI FBS (Corning) and penicillin/streptomycin (Gibco) was used as the HEK and HeLa cell culture media. Cells were then grown in a cell culture incubator at 37 ^◦^C with 5% CO_2_. After reaching 50-70% confluence, cells were transferred to 24-well glass-bottom plates via trypsin treatment. The 24-well glass-bottom plates were pre-treated with matrigel at room temperature for 30 min before cell plating. 24 h after cell plating, genes were delivered using the *Trans*IT-X2 Dynamic Delivery System kit (Mirus Bio).

For structural monomer expression experiment (**Fig. 2**), 100 ng structural monomer plasmid (*UbC*-CytoTape-HA) and 20 ng pUC19 plasmid were first diluted in 50 µL Opti-MEM medium (Gibco), followed by the addition of 0.9 µL *Trans*IT-X2 reagent. The mixture was incubated at room temperature for 25 min and then added to the single well. The cells were further incubated at 37 °C with CO_2_ for 24 h. For timestamped CytoTape expression experiment (**Fig. 3**, **Extended Data Fig. 4**), 100 ng structural monomer plasmid (*UbC*-CytoTape-HA), 100 ng timestamp monomer plasmid (*UbC*-CytoTape-HaloTag), and 190 ng pUC19 plasmid were first diluted in 50 µL Opti-MEM medium, followed by the addition of 0.9 µL *Trans*IT-X2 reagent. The mixture was incubated at room temperature for 25 min and then added to the single well. The cells were further incubated at 37 °C with CO_2_ for 24 h. For single-signal recording experiment (**Extended Data Fig. 7**), 100 ng structural monomer plasmid (*UbC*-CytoTapeHA), 10 ng timestamp monomer plasmid (*UbC*-CytoTape-HaloTag), 25 ng signal monomer plasmid (c-*fos*, CREB, or *HSPA1A*-CytoTape-V5), and 165 ng pUC19 plasmid were first diluted in 50 µL Opti-MEM medium, followed by the addition of 0.9 µL *Trans*IT-X2 reagent. The mixture was incubated at room temperature for 25 min and then added to the single well. The cells were further incubated at 37 °C with CO_2_ for 24 h. For two-signal recording experiment (**Fig. 6**), 100 ng structural monomer plasmid (*UbC*-CytoTape), 10 ng timestamp monomer plasmid (*UbC*-CytoTape-HaloTag), 2 × 25 ng signal monomer plasmids (CREB-CytoTape-HA and Fos-CytoTape-OLLAS), and 140 ng pUC19 plasmid were first diluted in 50 µL Opti-MEM medium, followed by the addition of 0.9 µL *Trans*IT-X2 reagent. The mixture was incubated at room temperature for 25 min and then added to the single well. The cells were further incubated at 37 °C with CO_2_ for 24 h. For five-signal recording experiments (**Fig. 5**), 180 ng structural monomer plasmid (*UbC*-CytoTape), 20 ng CytoTape-HaloTag plasmid (*UbC*-CytoTape-HaloTag), and 5 × 25 ng signal monomer plasmid (CREB-CytoTape-HA, *Egr1*-CytoTape-Etag, c-*fos*-CytoTape-OLLAS, *Arc*-CytoTape-V5, and Npas4-CytoTape-FLAG) were first diluted in 50 µL Opti-MEM medium, followed by the addition of 1.5 µL *Trans*IT-X2 reagent. The mixture was incubated at room temperature for 25 min and then added to the well. The cells were further incubated in the cell culture incubator at 37 °C with 5% CO_2_ for 24 h.

#### CytoTape timestamps labeling

CytoTape was labeled with cell-permeant HaloTag-ligand Janelia Fluor (JF) dyes (Promega) (**Fig. 3**, **5**, **6**, and **7**, **Extended Data Fig. 4**, 7, and **8**). JF_503_, JF_585_, and JF_635_ were used in this study for timestamps. The JF dyes were in lyophilized powder form and stored at -20 °C before use. The JF dye powder was first dissolved in 50 µL dimethyl sulfoxide (DMSO) and then diluted into 10 mL of HEK and HeLa cell culture media (for HEK and HeLa cells) or into 10 mL of neuronal culture media (for cultured neurons and glial cells) at a final concentration of 0.1 µM. The dyed medium was filtered using a sterile 0.22 µm syringe filter before being added to the culture. It was then used to replace the original media in cell cultures at designated time points. During the dye-switching process, the medium containing the original dye was fully removed, followed by thorough washes of the cell cultures five times with 37 °C DMEM for HEK and HeLa cells, and MEM for cultured neurons and glial cells. Finally, the cells in each well of the 24-well plate were cultured in 500 µL of fresh HEK and HeLa cell culture media (for HEK and HeLa cells) or 1 mL neuronal culture media (for cultured neurons and glial cells) supplemented with a new JF dye at 0.1 µM final concentration in the cell culture incubator at 37 °C with 5% CO_2_.

#### Chemical stimulation of cultured neurons

For KCl stimulation (**Fig. 4, 5, and 7**, **Extended Data Fig. 8**), a KCl stock buffer was prepared, containing 170 mM KCl, 2 mM CaCl_2_, 1 mM MgCl_2_, and 10 mM HEPES. Then, a KCl depolarization medium was prepared by mixing the KCl stock buffer with fresh neuronal culture medium, ensuring that the final concentration of K^+^ after mixing was 55 mM or 30 mM (accounting for the K^+^ already present in the fresh neuronal culture medium). The original culture medium from neuron cultures was transferred into a fresh 24-well plastic-bottom plate, where the medium from different neuron cultures was stored in separate wells and kept in the neuronal incubator until the end of the KCl-induced depolarization treatment. Then, 500 µL of KCl depolarization medium was added to each well containing neuron cultures. The neuron cultures were placed back into the incubator and incubated for 10 min, 30 min, 1 h, 2 h, or 3 h. Finally, the KCl depolarization medium was removed, and the original neuronal culture medium was transferred back into the corresponding wells. The neuron cultures were then returned to the cell culture incubator.

For forskolin (FSK) stimulation (**Fig. 4, 5** and **7**), FSK powder was dissolved in DMSO and the final concentration of it is 5 mM. Then, a FSK stimulation medium was prepared by mixing the 5 mM FSK solution with a fresh neuronal culture medium, ensuring that the final concentration of FSK after mixing was 5 µM, 10 µM, 20 µM, and 25 µM. The FSK stimulation medium was filtered using a sterile 0.22 µm syringe filter (VWR, Avantor) before being added to neurons. The original culture medium from neuron cultures was transferred into a fresh 24-well plastic-bottom plate, where the medium from different neuron cultures was stored in separate wells and kept in the neuronal incubator until the end of the FSK stimulation. Then, 500 µL of FSK stimulation medium was added to each well containing neuron cultures. The neuron cultures were placed back into the incubator and incubated for 1 h. Finally, the FSK stimulation medium was removed, and the original neuronal culture medium was transferred back into the corresponding wells. The neuron cultures were then returned to the cell culture incubator.

#### Chemical stimulation of cultured mouse hippocampal glial cells

For KCl stimulation (**Extended Data Fig. 6**), a KCl stock buffer was prepared, containing 170 mM KCl, 2 mM CaCl_2_, 1 mM MgCl_2_, and 10 mM HEPES. Then, a KCl depolarization medium was prepared by mixing the KCl stock buffer with fresh neuronal culture medium, ensuring that the final concentration of K^+^ after mixing was 55 mM (accounting for the K^+^ already present in the fresh neuronal culture medium). The original culture medium was transferred into a fresh 24-well plastic-bottom plate, where the medium from different cultures was stored in separate wells and kept in the neuronal incubator until the end of the KCl-induced depolarization treatment. Then, 500 µL of KCl depolarization medium was added to each well containing cultures. The cultures were placed back into the incubator and incubated for 1 h. Finally, the KCl depolarization medium was removed, and the original culture medium was transferred back into the corresponding wells. The cultures were then returned to the cell culture incubator.

#### Chemical stimulation of HEK cells

For FSK stimulation (**Fig. 5** and **6**, **Extended Data Fig. 7**), FSK powder was dissolved in DMSO to a final concentration of 5 mM. Then, an FSK stimulation medium was prepared by mixing the 5 mM FSK solution with fresh HEK cell culture medium, ensuring that the final concentration of FSK after mixing was 10 µM, 20 µM, and 50 µM. The FSK stimulation medium was filtered using a sterile syringe filter before being added to HEK cells. The original culture medium from HEK cell cultures was removed. Then, 500 µL of FSK stimulation medium was added to each well and incubated for 1 h. Finally, the FSK stimulation medium was removed, and fresh culture medium was transferred into the corresponding wells. The HEK cell cultures were then returned to the cell culture incubator.

#### Heat treatments of HEK and HeLa cells

HEK (**Extended Data Fig. 7**) and HeLa cells (**Extended Data Fig. 2**) were cultured in cell culture media in a cell culture incubator set to 5% CO_2_ at 37 °C. Cells were plated in 24-well glass-bottom plates to achieve 90-100% confluence prior to the heat shock treatment. Then, the plates containing the cells were transferred into a mini incubator preheated to 42 °C and incubated there for 1 h. After the heat shock, the plates were returned to the original 37 °C cell culture incubator for recovery.

#### Antibodies and Nissl stain

Primary antibodies (1:500 for immunofluorescence of cultured cells): anti-Etag (Invitrogen MA5-38276), anti-V5 (Invitrogen R960-25), antiOLLAS (Invitrogen MA5-16125), anti-FLAG (Invitrogen 740001), and anti-HA (Cell Signaling Technology 11846S; Invitrogen PA5-33243).

Fluorescent secondary antibodies (1:500 for immunofluorescence of cultured cells): Mouse IgG1 ATTO 425 conjugated antibody pre-absorbed (Rockland), Goat anti-Mouse IgG2a Alexa Fluor 488 (Invitrogen A-21131), Goat anti-Rat IgG(H+L) Alexa Fluor 546 (Invitrogen A-11081), Goat anti-Rabbit IgG(H+L) Alexa Fluor 594 (Invitrogen A-11012), Goat anti-Chicken IgY(H+L) Alexa Fluor Plus 647 (Invitrogen A32933), Goat anti-Mouse IgG2a Alexa Fluor 546 (Invitrogen A21133), and Goat anti-Rabbit IgG(H+L), Alexa Fluor Plus 488 (Invitrogen A-11008).

Nissl stain: NeuroTrace 435/455 Blue Fluorescent Nissl Stain (Invitrogen N21479, 1:1000 for immunofluorescence of cultured cells). Additional details of primary antibodies, secondary antibodies, and Nissl used in this study are listed in **Supplementary Table 1**.

#### Immunofluorescence of cultured cells

Cells (neurons, glial, HEK, and HeLa cells) were fixed in TissuePrep-buffered 10% formalin (Fisherbrand) at room temperature (RT) for 10 min, followed by three 5-minute washes in 1×PBS at RT. Blocking was performed in MAXBlock blocking medium (Active Motif) supplemented with 0.1% Triton X-100 and 100 mM glycine for 20 min at RT, followed by three additional 5-min washes in MAXwash washing medium (Active Motif) at RT. Cells were then incubated with primary antibodies diluted in MAXbind staining medium (Active Motif) for 1 h at RT. Afterward, cells underwent three 5-min washes in MAXwash washing medium at RT. Secondary antibodies were applied in MAXbind staining medium and incubated for 1 h at RT. Cells were then washed three times with MAXwash washing medium for 5 min each at RT. Finally, cells were incubated with NeuroTrace 435/455 Blue Fluorescent Nissl Stain (Invitrogen) for 10 min and stored in 1×PBS at 4 °C until imaging.

#### Fluorescence microscopy of immunostained cells

Fluorescence microscopy was performed using a spinning disk confocal microscope (Yokogawa CSU-W1 Confocal Scanner Unit on a Nikon Eclipse Ti2 inverted microscope) equipped with a 40X 1.15 numerical aperture water immersion objective (Nikon MRD77410), a 10X objective, a Hamamatsu ORCA-Fusion BT sCMOS camera controlled by the NIS-Elements AR software, and laser/filter sets for 405 nm, 488 nm, 561 nm, 594 nm, and 640 nm optical channels. Multi-channel volumetric imaging was conducted at 0.4 µM per Z-step for each field of view under the 40X objective. Imaging parameters remained consistent across all samples within each experimental set.

#### Photobleaching of Janelia Fluor dyes

For the experiments described in **Fig. 5, 6,** and **7**, once imaging the Janelia Fluor (JF) dyes in cell samples with confocal microscopy, we performed LED exposures to completely photobleach the fluorescence signals and free up the optical channels for subsequent immunofluorescence staining. Samples in 24-well plates were placed approximately 30 cm away from a broadband white LED light source (ThorLabs, MNWHLP1) to be uniformly exposed to the LED illumination. We checked the fluorescence of JF dyes in the samples after every 10 min of LED exposure and continued such exposures until the fluorescence of JF dyes dropped below the detection limit of the confocal microscope. The total LED exposure duration typically ranged from 10 to 60 min, depending on the initial fluorescence of JF dyes in the samples before LED exposure.

#### Software for image analysis

Image analysis was performed in ImageJ (National Institutes of Health) and Python.

#### Segmentation and geometrical analysis of fibers in images

To capture precise information from CytoTape and improve data consistency, a computational analysis pipeline was developed in Python to extract information from raw 3D confocal microscopy images. The pipeline included several key steps: image deconvolution (in ImageJ. The rest of the steps were performed in Python); CytoTape fiber segmentation with the adaptive threshold strategy; centerline detection; fiber morphology verification, which involved removing intertangled protein fibers which has been recognized by algorithm as a whole (would result in information ambiguity) and noise-like short objects (mainly caused by excessive monomer nucleation events which not growing into a long fiber); and extraction of object properties, such as fluorescent intensity along the centerline, fiber length, thickness, curvature, and spatial locations in the 3D space. The pipeline’s reliability was validated using ground truth datasets from human annotation. With this pipeline, we obtained the fluorescent intensity profiles along the fibers for this work.

#### Readout information from intensity profiles

Fluorescence intensity profiles along the fiber were extracted from images as previously described^38^.

#### Time recovery with HaloTag-based timestamp

For each fiber, we divided it into two segments based on an optimal split point as previously described^38^. This point was determined by iterating through hypothetical split locations within a window spanning 20% of the total fiber length around its geometrical center point and selecting the one that maximized the Pearson correlation between the left and right halves. We reasoned that this optimal split best balances the trade-off between the inherent symmetry of CytoTape elongation and the variability in protein assembly introduced by the complex cellular environment.

For each half fiber (representing a complete time axis), we plotted the fluorescent intensities from different HaloTag-based timestamp channels, and manually identified critical points for dye color switching that represented real-time events along the fiber, serving as a bridge between fiber growth and real-time events. These critical points included the onset rise point of the second dye color relative to the first dye color for the first timestamp (reflecting when the second dye was actually added), the onset rise point of the next dye color (if present) relative to the second color for the second timestamp (reflecting when the subsequent dye was actually added), and the fiber endpoint (reflecting the actual cell fixation time). The normalized position along the fiber (dividing the distance from the initial point by its total length) were also calculated for each point, and we termed it as a fraction of fiber length.

Time axis recovery was achieved by fitting a polynomial curve between the the actual time points when dyes were added or cells were fixed (*e.g.*, Day 2, Day 5.5, and Day 11) and their corresponding fraction of fiber length values (*e.g.*, 0.5, 0.7, and 1). If there was only one dye switch, resulting in two timestamps (the dye switch and the cell fixation time corresponding to the fiber termini), a linear polynomial (first-degree) fitting is performed. If there were two dye switches, resulting in three timestamps (dye switch 1, dye switch 2, and the cell fixation time), a quadratic polynomial (second-degree) fitting or power fitting is used to best characterize the fiber growth kinetics, which has been experimentally characterized in **Fig. 2b**, right panel. After curve fitting, the resulting curve served as a transfer function between the spatial axis along the fiber and the real-world time axis. Fluorescent intensities along the fibers from the signal monomer channel(s) were initially plotted along each of the half fibers with respect to the spatial axis representing the fraction of the fiber length. This fractional fiber position is then converted into the real-world time axis using the fitted curve as the transfer function, thereby establishing the time axis for the recorded signal(s).

#### Statistical analysis

Statistical analysis was performed in Prism (GraphPad). All statistical tests were two-sided. Details of the statistical analysis are described in the figure captions and full statistical tables are in **Supplementary Table 2**.

## Supporting information

Statistical Analysis

## Supplementary files

**Supplementary information** includes the amino acid sequences of protein motifs, constructs of structural monomers, and constructs of signal monomers. **Supplementary file 1** includes the data presented in this study. **Supplementary file 2** includes the code for analysis. **Supplementary Table 1** includes reagents and resources used in this study. **Supplementary Table 2** includes the details of statistical analysis.

## Acknowledgments

We thank Dawen Cai, Wenjing Wang, William Joesten, Michael Widener, Cong Ma, Junfeng Yang, Yang Tan, Jeffrey Dudley, and Dylan Cable for discussion. L.Z. would like to thank the Michigan Neuroscience Institute Postdoctoral Advancement Program and the eLife Ambassadors Program. C.L. is supported by the National Institutes of Health (NIH) Director’s New Innovator Award (1DP2MH140133), the Chan Zuckerberg Initiative Collaborative Pairs Pilot Project Award, the Glenn Foundation for Medical Research and American Federation for Aging Research Grant Award for Junior Faculty, and the Whitehall Foundation.

## Declarations

### Conflict of interest

L.Z. and C.L. declare that they applied for a US provisional patent application (application number: 63/801,605) based on the work presented in this paper.

### Data availability

The plasmids and the corresponding sequence maps of working CytoTape constructs reported in this paper will be available at Addgene (plasmid IDs 239423-239430 and 239616) upon completion of the deposition process. The sequences of the CytoTape structural monomer, timestamp monomer, and all signal monomers will be available at GenBank upon completion of the deposition process.

### Code availability

The *ProtSSN* model for protein mutation prediction can be found at https://github.com/tyang816/ProtSSN. The *CPDiffusion* model for protein sequence generation prediction can be found at https://github.com/bzho3923/ CPDiffusion.

### Author contribution

L.Z. and C.L. conceived the concept of CytoTape and made high level plans for this project. B.Z. and L.Z. performed protein site mutation, protein sequence generation, and *in silico* protein characterization with the help from P.L. L.Z. performed rational design. L.Z. designed timestamps, transcriptional recorders, and multiplexing strategy with the help from D.S., B.A., E.S.B., and C.L. L.Z. prepared HEK293T and HeLa cells with the help from E.K., J.L., and D.S. J.L. prepared cultured neurons and glial cells. L.Z. performed the recording experiments. Y.Y. and L.Z. analyzed and interpreted the data with the help from D.W., C.L., B.K., and D.J.C. S.P. helped with the heat shock experiment. L.Z., Y.Y., B.Z., and C.L. wrote the manuscript. C.L. supervised the project.

---

**Extended Data Fig. 1.**
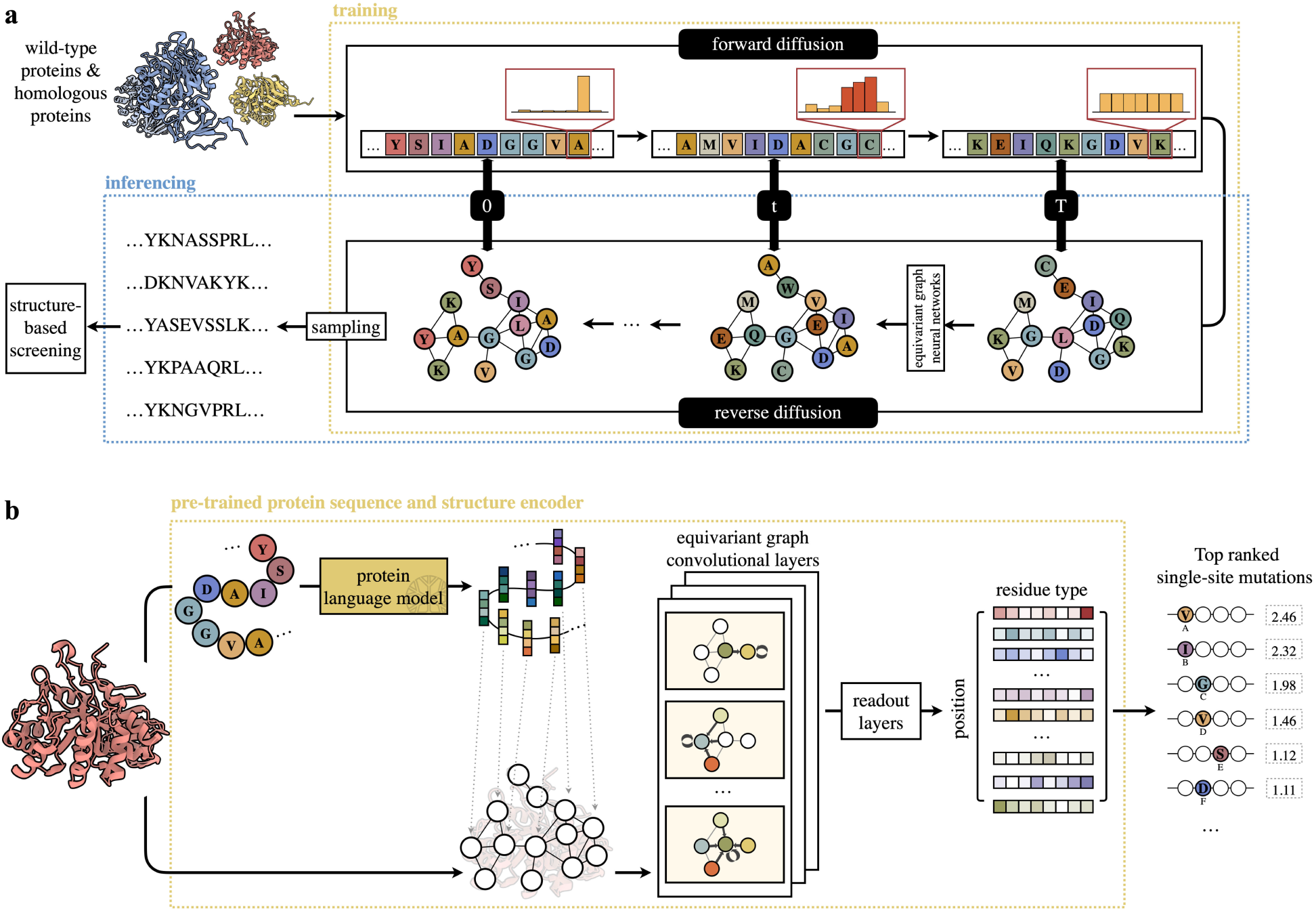
Workflow of *CPDiffusion* and *ProtSSN* for designing CytoTape monomer. (**a**) *CPDiffusion* comprises a forward diffusion process and a reverse generative process. The training phase (yellow box) includes both forward and reverse diffusion over *T* steps, where the denoising process is parameterized by trainable EGNNs that iteratively update the residue-type distributions. During inference (blue box), the model starts from a uniformly distributed residue sequence and applies the learned EGNN denoising network in reverse diffusion steps to progressively recover the residue-type distributions, conditioned on the input protein backbone. The generated novel sequences can then be sampled from this distribution and further screened using structure-based criteria to identify candidates for experimental validation. (**b**) *ProtSSN* scores mutation effects based on a wild-type template protein using pre-trained sequence and structure co-embedding modules (yellow box). With a frozen protein language model and a trainable roto-translation equivariant graph neural network, *ProtSSN* outputs a residue-wise score matrix across all residue types. This matrix is used to compute the fitness scores for mutant candidates, which can then be used to rank the most desirable mutations.

**Extended Data Fig. 2.**
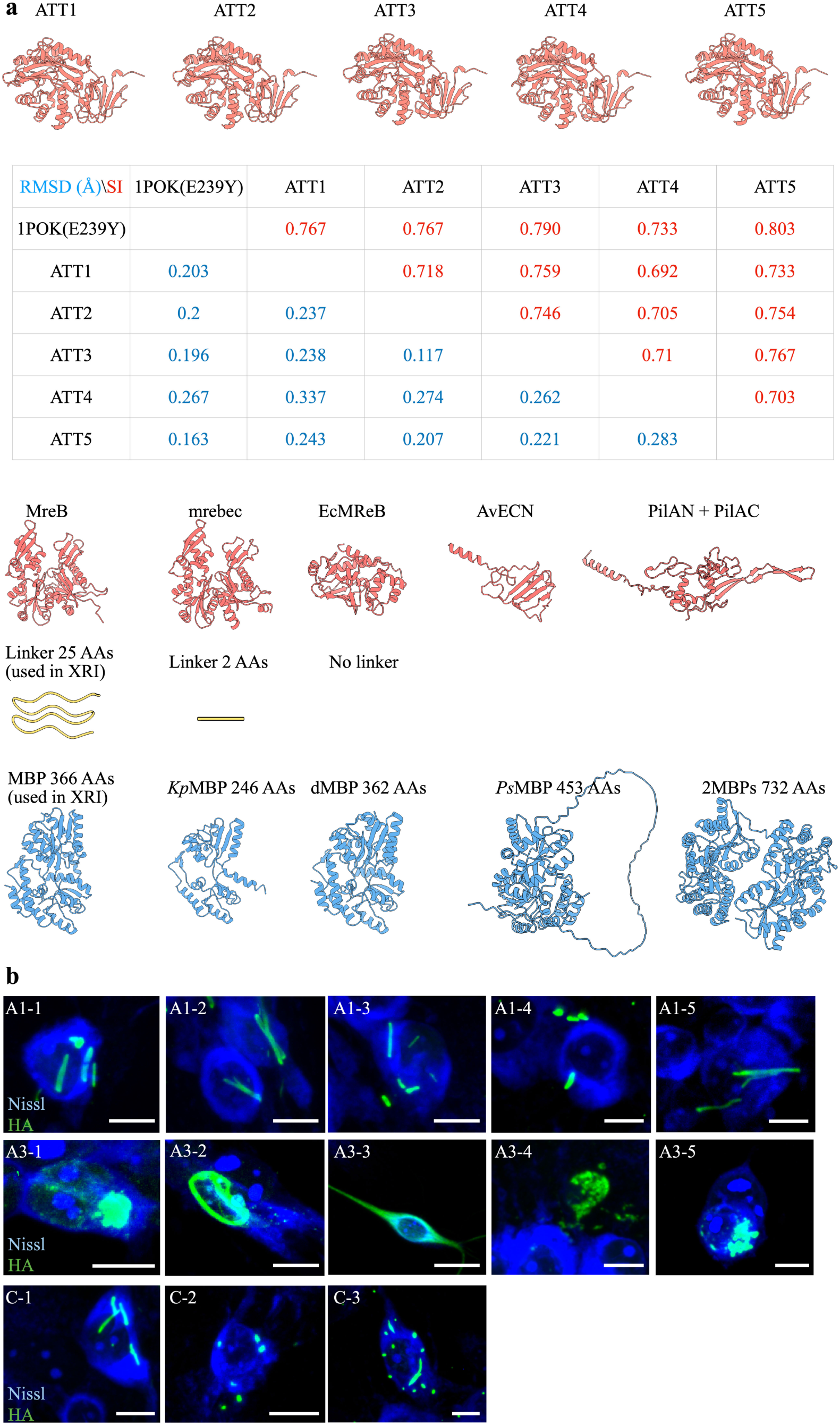
Protein assembly design screening and characterization in cultured neurons. (**a**) Atomic structures used in monomer designs. First row, five artificial scaffolds (artificial ticker tapes (ATTs); used in designs A1-1 to A1-5) generated by *CPDiffusion* for replacing 1POK(E239Y). The table below shows the sequence identity (SI) and root mean square deviation (RMSD) between each pair of scaffolds. The *CPDiffusion*-generated structures share a similar conformation with 1POK(E239Y), but vary in the amino acid sequences. Second row, natural scaffold for replacing 1POK(E239Y) (used in designs A3-1 to A3-5). Third row, linker length optimization (used in designs B-1 and C-4); fourth row, MBP homologs identified via BLAST search and their engineered variants. Note that the 2MBPs shown in the third row represent a direct fusion of two MBPs (366 AAs) (used in designs C-1 to C-5). All the atomic structures are predicted by AlphaFold3. (**b**) Confocal images of cultured mouse hippocampal neurons expressing design variants fused to the HA tag, and after fixation on day 7, Nissl staining, and immunostaining against the HA tag. The constructs transfected into neurons are listed in **Supplementary information**. Scale bars, 10 µm.

**Extended Data Fig. 3.**
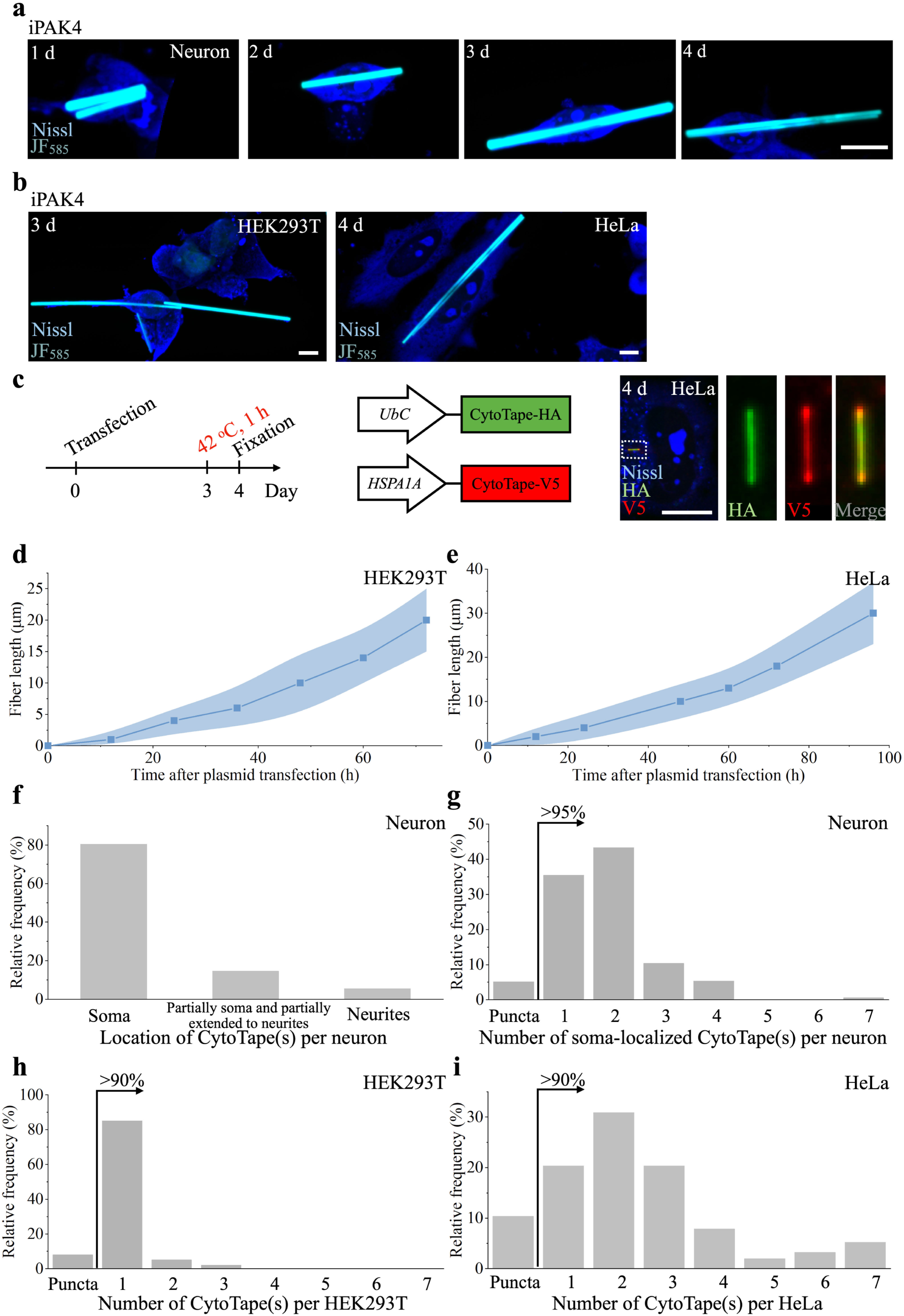
Analysis of iPAK4 and CytoTape formation in cultured neurons, HEK, and HeLa cells. Representative confocal images of (**a**) cultured mouse hippocampal neurons, as well as (**b**) HEK and HeLa cells expressing iPAK4 with iPAK4-HaloTag. Cells were labeled with JF_585_ dye added to the culture, fixed at different time points, and subsequently stained with Nissl. Scale bars, 10 µm. (**c**) Left panel, the *HSPA1A*-promoter-driven expression experiment timeline. Middle panel, the constructs transfected into HeLa cells. Right panel, confocal image of HeLa cell expressing CytoTape-based constructs, taken after fixation on day 4, Nissl staining and immunostaining against the HA tag and the V5 tag. Scale bar, 10 µm. The three rows of rectangular panels on the right are enlarged views of the regions marked by white rectangles in the left column of square panels. The growth kinetics of CytoTape in (**d**) HEK cell and (**e**) HeLa cell. 20 CytoTapes from two cell cultures were analyzed for length calculations. Blue boundary, s.d. (**f**) Histogram of the number of CytoTapes located in the neuron soma, partially soma and partially extended to neurites, and neurites (n = 50 neurons from five cultures). (**g**) Histogram of the number of soma-localized CytoTapes per neuron (n = 47 neurons from four cultures). ‘*>* 95%’ with an arrow, *>* 95% of the neurons have CytoTape(s) rather than puncta. Histogram of the number of CytoTapes per (**h**) HEK cell or (**i**) HeLa cell (n = 69 HEK cells from six cultures, n = 40 HeLa cells from two cultures). ‘*>* 90%’ with an arrow, *>* 90% of the cells have CytoTape(s) rather than puncta.

**Extended Data Fig. 4.**
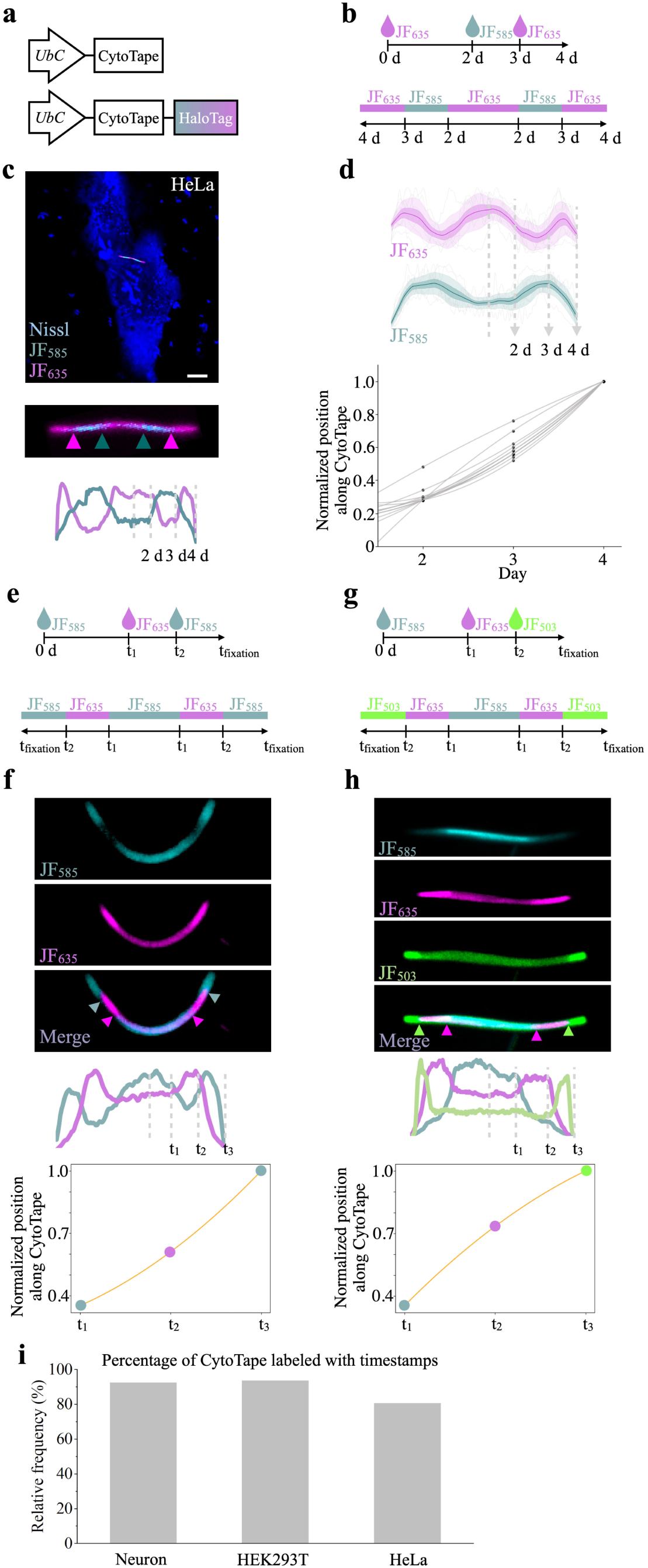
Development of timestamps for CytoTape via sequentially labeled dyes in HeLa cells and analysis of timestamp with different dye switches. (**a**) Schematic of the constructs transfected into HeLa cells. (**b**) Top panel, time points of JF_585_ and JF_635_ addition and fixation. Bottom panel, expected dye distribution along the CytoTape. The JF_585_ and JF_635_ dyes were used for labeling time within the CytoTape. (**c**) A representative confocal image of CytoTapes with timestamps in HeLa cells, which is taken after fixation on day 4 and Nissl staining. The middle row and bottom row show the CytoTape with timestamps and fluorescence line profiles, respectively. Scale bar, 10 µm. (**d**) Top panel, statistical analysis of fluorescence line profiles from the experiments described in (**b**) n = 8 CytoTapes from 8 HeLa cells from two cultures Bottom panel, interpolation between the spatial axis along fiber and the time axis using the timestamps from top panel. Time axis recovery via B-spline curve fitting of timestamps and fixation point. Each raw trace was normalized to its peak to show relative changes before averaging. Thick centerline, mean; darker boundary in the close vicinity of the thick centerline, s.e.m.; lighter boundary, s.d.; lighter thin lines, data from individual CytoTapes. (**e**) and (**g**) Time points of dye addition and fixation and expected dye distribution along the CytoTape. (**e**) JF_585_ and JF_635_ and (**g**) JF_585_, JF_635_, and JF_503_ were used for labeling time within the CytoTape. Images of timestamps with (**f**) two dye switches and (**h**) three dye switches. The middle and bottom rows show fluorescence line profiles and fitted curves, respectively. The details of the interpolation between the spatial axis along fiber and the time axis using timestamps are described in **Methods**. (**i**) The success rate of CytoTapes having timestamps in neuron, HEK, and HeLa cells (n = 40 CytoTapes from 40 neurons from five cultures; n = 50 CytoTapes from 50 HEK cells from three cultures; n = 30 CytoTapes from 30 HeLa cells from two cultures).

**Extended Data Fig. 5.**
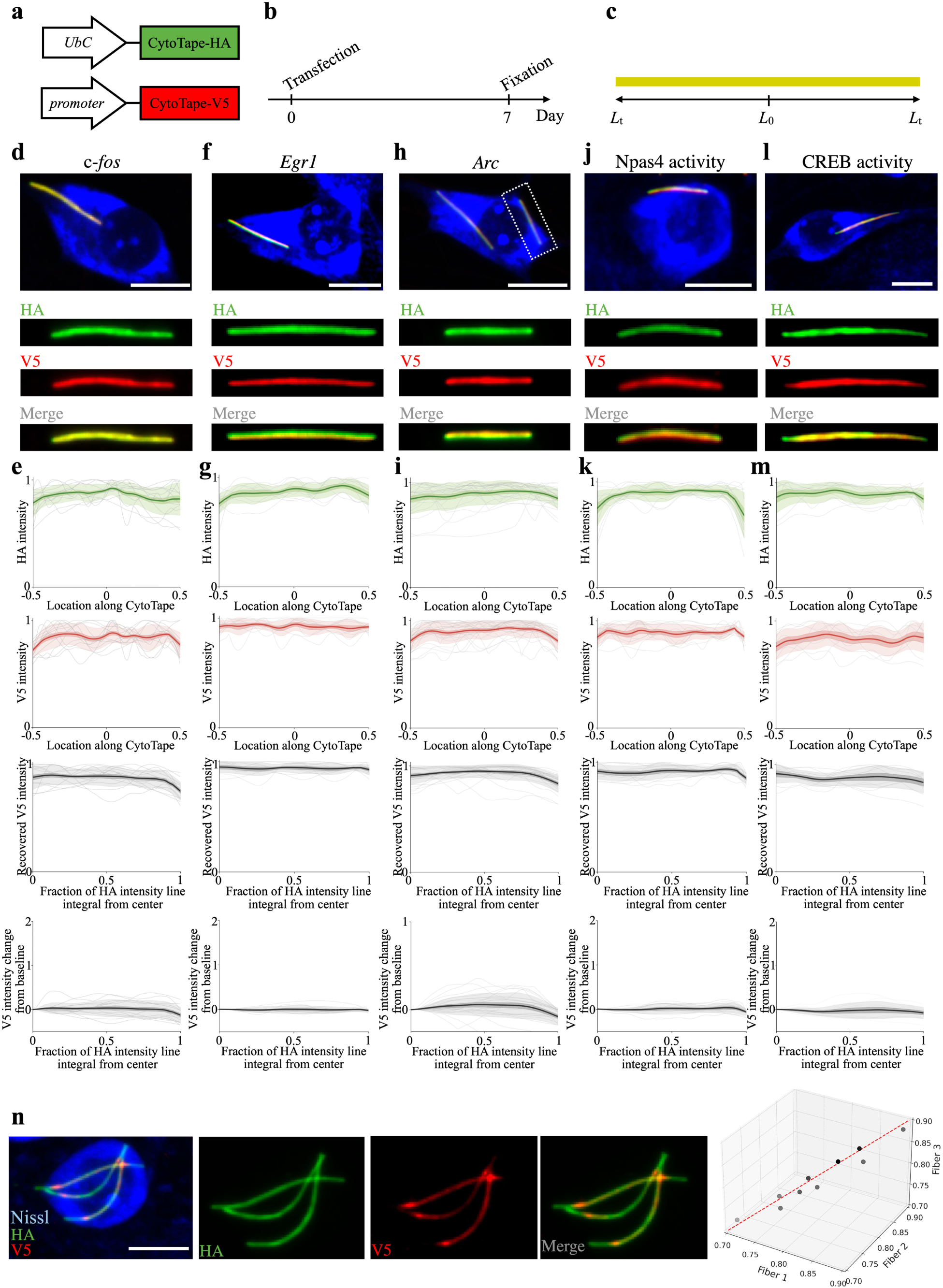
Control groups of transcriptional recorders for tracking c-*fos*-, *Arc*-, *Egr1*-, Npas4-, and CREB-promoter-driven expression with CytoTape. (**a**) Schematic of the constructs transfected into neuron cultures and (**b**) the experimental timeline. (**c**) Expected HA tag and V5 tag distribution along the CytoTape. (**d**), (**f**), (**h**), (**j**), and (**l**) Representative images of cultured neurons expressing the constructs shown in (**a**). Images were captured after fixation, Nissl staining, and immunostaining for HA and V5 tags. Enlarged views of the CytoTape in the top-row panels are shown in the three rows of rectangular panels below. Scale bars, 10 µm. (**e**), (**g**), (**i**), (**k**), and (**m**) Profiles of HA and V5 signal intensities along CytoTape, based on the experiment in (**b**)). First row, HA intensity profile; second row, V5 intensity profile; third row, recovered V5 signal, calculated from the intensity profiles, plotted as a function of the fraction of the HA intensity line integral; fourth row, V5 signal relative to baseline (calculated as the ratio of V5 signal to the center V5 signal) plotted as a function of the fraction of the HA intensity line integral. In the first three rows, raw traces were normalized to their peaks to highlight relative changes before averaging. Data were from c-*fos*, n = 12 CytoTapes from 12 neurons from two cultures; *Egr1*, n = 13 CytoTapes from 13 neurons from two cultures; *Arc*, n = 24 CytoTapes from 24 neurons from three cultures; Npas4 activity, n = 11 CytoTapes from 11 neurons from two cultures; CREB activity, n = 9 CytoTapes from 9 neurons from two cultures. Thick centerline, mean; darker boundary near the centerline, s.e.m.; lighter boundary, s.d.; thin light lines, individual CytoTape data. (**n**) A confocal image of a neuron showing three fibers. The experimental conditions are the same as in (**d**). The right panel shows the V5 signal peak position plotted against the HA signal. Each gray sphere represents a neuron (n = 10). The red dashed line represents x = y = z. Scale bar, 10 µm.

**Extended Data Fig. 6.**
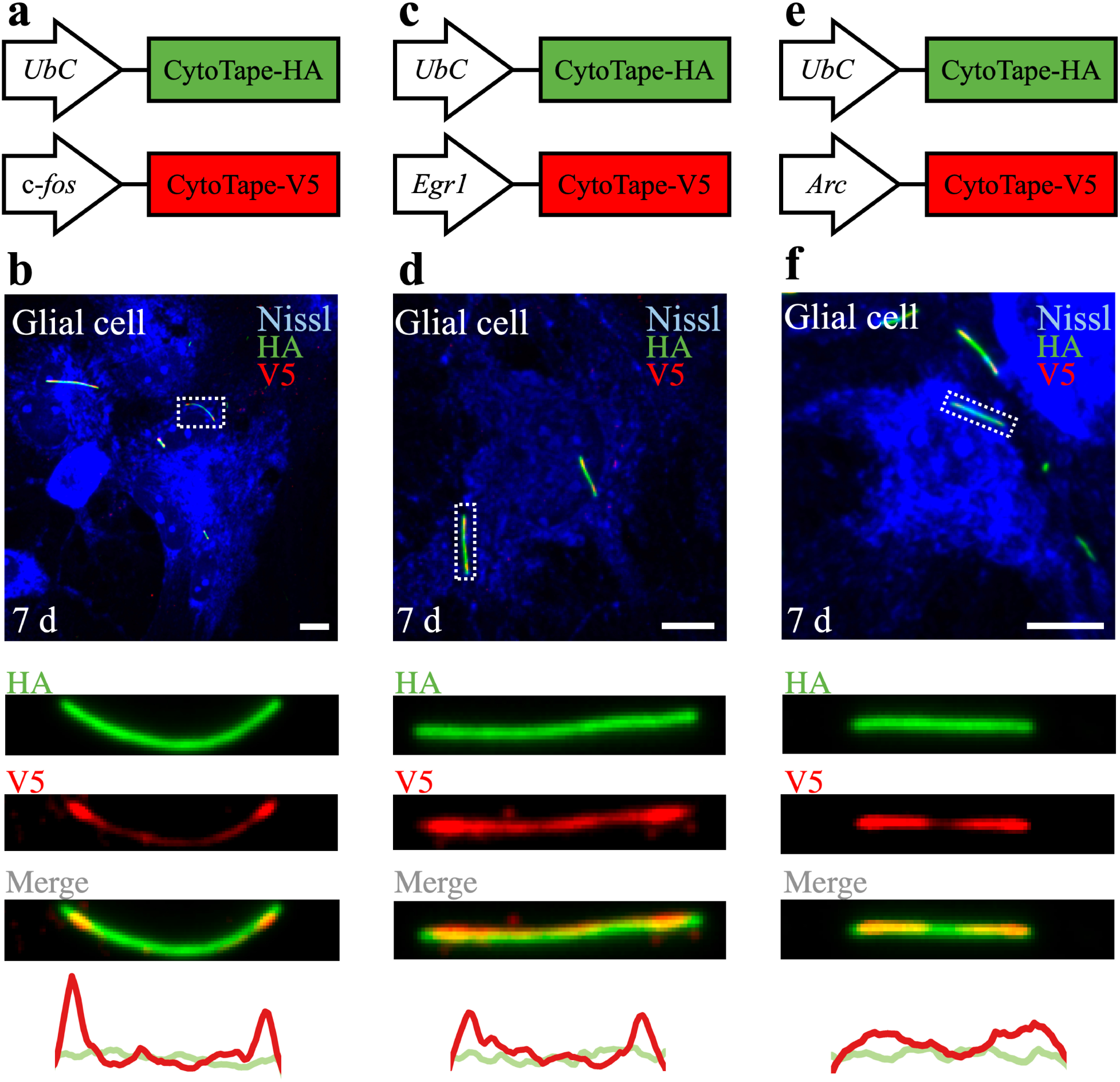
Waveform analysis of c-*fos*-, *Egr1*-, and *Arc*-promoter-driven expression with CytoTape in cultured mouse hippocampal glial cells. (a), (c), and. (**e**) Schematic of the constructs transfected into cultured mouse hippocampal glial cells. CytoTape-HA refers to CytoTape fused to the HA tag, and CytoTape-V5 refers to CytoTape fused to the V5 tag. (**b**), (**d**), and (**f**) Representative images of cultured mouse hippocampal glial cells expressing the constructs shown in (**a**), (**c**), and (**e**). Images were captured after fixation, Nissl staining, and immunostaining for HA and V5 tags. Glial cells were stimulated with 55 mM KCl for 1 h and fixed on day 7. Enlarged views of the white-marked regions in the top-row panels are shown in the three rows of rectangular panels below. The bottom row shows the fluorescence line profiles of HA and V5 tags. Scale bars, 10 µm.

**Extended Data Fig. 7.**
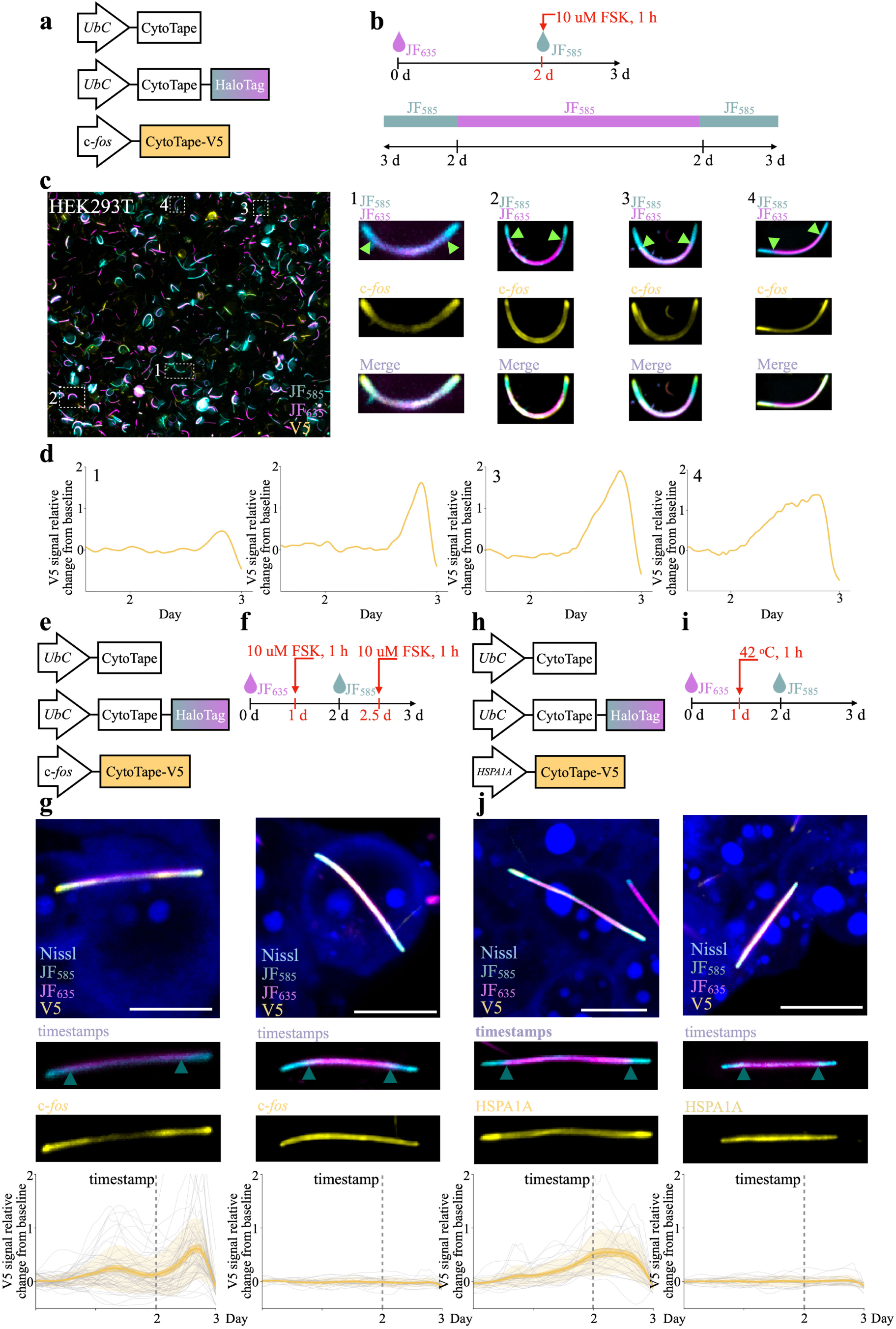
Recording c-*fos*- and *HSPA1A*-promoter-driven expression histories with CytoTape in HEK cells. (**a**) Schematic of the constructs transfected into HEK cells. (**b**) Top panel, time points of JF_585_ and JF_635_ dyes addition and stimulation (10 µM FSK for 1 h). Bottom panel, expected dye distribution along the protein fiber. The JF_585_ and JF_635_ dyes were added to the HEK cell culture on day 0 and day 2, and fixation was performed on day 3. (**c**) Left panel, low magnification images of CytoTape in HEK cells, which is taken after fixation on day 3, and Nissl staining and immunostaining against the V5 tag. Scale bar, 100 µm. Right panel, enlarged confocal images of CytoTapes indicated in left panel. Arrows indicate the positions of dye switches within the protein fiber. (**d**) c-*fos* signal (from (**c**)) relative change from baseline plotted against recovered time (interpolated with timestamps and fixation time) after calcium phosphate transfection. (**e**) and (**h**) Schematic of the constructs transfected into HEK cells and the experimental timeline. The JF_585_ and JF_635_ dyes were added to the HEK cell culture on day 0 and day 2, and fixation was performed on day 3. (**g**) and (**j**) Images of HEK cells expressing the constructs shown in (**e**) and (**h**) after stimulation with FSK for 1 h at different time points (left panels) and without stimulation (right panels). Images were captured after fixation, Nissl staining, and immunostaining for V5 tag. Enlarged views of the CytoTape in the top-row panels are shown in the two rows of rectangular panels below. The second-row and third-row panels show the timestamps and activity-dependent-promoter-driven expression, respectively. The third-row panel shows statistical analysis of V5 signal relative change from baseline plotted against recovered time after calcium phosphate transfection. Stimulation group, data are from n = 10 CytoTapes for c-*fos* from 10 HEK cells from two cultures; n = 13 CytoTapes for *HSPA1A* from 13 HEK cells from two cultures. Control group, data are from n = 10 CytoTapes for c-*fos* from 10 HEK cells from two cultures; n = 10 CytoTapes for *HSPA1A* from 10 HEK cells from two cultures. The dashed gray lines indicate the timestamps, marking the points after which the recovered time is accurate. Thick centerline, mean; darker boundary near the centerline, s.e.m.; lighter boundary, s.d.; thin light lines, individual CytoTape data. Scale bars, 10 µm.

**Extended Data Fig. 8.**
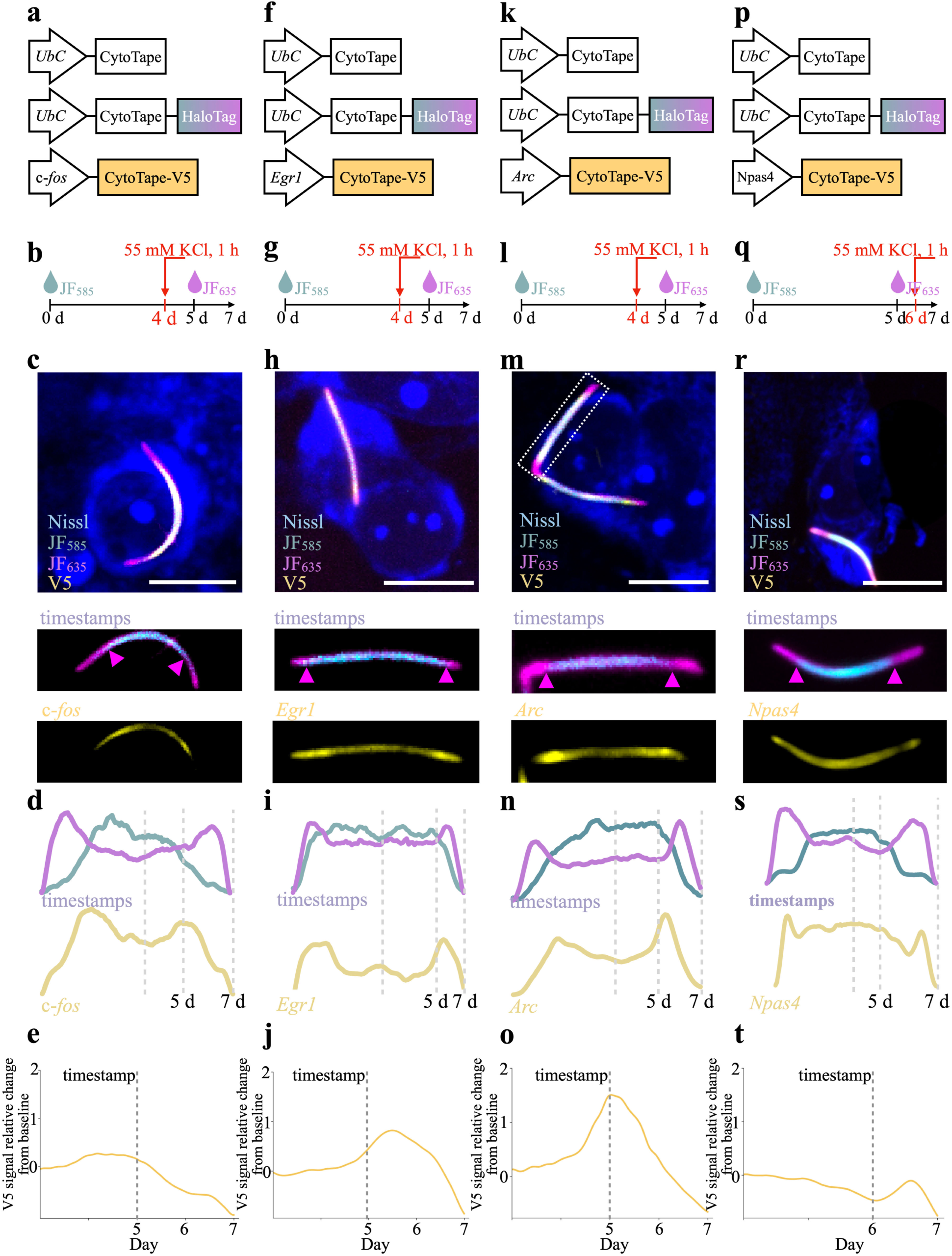
Recording c-*fos*-, *Arc*-, *Egr1*-, and Npas4-promoter driven expression histories in cultured neurons. (**a**), (**f**), (**k**), and (**p**) Schematic of the constructs transfected into cultured mouse hippocampal neurons and (**b**), (**g**), (**l**), and (**q**) the experimental timeline. The JF_585_ and JF_635_ dyes were added to neuron cultures on day 0 and day 5, respectively, and fixation was performed on day 7. Neurons were stimulated with 55 mM KCl for 1 h on day 5 for c-*fos*, *Arc*, *Egr1* and day 6 for Npas4. CytoTape-HA refers to CytoTape fused to the HA tag and CytoTape-V5 refers to CytoTape fused to the V5 tag. (**c**), (**h**), (**m**), and (**r**) Images of cultured neurons expressing the constructs shown in (**a**), (**f**), (**k**), and (**p**) following stimulation with 55 mM KCl for 1 h. Images were captured after fixation, Nissl staining, and immunostaining for HA and V5 tags. Enlarged views of the CytoTape in the top-row panels are shown in the two rows of rectangular panels below. The second-row and third-row panels show the timestamps and activity-dependent-promoter-driven gene expression, respectively. Scale bars, 10 µm. (**e**) c-*fos*, (**j**) *Egr1*, (**o**) *Arc*, and (**t**) Npas4 signal relative change from baseline plotted against recovered time (interpolated with timestamps and fixation time) after calcium phosphate transfection. The dashed gray lines indicate the timestamps, marking the points after which the recovered time is accurate.

**Extended Data Fig. 9.**
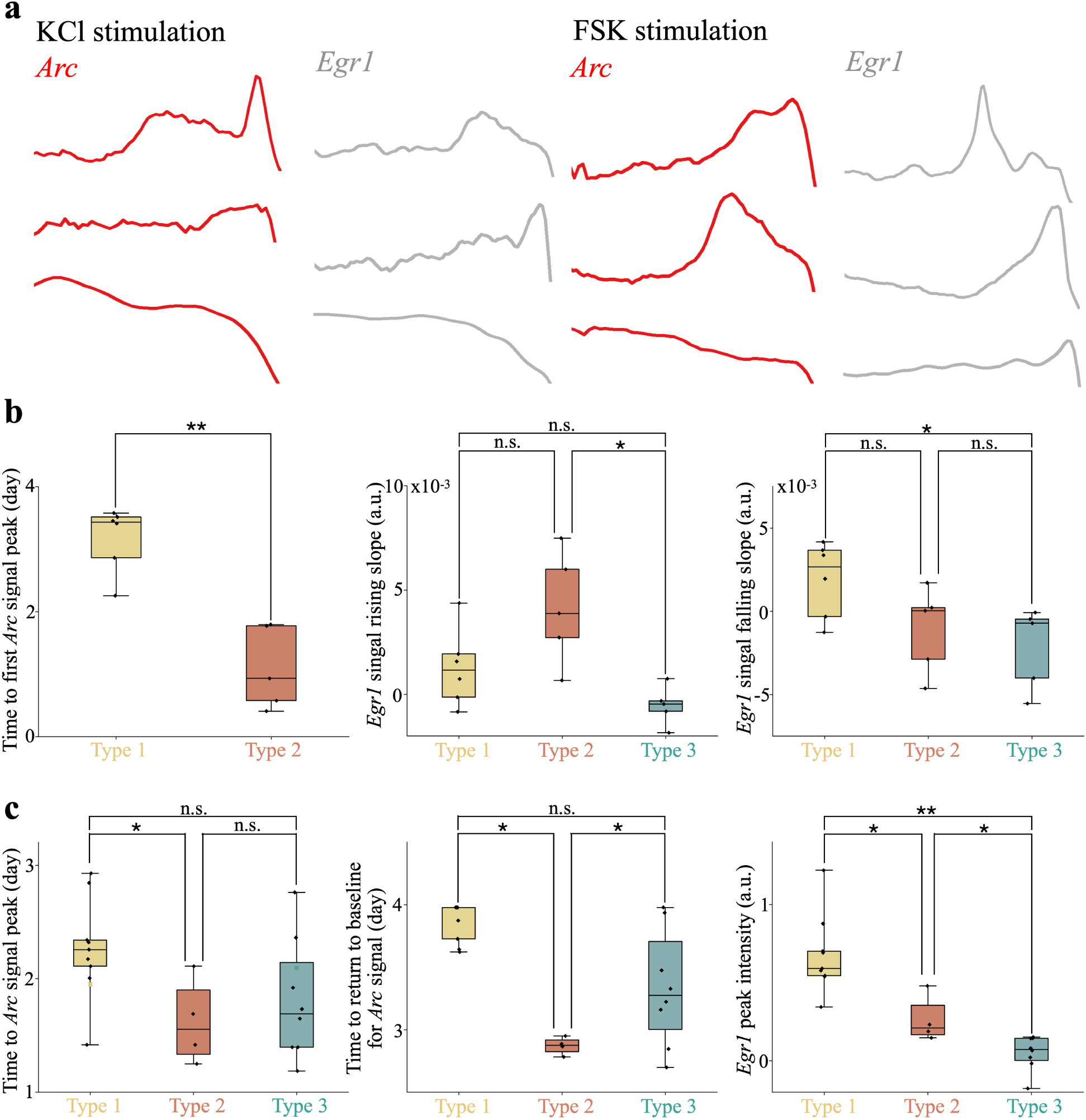
Temporal analysis of multiplexed recording of *Arc*- and *Egr1*-promoter-driven expression histories with CytoTape in cultured neurons. (**a**) Examples of single traces of *Arc* and *Egr1* signals under KCl (left panel) and FSK stimulation (right panel) recorded in separate CytoTapes without multiplexing. (**b**) Scatter plots showing the time between stimulation point (day 17) and the first peak maximum amplitude point of *Arc* signal (left panel, only Type 1 and Type 2 data are shown as Type 3 has no peak), the rising slope of the *Egr1* signal (middle panel), and the falling slope of the *Egr1* signal (right panel) across three neuronal subtypes under KCl stimulation. Data are from n = 16 CytoTapes from 16 neurons from three cultures. (**c**) Scatter plots showing the time between stimulation point (day 17) and the peak maximum amplitude point of *Arc* signal (left panel), the time between stimulation point (day 17) and peaks’ returning to baseline point for *Arc* signal (middle panel), and the peak maximum intensity of the *Egr1* signal (right panel) across three neuronal subtypes under FSK stimulation. Data are from n = 21 CytoTapes from 21 neurons from three cultures. Middle line in box plot, median; box boundary, interquartile range; whiskers, minimum and maximum; black dots, individual data points. *, *P <* 0.05; **, *P <* 0.01; n.s., not significant; Kruskal–Wallis analysis of variance followed by Dunn’s post hoc tests. See **Supplementary Table 2** for details of statistical analysis.

**Extended Data Table 1:** Comparison of XRI, iPAK4, and CytoTape in different cell types.

| **Tech Specs** | **XRI<sup>38</sup>** | **iPAK4<sup>39</sup>** | **CytoTape (this work)** |
| --- | --- | --- | --- |
| Morphology in cultured neurons | fiber | fiber | fiber |
| Morphology in HEK cells | puncta and fiber | fiber | fiber |
| Morphology in HeLa cells | puncta and fiber | not tested | fiber |
| Morphology in cultured glial cells | not tested | not tested | fiber |
| Protruding outward and significantly deforming the cell membrane when the fiber is longer than the cell | not observed | observed | not observed |
| Time recovery method | Time calibration via tamoxifen-induced Cre/FLEX system | Timestamps via HaloTag and dye switches | Timestamps via HaloTag and dye switches |
| Longest recording duration reported in cultured neurons (day) | 7 | 3 | 21 |
| Longest recording duration reported in HEK cells (day) | not tested | 1 | 5 |
| Longest recording duration reported in HeLa cells (day) | not tested | not tested | 4 |
| Number of developed recorders for distinct cell physiological activities | 1 (*c-fos* transcriptional activity) | 1 (*c-fos* transcriptional activity) | 7 (*c-fos*, *Arc*, *Egr1*, and *HSPA1A* transcriptional activities; CREB, Npas4, and Fos protein activities) |
| Reported number of simultaneously recorded distinct cell physiological activities | 1 | 1 | 5 |

**Supplementary Table 1.** Reagents and Resources.

| Reagent or Resource | Source | Identifier |
| --- | --- | --- |
| **Antibodies** |  |  |
| E Tag Monoclonal Antibody (11H12B3) | Invitrogen, Thermofisher Scientific | Catalog # MA5-38276; RRID: AB_2898191 |
| V5 Tag Monoclonal Antibody (SV5-Pk1) | Invitrogen, Thermofisher Scientific | Catalog # R960-25; RRID:AB_2556564 |
| OLLAS Tag Monoclonal Antibody (L2) | Invitrogen, Thermofisher Scientific | Catalog # MA5-16125; RRID: AB_11152481 |
| DYKDDDDK Tag Recombinant Superclonal™ Antibody (20H18L16, 20H1L23, 8H2L5, 8H8L17) | Invitrogen, Thermofisher Scientific | Catalog # 740001;RRID: AB_2610628 |
| HA Tag Polyclonal Antibody | Invitrogen, Thermofisher Scientific | Catalog # PA5-33243; RRID: AB_2550658 |
| HA-Tag (C29F4) Rabbit mAb | Cell Signaling Technology | Catalog # 11846S |
| Mouse IgG1 (Gamma 1 chain) ATTO 425 Conjugated Antibody pre-absorbed | Rockland | N/A |
| Goat anti-Mouse IgG2a Cross-Adsorbed Secondary Antibody, Alexa Fluor™ 488 | Invitrogen, Thermofisher Scientific | Catalog # A-21131; RRID: AB_2535771 |
| Goat anti-Rat IgG (H+L) Cross-Adsorbed Secondary Antibody, Alexa Fluor™ 546 | Invitrogen, Thermofisher Scientific | Catalog # A-11081; RRID: AB_2534125 |
| Goat anti-Rabbit IgG (H+L) Cross-Adsorbed Secondary Antibody, Alexa Fluor™ 594 | Invitrogen, Thermofisher Scientific | Catalog # A-11012; RRID: AB_2534079 |
| Goat anti-Chicken IgY (H+L) Cross-Adsorbed Secondary Antibody, Alexa Fluor™ Plus 647 | Invitrogen, Thermofisher Scientific | Catalog # A32933; RRID: AB_2762845 |
| Goat anti-Mouse IgG2a Cross-Adsorbed Secondary Antibody, Alexa Fluor™ 546 | Invitrogen, Thermofisher Scientific | Catalog # A-21133; RRID: AB_2535772 |
| Goat anti-Rabbit IgG (H+L) Highly Cross-Adsorbed Secondary Antibody, Alexa Fluor™ Plus 488 | Invitrogen, Thermofisher Scientific | Catalog # A-11008; RRID: AB_143165 |
| NeuroTrace™ 435/455 Blue Fluorescent Nissl Stain | Invitrogen, Thermofisher Scientific | Catalog # N21479 |
| **Janelia Fluor® Dyes** |  |  |
| Janelia Fluor® 503 HaloTag® Ligand | Promega | Catalog # HT1010 |
| Janelia Fluor® 585 HaloTag® Ligand | Promega | Catalog # HT1040 |
| Janelia Fluor® 635 HaloTag® Ligand | Promega | Catalog # HT1050 |
| **Chemicals, Peptides, and Recombinant Proteins** |  |  |
| MAXBlock Blocking medium | Active Motif | Cat # 15252 |
| MAXWash Washing medium | Active Motif | Cat # 15254 |
| MAXStain Staining medium | Active Motif | Cat # 15253 |
| Formamide | Millipore Sigma | Cat # F9037 |
| Glycine | Millipore Sigma | Cat#G5417 |
| Forskolin | Millipore Sigma | Cat#F6886 |
| AraC | Millipore Sigma | Cat#C6645 |
| HEPES | Millipore Sigma | Cat#H3375 |
| Minimum Essential Media, no glutamine, no phenol | Life Technologies | Cat#51200-038 |
| Matrigel | Corning, VWR | Cat#47743-715 |
| Heat inactivated fetal bovine serum | Corning, VWR | Cat#45000-736 |
| anhydrous DMSO | ThermoFisher Scientific | Cat#D12345 |
| **Critical Commercial Assays** |  |  |
| Plasmid mini-prep kit | QIAGEN | Cat#27104 |
| TransIT-X2 Dynamic Delivery System kit | Mirus Bio, VWR | Cat#10766-888 |
| Calcium phosphate transfection kit | Invitrogen, ThermoFisher Scientific | Cat#K278001 |
| **Experimental Models: Cell Lines** |  |  |
| HeLa | ATCC | CCL-2 |
| HEK293T | ATCC | CRL-11268 |
| **Experimental Models: Organisms/Strains** |  |  |
| Swiss Webster mice, pregnant female | Taconic | N/A |
| **Software and Algorithms** |  |  |
| ProtSSN | Github | https://github.com/tyang816/ProtSSN |
| ImageJ | NIH | https://imagej.nih.gov/ij/download.html |
| Fiji software GPL v2 | Fiji | https://fiji.sc/Fiji |
| Origin | OriginLab | https://www.originlab.com/ |
| Prism | GraphPad | https://www.graphpad.com/ |

| Other |  |  |
| --- | --- | --- |
| White Mounted LED (400 - 700 nm Wavelength Range) 4000 K, 1400 mW (Min) Mounted LED, 2000 mA | Thorlabs | MNWHLP1 |
| Adjustable Collimation Adapter with Ø2" Lens, AR Coating: 350 - 700 nm | Thorlabs | N/A |
| Microscope Collimation Adapters with Ø50 mm Lens Adapter with External SM1 Threads and Internal SM2 Threads | Thorlabs | N/A |
| LED Driver, 4.0 A Max, 14.0 V Max | Thorlabs | DC40 |

**Supplementary Table 2.** Statistical Analysis. (See Supplementary Table 2 in a separate Excel file in Supplementary Material at bioRxiv.)

## Supplementary information

### Supplementary Tables

**Supplementary Table S1.**
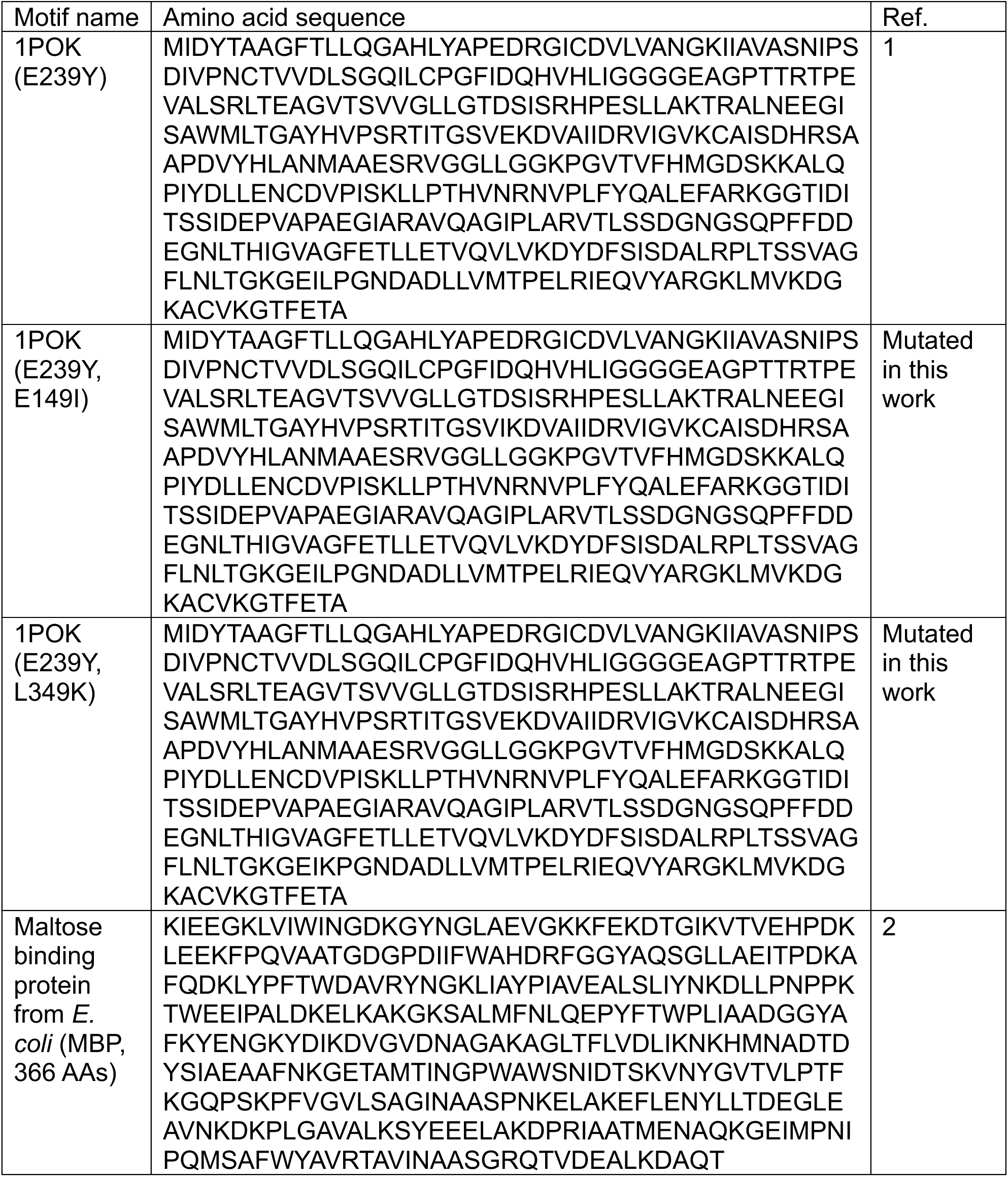
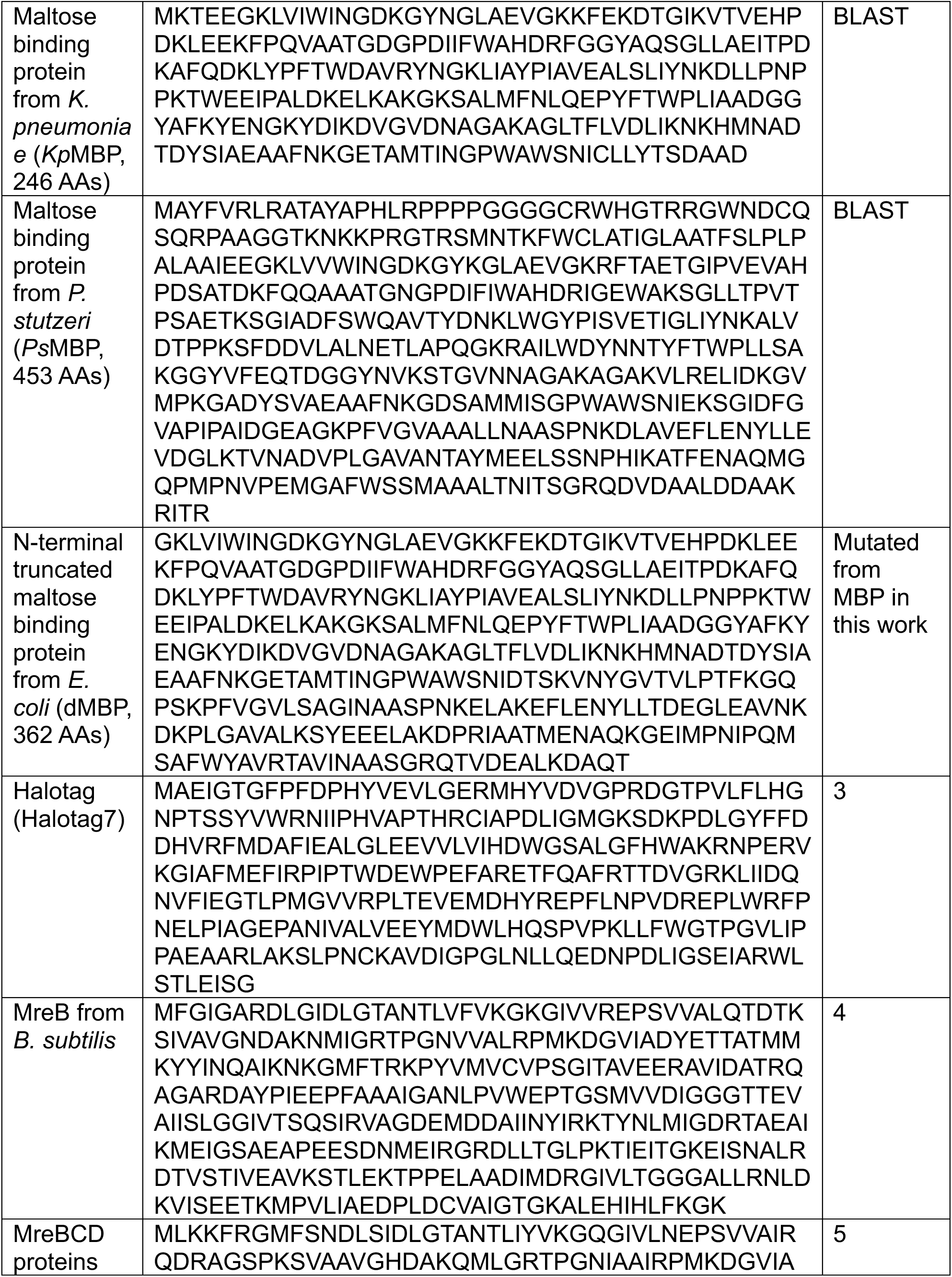
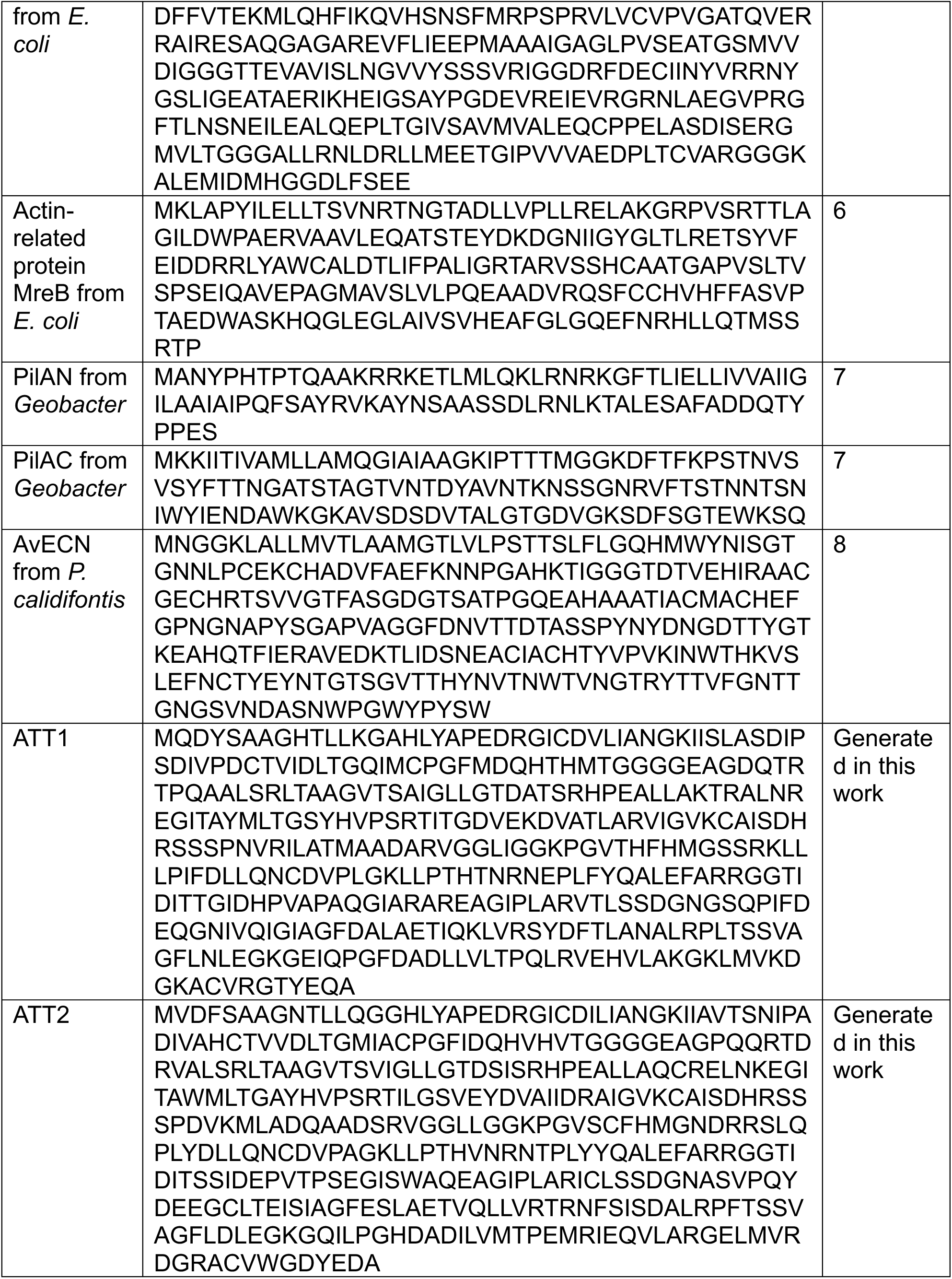
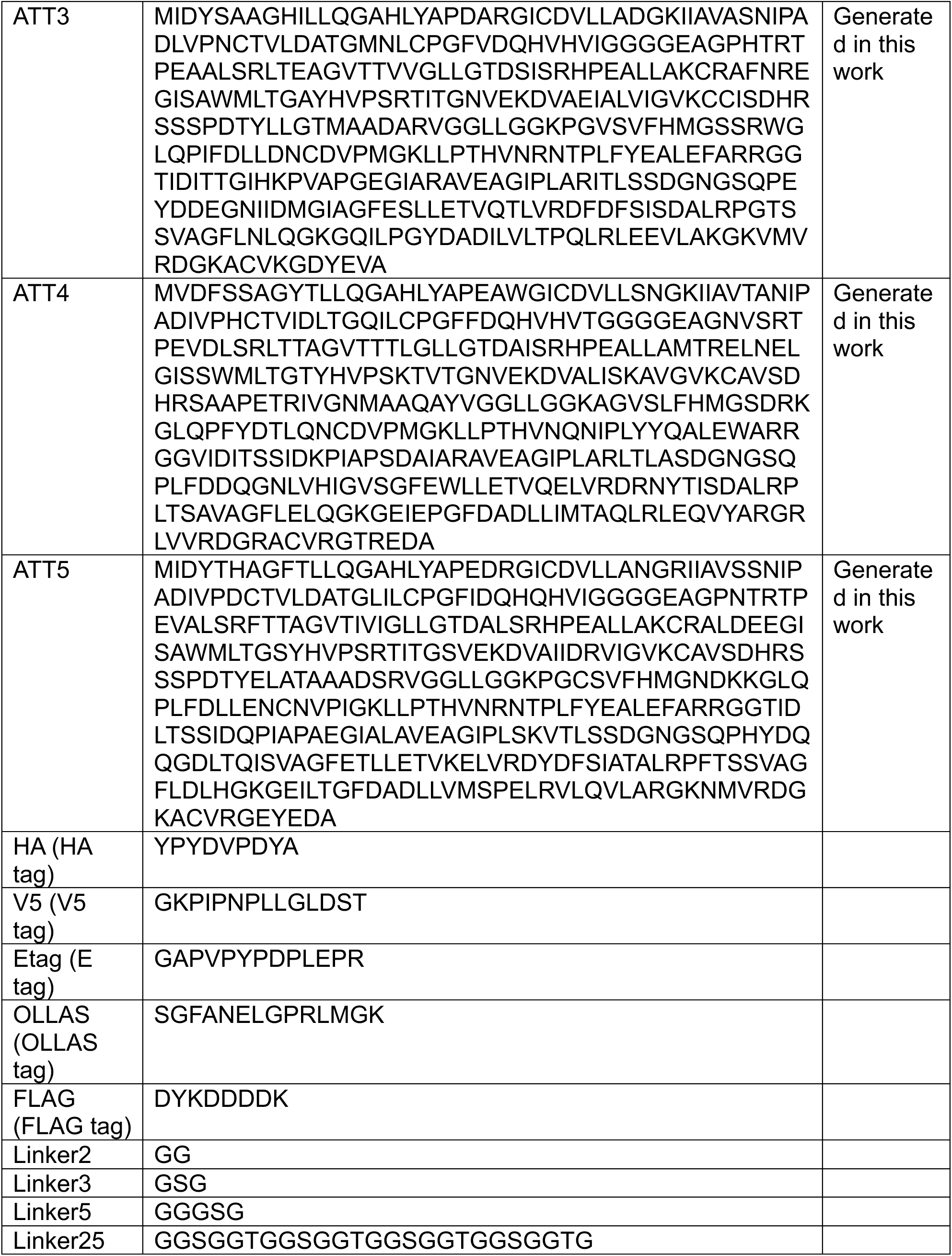
Sequences of protein motifs used in this study.

**Supplementary Table S2.** Constructs of structural monomer tested in cultured neurons in this study.

| **Name** | **Construct** (promoters are underlined) | **Resulted pattern of protein self-assembly** (in the cytosol unless noted otherwise) |
| --- | --- | --- |
| A1-1 | <u>UbC</u>-ATT1-Liner25-HA-Linker3-MBP | Fibers |
| A1-2 | <u>UbC</u>-ATT2-Liner25-HA-Linker3-MBP | Fibers |
| A1-3 | <u>UbC</u>-ATT3-Liner25-HA-Linker3-MBP | Fibers |
| A1-4 | <u>UbC</u>-ATT4-Liner25-HA-Linker3-MBP | Short fibers |
| A1-5 | <u>UbC</u>-ATT5-Liner25-HA-Linker3-MBP | Fibers |
| A2-1 | <u>UbC</u>-1POK(E239Y, E149I)-Linker25-HA-Linker3-MBP | Fibers |
| A2-2 | <u>UbC</u>-1POK(E239Y, L349K)-Linker25-HA-Linker3-MBP | Fibers |
| A3-1 | <u>UbC</u>-mhYFP-Linker5-B. subtilis MreB(mrebbs) | Puncta |
| A3-2 | <u>UbC</u>-mhYFP-Linker5-E. coli(mrebec) | Intertwined fibers |
| A3-3 | <u>UbC</u>-mGFP-Linker5-EcMReB | Intertwined fibers |
| A3-4 | <u>UbC</u>-PilAN-Linker3-HA-Linker3-PilAC | Puncta |
| A3-5 | <u>UbC</u>-AvECN-Linker3-HA | Puncta |
| B-1 | <u>UbC</u>-1POK(E239Y)-Linker2-MBP-HA | Fibers |
| C-1 | <u>UbC</u>-1POK(E239Y)-Liner25-HA-Linker3-*Kp*MBP | Fibers |
| C-2 | <u>UbC</u>-1POK(E239Y)-Liner25-HA-Linker3-*Ps*MBP | Puncta |
| C-3 | <u>UbC</u>-1POK-Liner25-HA-Linker3-MBP-Linker3-MBP | Puncta |
| C-4 | <u>UbC</u>-1POK(E239Y)-Linker2-dMBP-HA | Fibers |
| C-5 | <u>UbC</u>- mGFP-P2A-1POK(E239Y)-Liner25-HA-Linker3-MBP | Fibers |
| CytoTape | <u>UbC</u>-1POK(E239Y, L349K)-dMBP-HA | Fibers |

**Supplementary Table S3.** Constructs of signal monomer tested in neurons in this study.

| **Construct** (promoters are underlined) | Ref. for promoter sequence |
| --- | --- |
| *c-fos*-1POK(E239Y, L349K)-dMBP-V5 | 9 |
| <u>SARE</u> <u>ArcMin</u>-1POK(E239Y, L349K)-dMBP-V5 | 10 |
| <u>6xCRE</u> <u>CMVMin</u>-1POK(E239Y, L349K)-dMBP-V5 | 11 |
| <u>Egr1</u>-1POK(E239Y, L349K)-dMBP-V5 | 12 |
| <u>F-RAM</u>-1POK(E239Y, L349K)-dMBP-V5 | 13 |
| <u>N-RAM</u>-1POK(E239Y, L349K)-dMBP-V5 | 13 |
| <u>Egr1</u>-1POK(E239Y, L349K)-dMBP-OLLAS | 12 |
| <u>SARE</u> <u>ArcMin</u>-1POK(E239Y, L349K)-dMBP-OLLAS | 10 |

**Supplementary Table S4.**
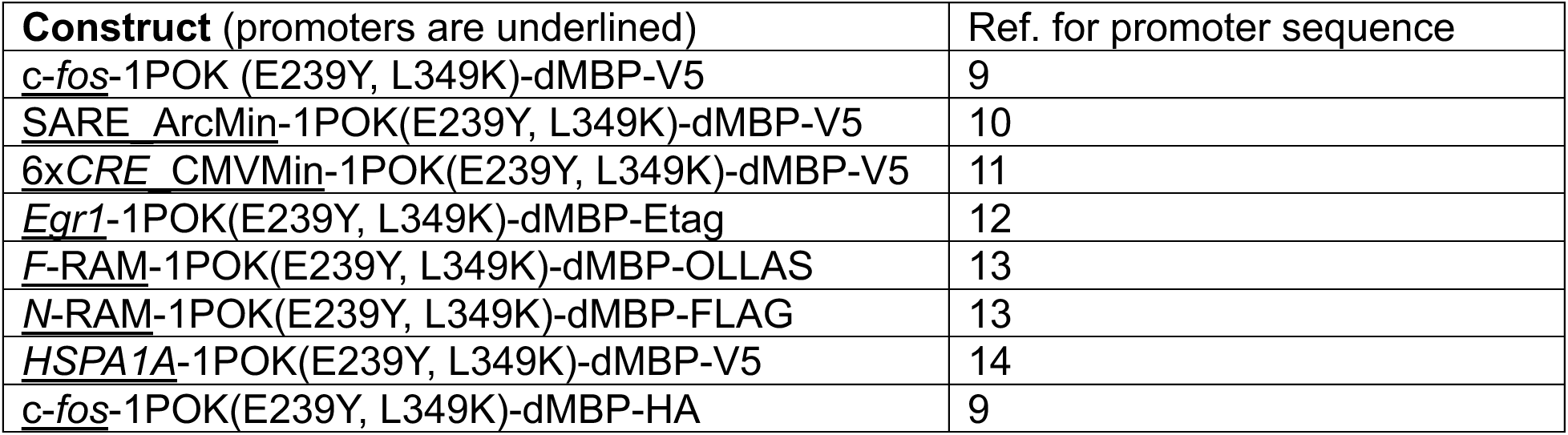
Constructs of signal monomer tested in HEK293T in this study.

**Supplementary Table S5.** Constructs of signal monomer tested in HeLa in this study.

| **Construct** (promoters are underlined) | Ref. for promoter sequence |
| --- | --- |
| <u>HSPA1A</u>-1POK(E239Y, L349K)-dMBP-V5 | 14 |

**Supplementary Table S6.** Constructs of timestamp monomer.

| **Construct** (promoters are underlined) | Ref. for HaloTag sequence |
| --- | --- |
| <u>UbC</u>-1POK(E239Y, L349K)-dMBP-HA-gsg-HaloTag | 3 |

